# The Coronavirus Envelope Is Modulated by Host Inflammation and Enriched in Bioactive Lipids Required for Replication

**DOI:** 10.1101/2025.11.03.686211

**Authors:** Victoria J Tyrrell, Andreas Zaragkoulias, Wendy Powell, Miran Y Elfar, Valeria Grossoni, Federica Monaco, Majd B Protty, Ewan H Stenhouse, James Heyman, Daniela Costa, Lisa Roche, Ceri-Ann Brigid Lynch, Peter Ghazal, Simon A Jones, Gérard Lambeau, Yvonne Benatzy, Joshua D Kandler, Denisa Bojkova, Ryan G Snodgrass, Bernhard Brüne, David W Thomas, Richard J Stanton, Valerie B O’Donnell

## Abstract

How inflammation or disease regulates coronavirus lipid membranes is currently unknown, while patient-derived viral envelopes have never been structurally characterized. Here, we show that four cultured SARS-CoV-2 strains (England2, Alpha, Beta, and Delta) possess conserved, phospholipid- and cholesterol-rich envelopes, with pro-thrombotic and infection-promoting aminophospholipids (aPL) displayed predominantly on the outer leaflet (approximately 70–80%). Exposure to interleukin-4 (IL-4) markedly altered envelope fatty acyl composition, whereas interleukin-6 (with or without its soluble receptor IL-6Rα) and dexamethasone had no detectable effect. Viral envelopes were susceptible to hydrolysis by secretory phospholipase A2 (sPLA2), an enzyme associated with adverse clinical outcomes. SARS-CoV-2 isolated directly from patient saliva exhibited cholesterol-enriched envelopes that were highly conserved across clinical isolates. In addition, clinical samples contained pro-coagulant oxidized phospholipids and bioactive lipoxygenase (LOX)-derived oxylipins. The dominance of external facing pro-coagulant aPL and eoxPL may support known thrombotic complications of severe COVID19 viremia. Last, gene-silencing experiments demonstrated that 15-LOX2 is required for replication of related coronaviruses. Together, these findings reposition the coronavirus envelope as an active, dynamic structure rather than a passive scaffold, and challenge the protein-centric view of viral function. The lipid envelope is proposed as a potential therapeutic target through modulation of host innate immunity, and dampening thrombotic potential.

**Significance statement:** Viruses such as SARS-CoV-2 are surrounded by a host-derived lipid envelope. Little is known about how this changes during infection/inflammation. We determined the lipid composition of the SARS-CoV-2 envelope using both laboratory-grown viruses and patient isolates. Across several pandemic strains, the envelope was rich in cholesterol and phospholipids and showed a consistent structure. Lipids linked to thrombosis and infection were mainly exposed on the outer virus surface. The inflammatory cytokine interleukin-4 altered the envelope’s fatty acid composition, while other treatments did not. Patient-derived viruses contained additional bioactive lipids, and blocking an enzyme that generates these lipids reduced coronavirus replication. In summary, the envelope is an active component of infection and potential target for new treatments to dampen infectivity and thrombosis.

## Introduction

Lipid envelopes are an essential structural and functional component of multiple virus families, including coronaviruses such as severe acute respiratory syndrome coronavirus 2 (SARS-CoV-2) and Middle East respiratory syndrome (MERS), as well as influenza, HIV, and herpes simplex virus(*1,2*). They act as a carrier for viral proteins and are thought to influence virion stability. They represent a direct target for anti-viral disruption and could potentially host bioactive or pro-coagulant lipids generated by the host, although this has not been examined. It is generally assumed that envelope composition reflects the membrane from which they bud. In this regard, the envelope of a laboratory-grown SARS-CoV-2 strain (England2) was shown in our recent study to be phospholipid (PL)-rich, with a cholesterol content consistent with its trafficking through lysosomes prior to shedding(*3*). It lacked asymmetry, with high levels of pro-coagulant phosphatidylserine (PS) and phosphatidylethanolamine (PE) on its outer surface(*3*). However, beyond this, many fundamental questions remain. For example, whether altered envelope lipid composition has contributed to the evolution of SARS-CoV-2 towards variants with increased pandemic potential has not been determined. Whether virion membrane lipids adapt to reflect host lipid metabolism in response to inflammatory mediators released during infection is unknown for this or indeed any enveloped virus, nor is it understood which lipids support viral replication. Furthermore, to date, viral envelopes have only been analyzed from in vitro-grown virus. How this relates to the composition of in vivo replicated virus is completely unknown.

Here, we address these fundamental questions using lipidomic and genomic approaches applied to SARS-CoV-2, MERS-CoV, and HCoV-229E, examining both viruses grown in vitro and viruses isolated directly from infected human patients. Our study provides new insights into the biology of the viral envelope, specifically: (i) variation in lipid molecular species and externalization of pro-coagulant phospholipids across multiple SARS-CoV-2 pandemic strains; (ii) modulation of envelope composition in response to inflammatory stimuli, including interleukin-6, interleukin-4, and dexamethasone; (iii) susceptibility of viral envelopes to hydrolysis by secreted phospholipases; (iv) detailed lipid profiles of patient-derived viruses, including major lipid classes, oxidized phospholipid damage-associated molecular patterns (DAMPs), and lipoxygenase-derived oxylipins; and (v) evidence that lipoxygenases are required for coronavirus replication in vitro.

Our findings provide fundamental new insights into the biology of coronavirus lipid envelopes, with potential relevance to disease pathogenesis and broader implications for understanding envelope biology in other human pathogens, including influenza virus, human immunodeficiency virus (HIV), herpes simplex virus, Zika virus, and others.

## Results

### SARS-COV-2 variants of concern (VOC) show conserved envelope lipid composition

Coronaviruses, including SARS-CoV-2, bud from the endoplasmic reticulum (ER)/Golgi intermediate complex and exit cells via lysosomes, all of which are organelles surrounded by a phospholipid bilayer, and relatively low in neutral lipids such as glycerides and sterol esters(*4–9*). There is also evidence for interplay between several enveloped viruses and lipid droplets, glyceride and sterol ester-rich organelles surrounded by a phospholipid (PL) monolayer associated with ER, mitochondria, peroxisomes, and endosomes(*10–13*). In the case of SARS-CoV-2, lipid droplets may directly act as an assembly platform for the virus(*14*). Considering this, the virus lipid envelope composition may be strongly influenced by its interactions with intracellular membranes. Furthermore, as SARS-CoV-2 evolved during the pandemic to generate new variants containing genetic alterations in multiple viral proteins, we hypothesized that virions with altered infectivity may show differences in envelope composition.

To test this, the envelopes of four variants of SARS-CoV-2 cultured in AAT cells were compared using targeted lipidomics. To standardize membrane composition between strains and biological replicates, lipid composition is expressed as relative abundance, enabling comparison within and across individual preparations. Across lipid categories, envelopes were remarkably similar between strains, with few significant differences detected (Figure 1 A,B, Supplementary Figure 1). Although the Beta strain appeared to contain more PE and PG, in tandem with less PC and PS, this was driven by high variability between biological isolates, and differences were generally not significant. Comparing individual molecular species also showed high similarity for most species (Supplementary Figure 2). For PEs, several low-abundance species were lower in the Beta strain, but this was driven by the higher (non-significant) levels of PE 16:0_16:0 and PE 16:0_16:1. Overall, the composition of the virus envelope for these 4 variants are remarkably similar. The ratio of cholesterol:phospholipids was also relatively similar across all four strains (Figure 1 C).

**Figure 1.**
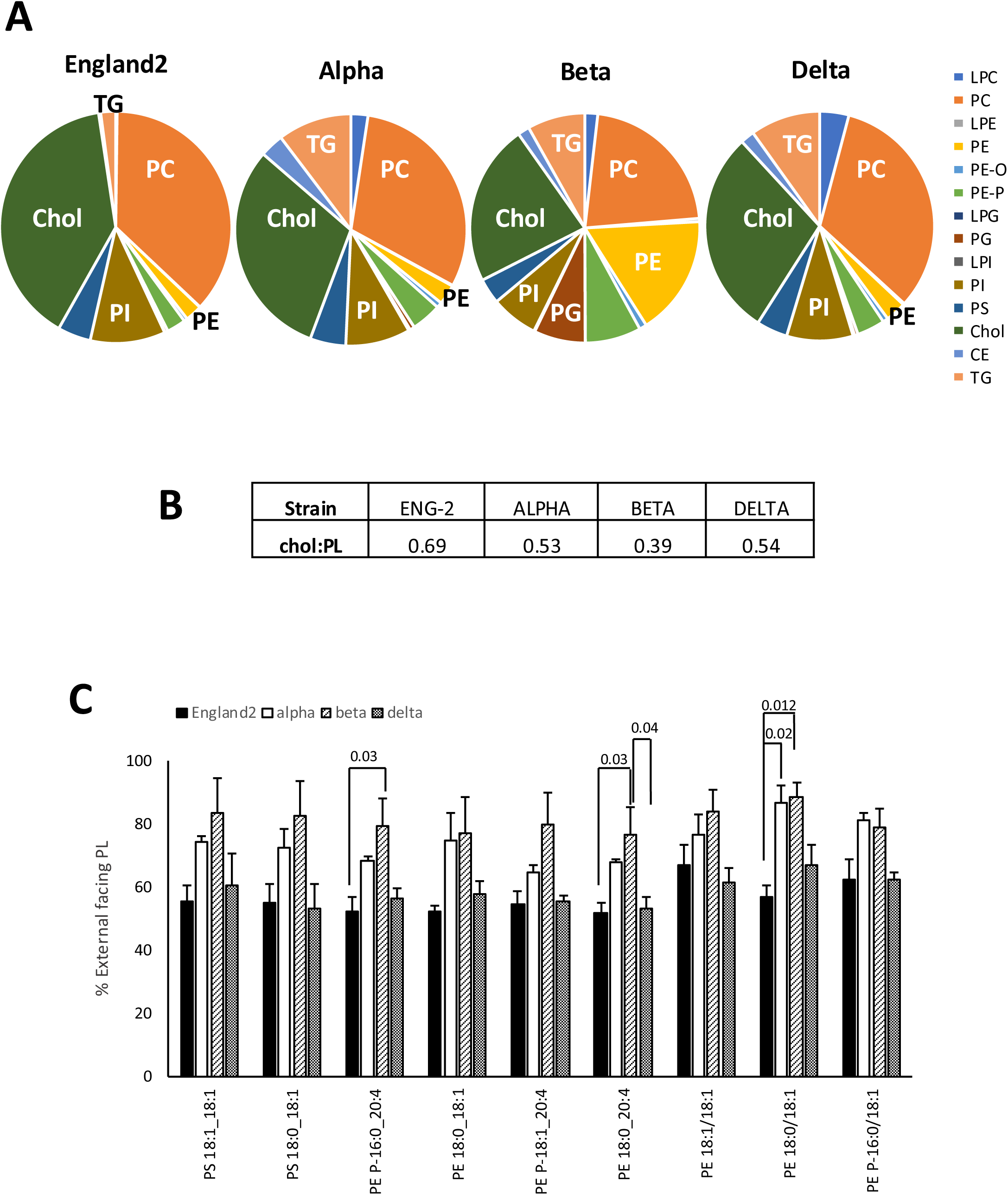
Lipidomics of four SARS-CoV-2 pandemic strain viruses reveals them to be relatively conserved with high externalization of PE and PS on the outer surface. *Panel A. Relative abundance of total lipid categories for four SARS-CoV-2 pandemic strains is similar.* Lab grown virus from AAT cells was isolated using density gradient centrifugation, lipids extracted and then analyzed using targeted LC/MS/MS as outlined in Methods. Each isolate was from a different culture obtained at least one week apart. Alpha, Beta, Delta (n=3 isolates mean +/- SEM), England2 (n=2 isolates, mean +/- SD). Relative abundance was calculated from ng values of all lipid species totaled, converted to molar amount, then related to a mean mass value for each category, and shown as a pie chart. *Panel B. Cholesterol:phospholipid ratio across the pandemic strains.* The ratio was calculated using molar amounts of cholesterol or total of all PL categories. *Panel C. The total PE or PS facing the outside of the viral envelope is extremely high, especially for alpha and beta strains*. External and total PE and PS was determined using LC/MS/MS as outlined in Methods for the four pandemic strains (n=3 isolates mean +/- SEM), one way ANOVA with Tukey Post Hoc test.

### High levels of phosphatidylserine (PS) and phosphatidylethanolamine (PE) are present on the outer surface of the lipid envelope for all variants

PE and PS on the outside of blood cell membranes are strongly pro-coagulant(*15*), while PS appears to be involved in viral entry(*16*). We previously showed using lipidomics that the England2 strain externalizes around 50 % of its PE and PS on the outer surface and that pathophysiological levels of virus can stimulate plasmatic clotting *in vitro*(*3*). Extending this analysis to the other 3 variants, while Delta was similar, Alpha and Beta strains frequently displayed around 70 – 80% of their PE and PS on the outer leaflet (Figure 1 C). Thus, other variants will be equally capable, if not more, of supporting PE- and PS-dependent functions of the viral lipid envelope, including infectivity and coagulation, potentially contributing to elevated thrombotic risk during disease.

### Inflammation has a significant and selective impact on virus envelope lipid composition

During viral infection and ensuing inflammation, the lipid composition of mammalian cell membranes is significantly impacted. Along with this, replication of enveloped viruses is associated with major changes in host lipid metabolism, required to support the biogenesis of new virion membranes(*17*). As one example, Hepatitis C virus takes over the liver pathway for VLDL assembly, maturation, and secretion to generate infectious virions(*17–19*). In the case of SARS-CoV-2, infected A549 cells show increases in triacylglycerides (TAGs), ceramides, and PL, while infected Caco-2 cells conversely downregulate metabolism of ceramides, glycerolipids, and ether lipids(*20*). Mapped onto this, inflammation directly impacts lipid metabolism, with phospholipases activated to cleave fatty acyls (FA) from PL, generating bioactive lysophospholipids (LysoPL). Additionally, FA are oxygenated to prostaglandins and other oxylipins during viral infection, including in COVID19(*21*). PL remodeling via the Lands cycle is also modulated during inflammation, with some lysoPL acyl transferases (LPATs) upregulated in human disease, such as non-alcoholic fatty liver disease and atherosclerosis (reviewed in(*22*)). Considering this, we hypothesized that virion envelope composition may be impacted by membrane dynamics of host cell organelles when the virus is replicating.

Three factors, which are either implicated in the pathogenesis of COVID19; interleukin-6 (IL-6), interleukin-4 (IL-4), or used for treatment (dexamethasone), that can all impact host lipid metabolism were tested. IL-6 is significantly increased in many viral diseases and in COVID19, is associated with disease severity, correlates with viral load, and contributes to complications such as acute respiratory distress syndrome (ARDS)(*23–29*). IL-4 is a Th2 cytokine which is elevated in and associated with severity and death in COVID19(*23,27–30*). Furthermore, severe asthma, where Th2 cytokines are elevated has been associated with worse outcomes(*31,32*). Activation of the STAT6 pathway by either IL-4 or IL-13 leads to generation of pro-coagulant oxidized phospholipids, including in primary human airway epithelia(*33,34*). Dexamethasone has been a recommended treatment for severe COVID19 since it was shown to result in lower mortality among patients on invasive ventilation or oxygen therapy(*35*). A mainstay treatment for COVID, Dex is administered intravenously or orally to patients in hospital during the inflammatory stage of disease(*36,37*) and also administered long-term by inhalation to asthmatics as a preventative drug. It is an agonist of the glucocorticoid receptor, that upregulates anti-inflammatory responses, while downregulating pro-inflammatory processes. It has multiple and complex effects on lipid metabolism in particular in the liver.

#### (i) IL-4 increases shedding of particles with dramatically alters phospholipid composition

AAT cells were pre-treated with IL-4 for 5 days before virus infection, as for induction of the Th2 phenotype in macrophages(*38*). Furthermore, IL-4 will already be present in asthmatics prior to COVID infection *in vivo*. Virus particles were harvested and purified after a further 72-hour culture. Lipidomics demonstrated that total amount of lipid detected was increased dramatically in response to IL-4 (Figure 2 A, Supplementary Figure 3 A). However, IL-4 had only a small impact on relative lipid class composition (Figure 2 B, Supplementary Figure 3 B,C). Specifically, while there were significant decreases for total PE ethers and PS species, other changes were not significant (Supplementary Figure 3 C). Next, the impact of IL-4 on fatty acyl (FA) composition of complex lipids was tested. Across all PL categories, particles isolated from IL-4 conditioned cells showed a consistent change in FA where 18:2/18:3-containing species were significantly enriched, while longer chain PUFA-containing PL (20:4, 20:5, 22:4, 22:5, 22:6) were depleted (Figure 2 C,D, Supplementary Figure 4). Also, PL containing 16:0, 18:0 and 18:1 at both *Sn1* and *Sn2* positions were significantly depleted (Figure 2 C,D, Supplementary Figure 4). Thus, IL-4 is causing significant changes to FA composition of shed particles as well as to the amount of particles released.

**Figure 2.**
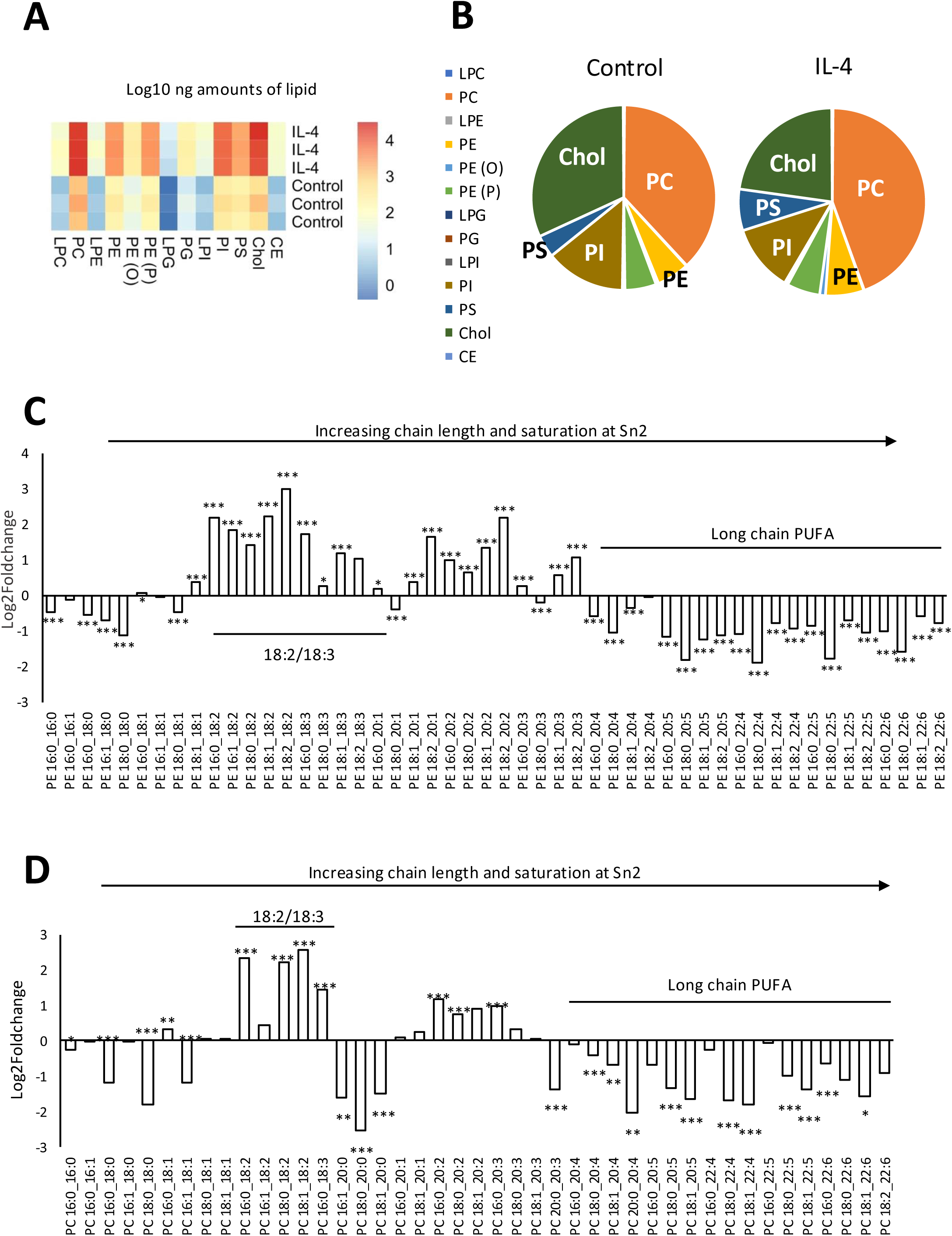
**IL-4 pretreatment of host AAT cells leads to increased shedding of particles with significantly altered FA, in particular reduced PUFA and saturated FA containing PE, but increased PE with 18:2/18:3**. *Panel A. Total virus lipid recovered is significantly increased from IL-4 treated host cells.* AAT cells were pretreated with IL-4 for 5 days as outlined in Methods, before infection with SARS-CoV-2 England2 strain. Virus was harvested using density gradient centrifugation and lipids extracted and quantified using LC/MS/MS as outlined in methods. Heatmap shows log10 ng values as total for each PL category, for 3 independent virus isolates. *Panel B. Lipid composition of the particles is not particularly changed by IL-4 treatment.* Data from Panel A was instead expressed as relative molar proportions within samples, by normalizing to an average mass value per category and shown in a pie chart (n=3 isolates mean +/- SEM). *Panels C,D. IL-4 treatment leads to significant remodeling of FA across PE species*. Amounts of individual PE or PC species detected were averaged and compared for an impact of IL-4 treatment, and are shown as log2foldchange. A decrease means reduction in the species by IL-4, (n=3 isolates, mean +/- SEM), unpaired Student t-test, * P<0.05, ** P<0.01, ***P<0.005.

#### (ii) IL-6 signaling does not impact viral envelope composition

To test whether inflammation alters the envelope composition, cells were first treated with the pro-inflammatory cytokine IL-6. Before testing the effect of IL-6, it was first established that while AAT cells express the gp130 subunit of the IL-6 receptor complex, they express little or no IL-6Rα (Figure 3 A,B), in agreement with a previous report(*39*). Thus, IL-6 requires soluble IL-6R (sIL-6Rα) to allow trans-signaling in these cells. AAT cells were infected at low MOI (0.05) with SARS-CoV-2 England2, then after 1 hr incubation, IL-6, with or without sIL-6Rα was added and cultured for a total of 72 hrs, with virus then isolated from supernatant as described in Methods. No difference in levels of recovered virus was found when measured as particle counts (Figure 3 C). Lipidomics also showed that IL-6 alone or IL-6/sIL-6Rα had no significant impact on relative proportions of lipid species in viral envelopes (Figure 3 D, Supplementary Figure 5 A,B). The ratio of cholesterol:phospholipid was also not strongly changed, apart from a small decrease following either treatment (Figure 3 E). The FA composition of individual lipid categories also showed only small impacts at the molecular level (Supplementary Figure 5 C-V). Last, when comparing ng yields of lipids for all categories, no differences were noted (Supplementary Figure 5 W). In summary, IL-6 signaling had no impact on the lipid envelope of SARS-CoV-2 or viral replication.

**Figure 3.**
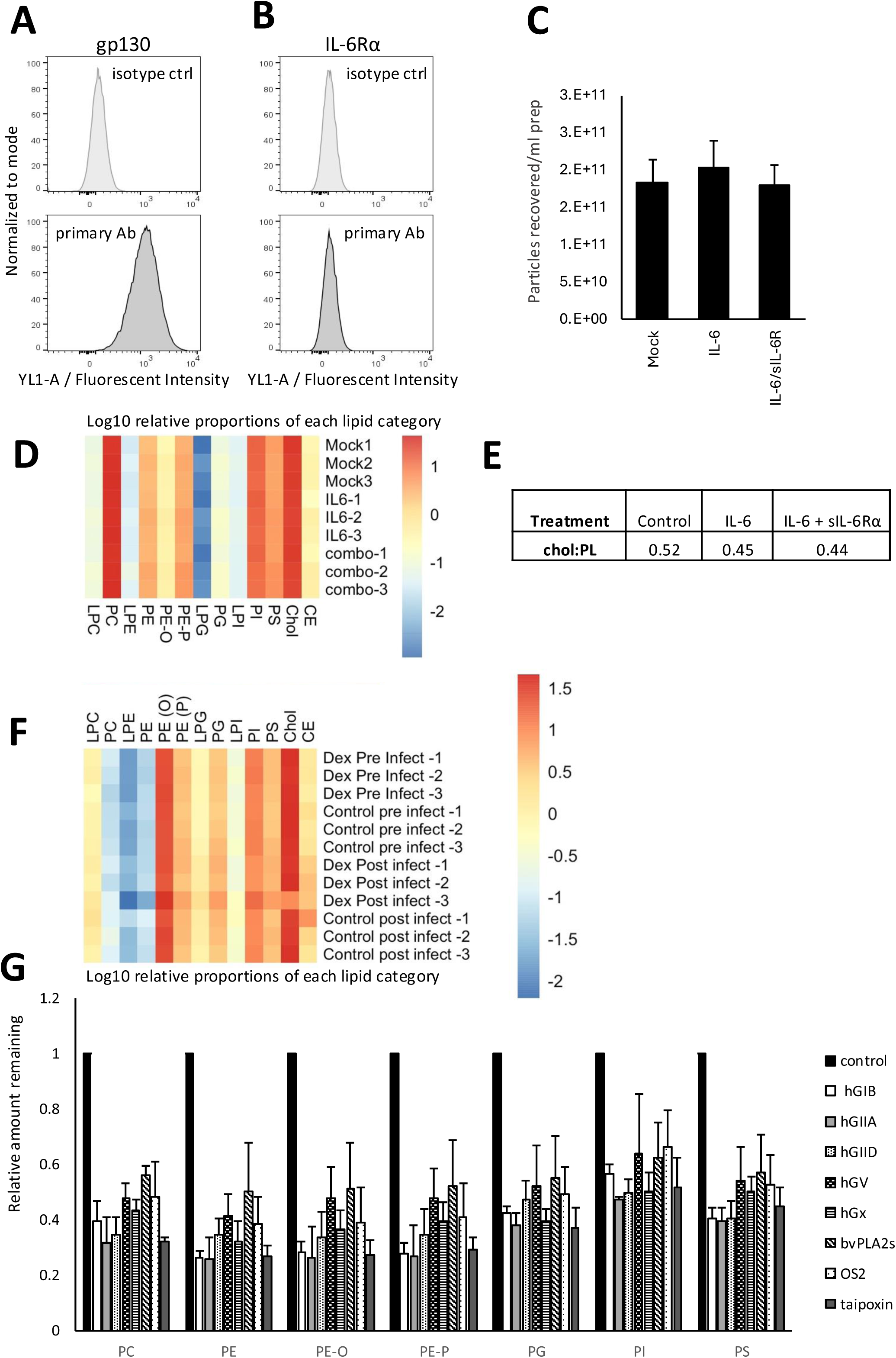
IL-6 or dexamethasone treatment of host cells has no impact on SARS-CoV-2 membrane composition, while the virus is sensitive to sPLA2 hydrolysis by several isoforms. *Panels A,B. AAT cells express gp130 but not IL-6Rα.* AAT cells were harvested and stained with anti-human gp130, anti-human IL-6Rα or respective isotype controls, as outlined in methods before secondary detection using Alexa 488 using flow cytometry. *Panel C. IL-6 signaling had no impact on particles recovered from AAT cells.* Virus was harvested and particles counted as outlined in Methods (n=3 isolates, mean +/- SEM). *Panel D. IL-6 signaling has no impact on lipid composition*. Lipids were extracted from virus from IL-6 or IL-6/sIL-6Rα-treated AAT cells then analyzed using LC/MS/MS as outlined in methods. Lipid amounts are expressed as log10 of relative molar%, by normalizing to an average mass value per category and shown in a heatmap. *Panel E. IL-6 signaling doesn’t impact cholesterol:PL ratio of virus envelopes*. The ratio was calculated using molar amounts of cholesterol or total of all PL categories. *Panel F. Dexamethasone treatment of AAT cells either pre- or during (post-) infection has no impact on lipid composition*. Lipids were extracted from virus from AAT cells treated with dexamethasone, then analyzed using LC/MS/MS as outlined in methods. Lipid amounts are expressed as log10 of relative molar%, by normalizing to an average mass value per category and shown in a heatmap. *Panel G. Several sPLA2 isoforms efficiently hydrolyze SARS-CoV-2 PL.* Virus was incubated with sPLA2 isoforms as indicated in Methods, then lipids extracted and analyzed using LC/MS/MS. The relative amounts of each molecular species remaining was determined, and totaled per lipid category, then normalized to untreated control virus PL levels (n=3 isolates, mean +/- SEM).

#### (iii) Dexamethasone does not impact viral envelope lipid composition

To mimic administration of Dex either prior to or during infection, AAT cells were incubated with the drug (100 nM) either 24 hr before, or 24 hr after inoculation of cells with SARS-CoV-2-England2. There were very minor changes in relative abundance of PL categories between groups following this treatment (Supplementary Figure 6 A). Furthermore, total lipid amounts recovered from individual isolates was relatively similar (Supplementary Figure 6 B). Comparing levels of individual molecular species showed that Dex did not lead to any dramatic changes, and there was no obvious impact of treatment on FA composition (Supplementary Figure 6 C-V). Although some molecular species achieved significance, there were no obvious trends, and differences were low, reducing biological relevance. Overall, Dex did not appear to have a major impact on virus envelope composition.

#### (iv) Secretory phospholipase A2 (sPLA2) enzymes hydrolyze the SARS-CoV-2 envelope

The upregulation and activation of phospholipases is a common feature of inflammation. During COVID19, a role for sPLA2 isoforms was indicated since they increased significantly and correlated with disease severity(*21,40–44*). Furthermore, sPLA2s have an impact on HIV replication(*43*). The hydrolysis of virion envelopes by sPLA2s could have several consequences, including inactivation or enhancement of infectivity, or generation of bioactive lipid mediators. To test whether virions could be hydrolyzed by sPLA2, the hydrolytic activity of several human and non-mammalian isoforms towards lipid membrane composition was tested using cultured virus. Virus was incubated with enzymes for 6 hrs at 37 °C, then lipids extracted and analyzed. Overall, all isoforms tested hydrolyzed around half of the envelope PL, without a preference for any particular headgroup or fatty acyl composition (Figure 3 G, Supplementary Figure 7).

### Lipid composition of human saliva virus isolates is conserved across patients, but highly enriched in triglycerides compared to laboratory-grown virus

To date, the lipid composition of virions has only ever been determined following *in vitro* growth. We therefore determined the lipid composition of *in vivo* cultured SARS-CoV-2 obtained from human patients. During the Omicron wave, saliva was obtained from 225 patients who tested positive for SARS-CoV-2 <5 days before sampling. On re-testing these stored samples using RT-qPCR, several were negative and excluded from analysis. Saliva is a complex biofluid containing components that include glycosylated mucins and proteins(*45*), which may trap virus leading to difficulty with purification. Initial studies established that gradient centrifugation could not isolate virus from saliva, so magnetic beads coupled to either anti-SARS-CoV-2 spike RBD or human ACE2 were tested instead. Lab grown virus added to a saliva substitute was first tested prior to using patient saliva allowing optimization of virus isolation, with ACE2 beads proving superior (data not shown). Plaque assays performed using saliva, or samples isolated using ACE2-coupled beads confirmed the ability of these beads to recover live virions (Supplementary Figure 8 A-C). In this experiment, 10 RT-qPCR positive saliva samples were pooled, generating a sample with a titer >10^7^ pfu/ml. A two-stage purification method was then established. The first bead isolation (DB1) recovered virus with titers >10^6^ pfu/ml, with the second (DB2) also showing titers >10^6^ pfu/ml (Supplementary Figure 8 B,C). This confirmed that the method successfully recovered infective virus from saliva. Next, 26 patient isolates with high qRTPCR titers were individually subjected to the magnetic bead isolation process. Lipids were then extracted from isolated virus, and analyzed using lipidomics.

Yields of lipid recovered varied between different clinical isolates, reflecting the different levels of virus in individual patients (Figure 4 A). No clear correlation was seen between lipids levels and infectivity measured by RT-qPCR although this was determined prior to virus isolation (Supplementary Figure 8 D). The most abundant lipids were TG, PC and cholesterol. However, plotting data as relative abundance, which corrects for differences in overall yield, and comparing with lab grown Alpha strain virus (since lab grown Omicron was not available when the lipidomics was performed) showed that clinical isolates contained strikingly higher proportions of TG than Alpha, at around 80 % total (Figure 4 B). As salivary glands are a major reservoir for viral replication(*46*), it is possible that salivary lipids may have associated with virus particles and were co-purified in our isolation. Saliva from healthy humans is rather low in lipid overall, with the most abundant being non-polar, including CEs and TG with little PL present(*47–50*). Diseases that impact saliva, such as cystic fibrosis, Sjögren’s syndrome, and periodontal disease are known to lead to dramatic increases in salivary TG(*51,52*). Thus, comparing isolated virus with uninfected controls would not rule out saliva TG contamination, which may be increased in viral infection(*53*). Instead, to test whether the TG might come from other co-purifying particles, a correlation analysis was performed using data on % lipids (molar amounts). Here, lipids that are co-located together, for example based in virus particles, would be expected to correlate with each other. Strikingly, all lipids apart from TGs correlated positively, suggesting co-location in virus particles. In contrast, TG uniquely showed a strong negative correlation with all other categories, indicating it may not be viral in origin (Figure 4 C). It is suspected that TG is associating with virus particles and being co-purified. When compared on a molar basis, there was very low variability seen between individuals for lipid composition indicating that the virus envelope is highly conserved between genetically-unrelated humans (Supplementary Figure 9 A,B). After excluding TG, clinical isolates were found to be relatively similar to lab grown Alpha virus for most PLs and lysoPLs, although there was somewhat less PC, PE and PI, and more free cholesterol detected (Figure 4 D). Reflecting this, the cholesterol:PL ratio was around 4-fold higher, at 2.4, versus 0.53 for the lab grown Alpha strain.

**Figure 4.**
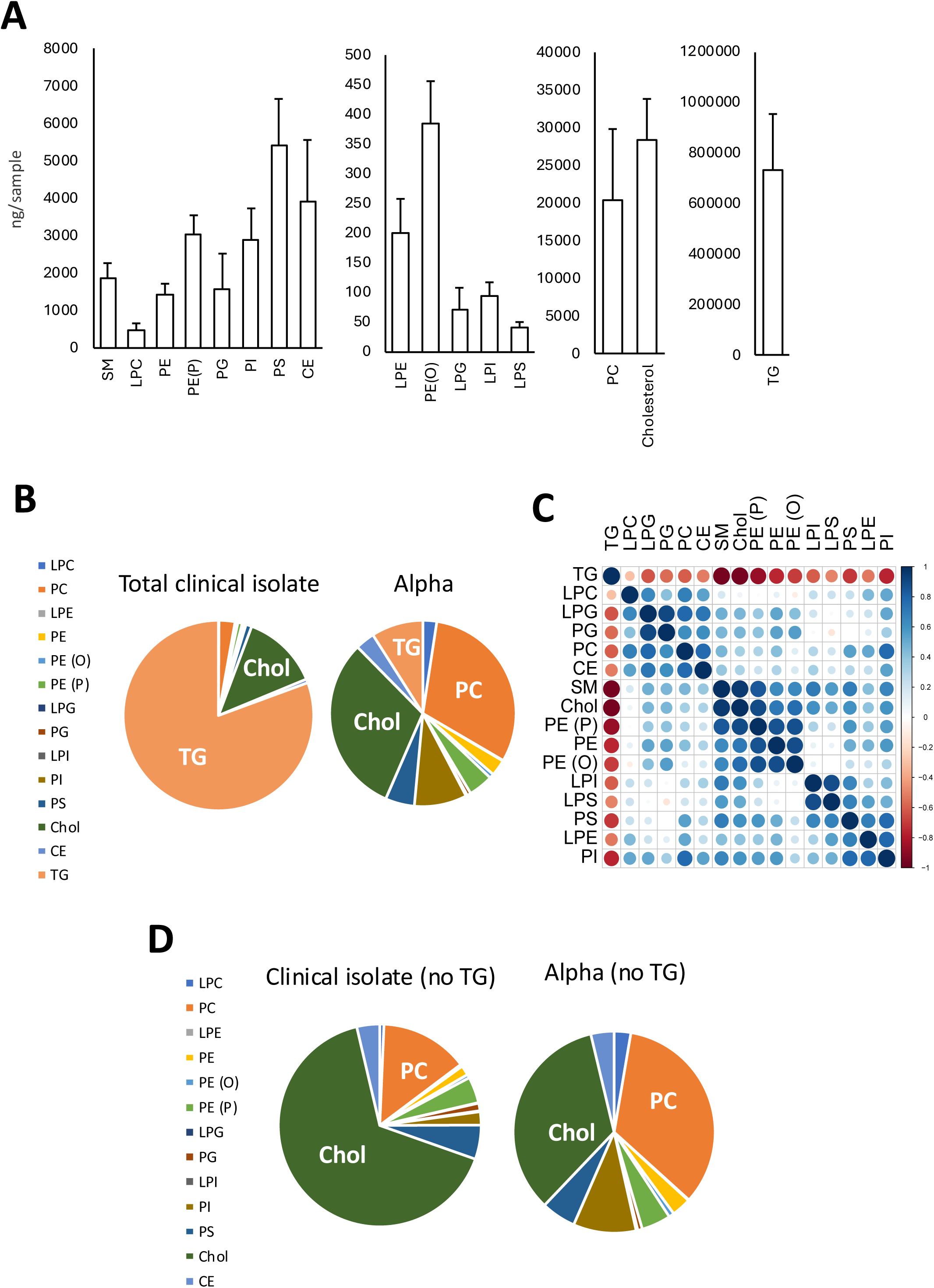
The lipid composition of clinical virus shows SARS-CoV-2 is conserved between individual donors but has a higher cholesterol content than lab grown strains. Virus was recovered from 26 donor saliva isolates using ACE2-coupled magnetic beads as outlined in Methods, then lipids extracted and analyzed using LC/MS/MS. *Panel A. Amounts of lipids recovered from saliva show some variability between individuals.* Lipid amounts (ng/sample) were summed to give total per category (n = 26 donor isolates). *Panel B. Lipid composition of recovered particles shows enrichment with TGs in clinical isolates, as compared with Alpha strain*. Lipid amounts (ng/sample) were converted to molar, totaled and expressed as a % total lipid using an average mass value per category (n=26 or 3, for clinical isolates and lab grown Alpha strain respectively), shown in a pie chart. *Panel C. Correlation analysis suggests that TGs are not originating from virus particles.* Pearson correlation was performed on total lipid molar% values for each isolate (n=26). *Panel D. Lipid composition of recovered particles shows higher levels of cholesterol in clinical isolates, as compared with Alpha strain*. Lipid amounts (ng/sample), omitting TG, were converted to molar, totaled and expressed as a % total lipid using an average mass value per category (n=26 or 3, for clinical isolates and lab grown Alpha strain respectively), shown in pie charts.

### Phospholipid fatty acyl composition of clinical virus shows a distinct pattern to that of Alpha strain

Next, the FA composition of clinical isolates was compared with that of the lab grown Alpha strain. The major molecular species for all categories were generally detected, although there were some significant differences seen in relative abundance (Figure 5 A, Supplementary Figure 9 C,10,11). The most abundant PC and PE species contained 16:0, 18:0, 18:1 and 20:2 for both strains, with unsaturated PUFA such as 20:4, 20:5, and 22:4-22:6 being relatively low abundance (Supplementary Figure 9 C). This was consistent for all PL categories and indicates that the viral envelope is not enriched in unsaturated PUFA when generated *in vivo* in humans (Supplementary Figure 10,11). However, there were some notable PL class-specific differences between the strains. In the case of PE, for either diacyl or ether/plasmalogen forms, molecular species with 18:1 or 18:2 at Sn2 were significantly elevated, while species with 18:3, 20:1, 20:2, 20:3, 20:5 were significantly reduced (Figure 5 A, Supplementary Figure 9 C, 10,11). There were variable differences for species containing the long chain PUFA 20:4, and 22:4-22:6 (Figure 5 A, Supplementary Figure 9 C, 10,11). For PC, species with 18:2, 18:3 were significantly elevated in clinical isolates (Supplementary Figure 9 C, 10,11) but other FA, including from 20:3 onwards were variably changed without a consistent pattern. Higher levels of 18:2 in clinical isolates were consistently seen for all PL categories (Figure 5 A, Supplementary Figure 9 C, 10,11). Similar to PE, PS, PI and PG showed significant decreases in almost all PL species containing 20:1, 20:2 and 20:3 (Supplementary Figure 10,11). In summary, for clinical isolates, PL with C18 FA tended to be higher while C20 lipids tended to be reduced when compared to lab grown Alpha strain. Although we are comparing *in vivo* Omicron with *in vitro* Alpha, we suspect these FA differences result from host cell FA differences, since we earlier found that variants don’t differ greatly for FA composition (Supplementary Figure 2). The FA composition of the salivary gland tissue from which SARS-CoV-2 is generated in humans isn’t known so this can’t be directly compared with AAT cell FA composition in order to characterize these differences more fully.

**Figure 5.**
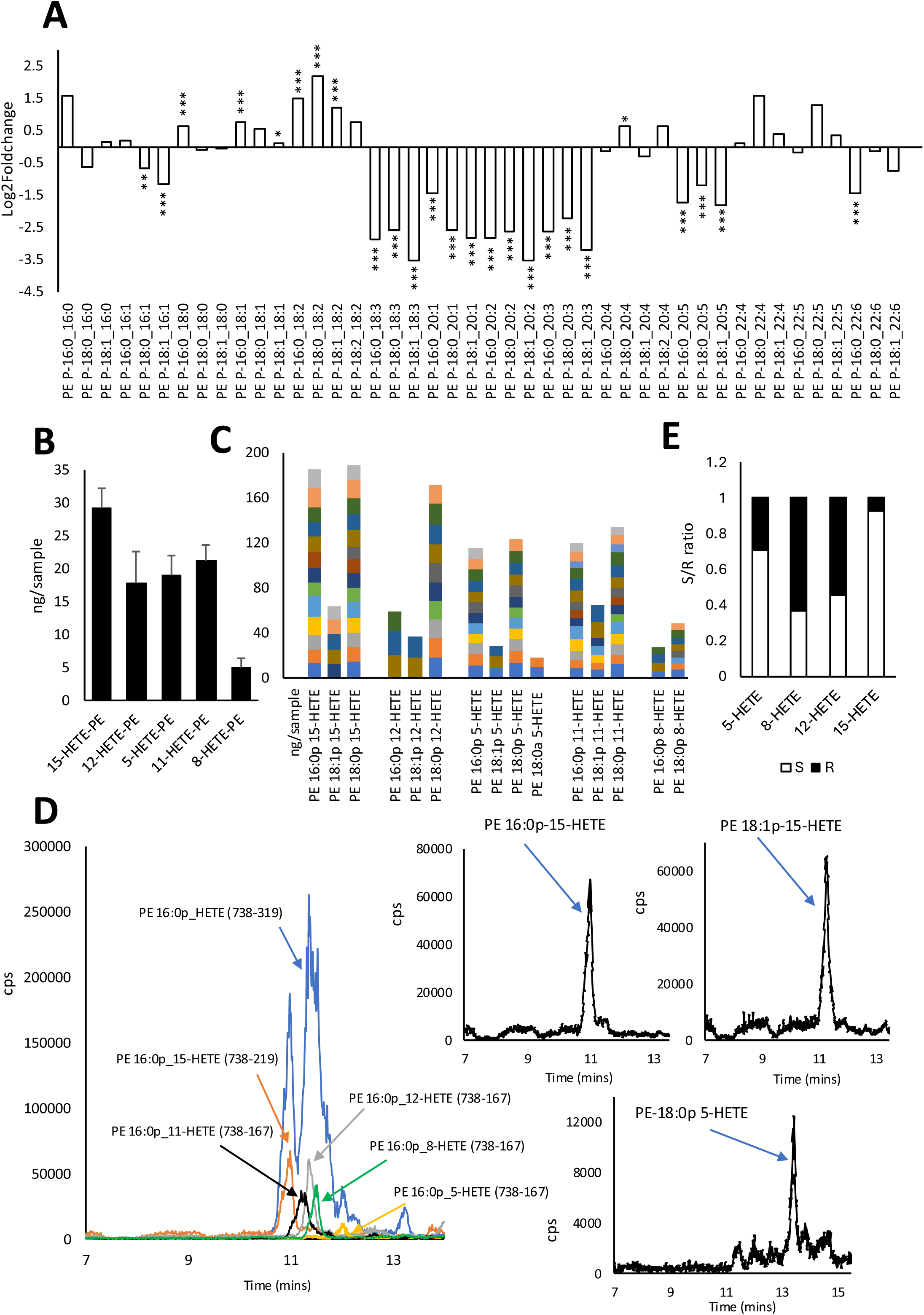
Clinical virus and Alpha strain have altered PUFA profiles in PL, while containing significant amounts of eoxPL from LOX activity. *Panel A. Altered PUFA profiles in plasmalogen/ether-PE comparing clinical and lab grown SARS-CoV-2.* Individual ether/plasmalogen-PE species detected were averaged and compared for an impact of IL-4, and are shown as log2foldchange. A decrease reflects reduction in the species in clinical isolates (n=26, clinical isolates, n=3 Alpha, mean +/- SEM), unpaired Student t-test, * P<0.05, ** P<0.01, ***P<0.005). *Panel B. Oxidized phospholipids are detected in clinical isolates of SARS-CoV-2.* OxPL were quantified in lipid extracts from clinical isolates using LC/MS/MS as outlined in methods, with detected levels in 15 shown (mean +/- SEM). *Panel C.* Data from Panel B replotted as individual molecular species detected in clinical isolates. Each colored bar represents an individual donor. *Panel D. Representative chromatograms showing elution of a series of HETE-PEs on reverse phase LC-MS/MS using primary and secondary MRMs.* Large panel shows a multiple reaction monitoring (MRM) transition corresponding to PE 16:0p_HETE (blue) which doesn’t distinguish positional isomers. Internal daughter ion-based MRMs are shown for 15-,11-,12-,8- and 5-HETE-PE species, showing co-elution with the precursor-319 MRM transition in the expected order. Smaller panels show three individual HETE-PE chromatograms. *Panel E. Chiral analysis shows that 15- and 5-HETEs are predominantly S enantiomers, while 8- and 12- are mixed.* Lipid extracts from a SARS-CoV-2 isolate were hydrolyzed and analyzed using chiral LC/MS/MS for S/R ratios of HETEs, as outlined in Methods.

### Clinical virus contains pro-coagulant oxidized phospholipids from lipoxygenases (LOX)

Inflammatory activation of cells leads to generation of enzymatically oxidized phospholipids (eoxPL) from LOXs, which are pro-coagulant *in vitro* and *in vivo*, through enhancing the ability of PS to bind and activate clotting factors(*54,55*). Generation of truncated oxPL products during infection of A549 cells in vitro, or in vivo in mouse lung, by influenza, was also reported previously(*56,57*). Importantly, in vivo, this coincided with cyclooxygenase-2 (COX-2) induction, which can also form eoxPL(*58,59*). Considering the well-known association of inflammation and thrombosis with COVID19, we reasoned that eoxPL might be generated by inflammatory activation of host cells and transferred to virus particles during replication *in vivo*, where they could modulate the pro-coagulant or infective activities of PS. Screening the 26 clinical SARS-CoV-2 isolates we found several molecular species of ether/plasmalogen PE containing either 15-, 12-, 11-, 8- or 5-HETE and either 16:0, 18:0 or 18:1 at the Sn1 position (Figure 5 B-D, Supplementary Figure 12). While amounts of 15-, 12-, 11- and 5-HETE-PE were overall similar, 8-HETE-PEs were considerably lower at only ∼20% of the average of the others (Figure 5 B). One diacyl (PE 18:0_5-HETE) was detected, but only in 2 isolates (Figure 5 C). OxPL were consistently detected in 15 isolates, corresponding to those with the highest total lipid yield and may be present in the other 11, but below limit of detection. Their isomeric pattern indicates enzymatic origin, since non-enzymatic would result in equal amounts of positional isomers. This was further confirmed through hydrolyzing total envelope lipids and measuring HETE chirality. 15- and 5-HETEs comprised 70 and 94 % S-isomers, respectively, in line with their generation by leukocyte 15- or 5-LOXs, respectively (Figure 5 E). A contribution of cyclooxygenase (COX) is also suggested since around 30 % of the 15-HETE was the R-enantiomer, with this enzyme also generating 11-HETE and 11-HETE-PE in platelets(*60,61*). 12-HETE comprised around equal amounts of both S and R isomers (Figure 5 E). This lipid could be synthesized in humans by platelet 12-LOX (12S-HETE), or epithelial 12R-LOX (12R-HETE), with the latter expressed in epithelial cells of tonsils or salivary glands(*62,63*). 8-HETE-PE is most likely non-enzymatically generated since its levels were far lower than the others, comprising an almost equal mixture of S and R isomers (Figure 5 E).

### Clinical virus contains oxylipins derived mainly from 12-LOXs

To examine the potential for enzymatic generation further, free oxylipins were next analyzed in a subset of clinical isolates. The most quantitatively abundant in every isolate was 12-HETE, which was >5-fold higher than the next, 15-HETE, confirming enzymatic origin (Figure 6 A). Considering that other HETE positional isomers were < 2 % the levels of 12-HETE, both 12- and 15-HETEs are confirmed to come from LOXs. Chiral analysis further confirmed this with 5- and 15-HETE being primarily S enantiomers, consistent with leukocyte-type 5- and leukocyte or epithelial 15-LOXs, while 8- and 12-HETE were mixtures of S and R (Figure 6 B). As for eoxPL above, we propose 8-HETE to be non-enzymatically generated, while the huge amounts of 12-HETE detected likely originate from both platelet 12-LOX and epithelial 12R-LOX. Comparing relative abundance of HEPEs and HDOHEs also further confirmed a role for 12-LOX, with the most abundant being 12-HEPE and 14-HDOHE. (Figure 6 E,F). Indicating 15-LOX activity, 13-HODE was around 5-fold more abundant than 9-HODE, and comprised mainly the S-enantiomer (Figure 6 C,D). A small number of additional oxylipins were measured at levels < 1ng/sample, but these were not consistently detected in all isolates (Supplementary Data). Relatively high amounts of 17-HDOHE were present, but in only of 2/6 isolates, notably the same 2 isolates with highest levels of 15-HETE, supporting the idea that these two are from 15-LOX. 7,17-diHDOHE was present in the same two isolates (only) at very low levels, and co-eluted with the resolvinD5 standard, however considering the low levels of the lipid in comparison to 7-HDOHE and 5-HETE, we assume that 5-LOX activity is unlikely contributing to 7,17-diHDOHE formation in these isolates and non-enzymatic oxidation of 17-HDOHE is more likely. In summary, the positional isomer and chiral patterns of HETEs, HODEs, and HEPEs/HDOHEs strongly evidence several LOXs to be contributing to oxylipins carried by the virus, with little or no contribution from P450 or cyclooxygenases.

**Figure 6.**
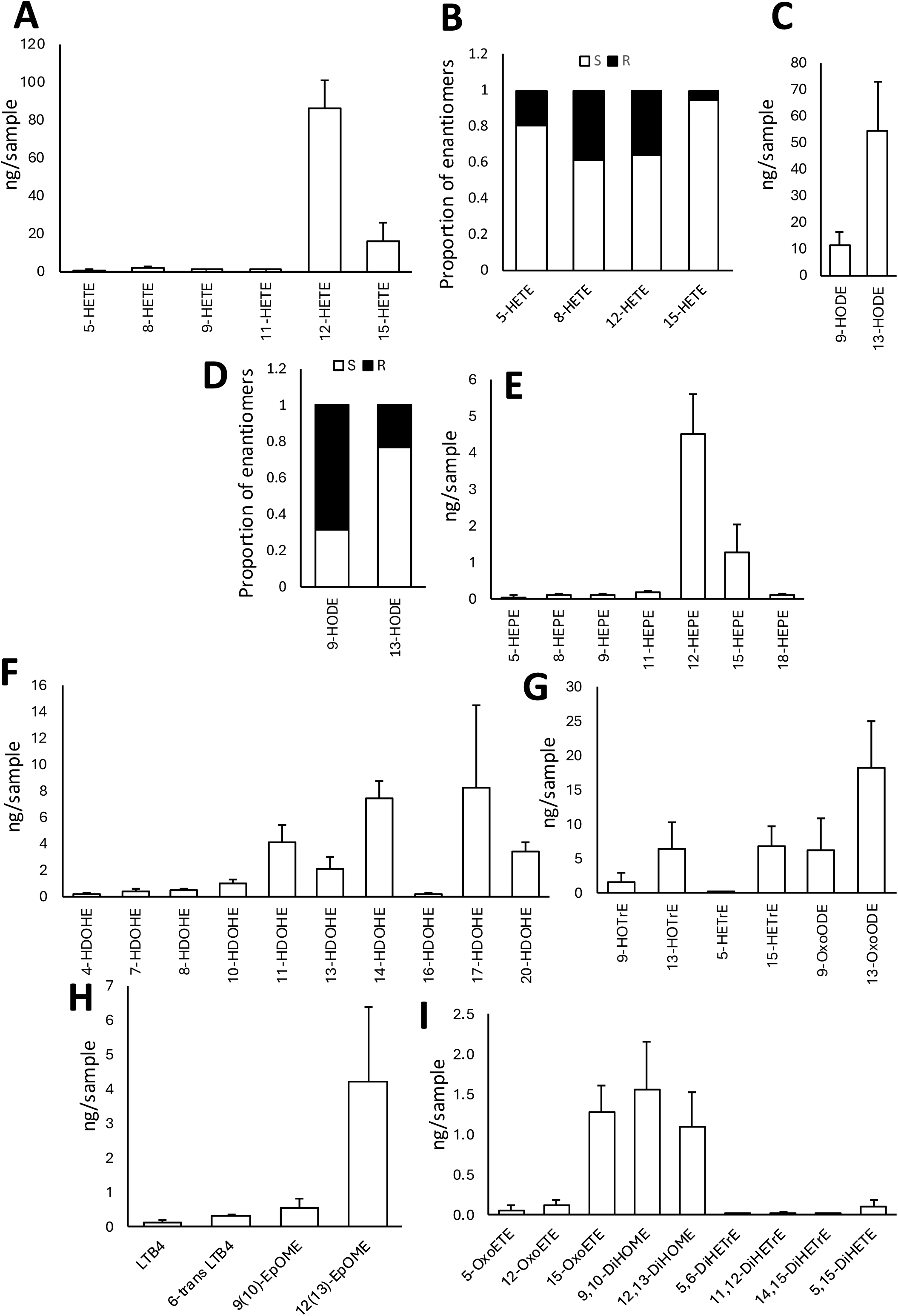
**Clinical virus contains several oxylipins primarily LOX products using reverse phase and chiral LC/MS/MS**. *Panel A. HETE positional isomers show a dominance of 12- and 15-HETEs*. Lipids were analyzed for oxylipins in extracts of clinical virus using LC/MS/MS as outlined in Methods (n=6, mean +/- SEM). *Panel B. Chiral analysis indicates enzymatic origin of free HETEs*. Chiral analysis of lipid extracts from clinical samples was carried out as indicted in methods (n=4 isolates). *Panel C. HODE positional isomers show a dominance of 13-HODE*. Lipids were analyzed for oxylipins in extracts of clinical virus using LC/MS/MS as outlined in Methods (n=6, mean +/- SEM). *Panel D. Chiral analysis indicates enzymatic origin of free HODEs*. Chiral analysis of lipid extracts from clinical samples was carried out as indicted in methods (n=4 isolates). *Panels E-I. Oxylipin analysis shows further dominance of 12-LOX products, 12-HEPE, 14-HDOHE, along with other lipids*. Lipids were analyzed for oxylipins in extracts of clinical virus using LC/MS/MS as outlined in Methods (n=6, mean +/- SEM).

### ALOX15B-silenced macrophages are less susceptible to viral infection with coronaviruses

Human cells express two 15-LOX isoforms, with *ALOX15B* (15-LOX2) widely expressed in epithelial cells, as reviewed here(*64*). This isoform can generate both free 15-HETE, and esterified forms including 15-HETE-PEs. Considering the tissue expression of *ALOX15B*, and that lipid peroxidation is required for replication of zika(*65*), another enveloped virus which buds from ER membranes, we decided to test whether *ALOX15B* expression impacts coronavirus infection. Notably, the nature of lipid peroxidation involved in zika replication is currently unknown. Here, two other coronaviruses that infect macrophages were tested, MERS-CoV and HCoV-229E. To test the impact of *ALOX15B*, macrophages lacking the gene through siRNA knockdown were compared with control cells which strongly express this isoform, with both viruses known to productively replicate in macrophages(*66,67*).

Analysis of MERS-CoV- and HCoV-229E-encoded gene expression 24 hours post-infection (hpi) indicated significantly reduced viral loads in *ALOX15B* KD cells compared to control siRNA-transfected macrophages (Figure 7A,B). To delineate whether reduced load at 24 hpi was mediated by altered viral entry or through a post-entry process, macrophages were infected with MERS-CoV and viral load assessed in a time-dependent manner from 2 to 48 hpi. Although not significantly different at 2 or 4 hpi, viral load in ALOX15B KD cells was significantly reduced at 8 hpi and was continually diminished at 24 and 48 hpi (Figure 7 C). As viral load at 2 hpi was not significantly different between KD and control cells, we normalized load to assess rate of replication. Here, MERS-CoV replication from 8 to 24 hpi was significantly reduced in ALOX15B-attenuated macrophages (Figure 7 D). These data suggest that *ALOX15B* KD influences post-entry processes rather than viral entry.

**Figure 7.**
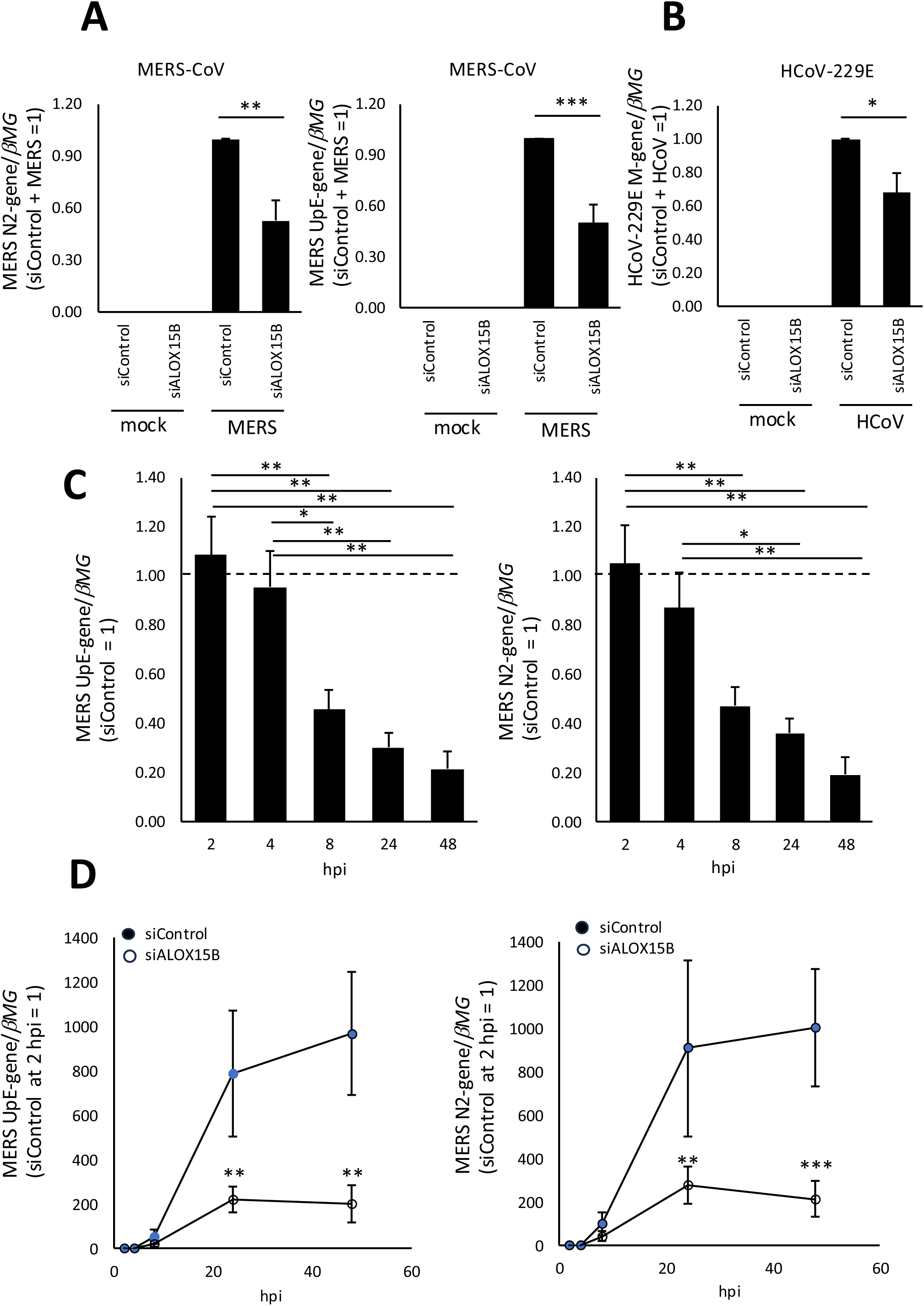
**ALOX15B is required for coronavirus replication in macrophages**. *Panel A.* Gene expression of Middle East respiratory syndrome (MERS)-(coronavirus)-CoV nucleocapsid 2 (N2)-gene and locus upstream from envelope (upE)-gene at 24 hours post-infection (hpi) *Panel B.* Gene expression of human CoV (HCoV)-229E membrane (M)-gene in control and ALOX15B KD macrophages at 24 hours post-infection (hpi). *Panel C.* Relative expression of MERS-CoV upE-gene or N2-gene in ALOX15B KD macrophages at 2, 4, 8, 24, and 48 hpi compared to control cells. *Panel D.* Gene expression of MERS-CoV upE-gene or N2-gene in control and ALOX15B KD macrophages at 2, 4, 8, 24, and 48 hpi after normalizing to viral load at 2 hpi (mean ± SEM from at least four independent experiments, two-tailed student’s t-test for MERS treatment in A and B, one-way ANOVA with Tukey’s post hoc test for Panel C, and two-way ANOVA with Sidak’s multiple comparisons test for D (*P<0.05, **P<0.01, ***P<0.001 and ****P<0.0001 vs siControl).

### The impact of ALOX15B on replication is independent of cholesterol or intermediates in its biosynthetic pathway

Many viruses target membrane cholesterol for viral binding and entry into host cells(*68,69*). Furthermore, replication was shown to require intermediates from the mevalonate and sterol biosynthesis arms of the cholesterol synthesis pathway, including mevalonate, squalene, and geranylgeraniol (GGOH)(*70–72*). As *ALOX15B*-silenced macrophages show reduced levels of cholesterol precursors desmosterol and lathosterol(*73*), levels of plasma membrane-accessible cholesterol, and whether deficiency in sterol intermediates contribute to the reduced viral replication observed was next determined. Plasma membrane cholesterol exists in three operational pools referred to as accessible, sphingomyelin (SM)-sequestered, and essential(*74*). Accessible cholesterol is not sequestered by SM or other phospholipids. When its level in the plasma membrane exceeds a certain threshold, the excess cholesterol rapidly traffics to the ER to signal cholesterol surplus(*74,75*).

To assess the pool of accessible PM cholesterol the cholesterol-dependent cytolysin (CDC) streptolysin O (SLO) derived from *Streptococcus pyogenes* was used(*76*). This is a pore-forming toxin targeting accessible PM cholesterol for its effector function(*77*). When paired with the membrane-impermeable dye propidium iodide (PrI), SLO functions as an effective tool to assess accessible cholesterol levels(*77,78*) (Supplementary Figure 13 A). Naïve macrophages co-treated with SLO and PrI, but not vehicle (DTT) and PrI, displayed red intracellular fluorescence when analyzed using Incucyte® live-cell imaging (Supplementary Figure 13 B). Quantification of PrI-mediated fluorescence using flow cytometry showed significantly diminished PrI signal in *ALOX15B* KD compared to control siRNA-transfected macrophages (Supplementary Figure 13 C). ML-351, a 15-LOX inhibitor, also significantly reduced PrI-mediated fluorescence in non-transfected macrophages (Supplementary Figure 13 D). Replenishment with exogenous cholesterol, using cholesterol-complexed methyl-β-cyclodextrin (C/MβCD), increased SLO-dependent pore formation (PrI-mediated fluorescence) in *ALOX15B* KD as well as control macrophages treated with SLO (Supplementary Figure 13 E). These data indicate that *ALOX15B*-silenced macrophages exhibit reduced accessible plasma membrane cholesterol, in tandem with reduced levels of precursors as previously shown(*73*).

Next, a metabolite rescue experiment in which control and *ALOX15B* KD macrophages infected with MERS-CoV were subsequently treated with mevalonate, GGOH, lathosterol or desmosterol was performed. However, in contrast to previously published results showing administration of mevalonate and GGOH could rescue viral growth in sterol regulatory element-binding protein (SREBP) 2-inhibited cells(*70–72*), neither mevalonate, GGOH nor lathosterol or desmosterol increased MERS-CoV viral load in *ALOX15B* KD cells (Supplementary Table 8). This suggests that intermediates in the mevalonate and sterol biosynthesis arms of the cholesterol synthesis pathway are not involved with *ALOX15B*-promoted viral infection. How *ALOX15B* deletion contributes to antiviral protection in primary human macrophages remains to be further investigated.

## DISCUSSION

In this study, the lipid envelope of SARS-CoV-2 was characterized, focusing on four human pandemic VOCs and *ex vivo* isolated virus from patients. An impact of common inflammatory factors such as cytokines and phospholipases was revealed, with the virus discovered to act as a reservoir for numerous bioactive, thrombotic and pro-coagulant lipids, some required for coronavirus replication. Our studies reveal many new findings concerning the biology of this so far poorly understood component of an important human pathogen. They indicate that coronavirus envelopes are highly conserved between variants but also respond acutely to host factors, generating a new paradigm for our understanding of virus biology, and paving the way for future studies on the envelopes of other important enveloped viruses.

Surprisingly, the viral membrane of coronaviruses or indeed any enveloped viruses has not been well studied before now, despite the fact it represents the environment in which proteins that drive infection reside, as well as a potential repository for many bioactive lipids. It is critically important to understand how changes in membrane composition might influence the ability of viruses to infect cells, as well as drive downstream pathology, such as inflammation and thrombosis. While the four SARS-CoV-2 virus strains cultured in vitro were quite similar overall, virions purified from human saliva were quite different, with reduced amounts of specific C18 and C20-containing species. The altered FA composition likely reflects the differences in lipids contained in AAT cells versus cells of the salivary gland where SARS-CoV-2 is known to replicate *in vivo*, including acinar and ductal epithelium(*46,79*). This suggests that studies on in vitro generated viruses need to consider how the host environment in vivo might alter membrane composition, with downstream impacts on membrane biology.

Up to now, little research has been conducted into the membrane composition of enveloped viruses with the only mass spectrometry characterization so far published on HIV (*80–82*) and influenza (*83,84*). Prior studies before this in the 1960s-1980s on viruses that included HIV, Sendai, Newcastle Disease Virus and Vesicular Stomatitis Virus used older methods such as thin layer chromatography, which do not provide molecular species information on individual lipid classes(*85–87*). Furthermore, no studies have so far attempted to profile the lipids of virus isolated from infected human hosts, examine how a virus envelope responds to host factors or profile aPL externalization, oxPL or oxylipins. Notably, all these viruses bud from plasma membrane and in reflection of this, their cholesterol/PL ratio is far higher, around 0.7-1 than we found for SARS-CoV-2, which was typically around 0.4-0.7 reflecting a similar ratio to intracellular organelles such as lysosomes from where it is proposed to bud(*88*). Unexpectedly, the cholesterol:PL ratio for SARS-CoV-2 isolated from human saliva was around 4-fold higher, at 2.4, versus 0.53 for the lab grown Alpha strain. This shows a difference between *in vitro* and *in vivo* replicated virus, which could have implications for the ability of membrane disrupting agents to inactivate the virus(*3*).

Notably, global lipid composition of virus from four SARS-CoV-2 VOCs was well conserved. This shows that mutations in viral proteins which can lead to changes in immunogenicity, infectivity or symptoms in human disease don’t drive major alterations in envelope lipids. Importantly, all four strains expressed large amounts of PE and PS on their outer surface, with two strains reaching 70-80%. This level of externalization is extremely high considering activated platelets only expose up to around 7-10 % during blood clotting(*15*) and reveals some strains to be likely “super activators”, should they come in contact with activated coagulation factors. Indeed, we previously showed that the England2 strain activates plasmatic coagulation at virion levels that are considerably lower than those reported in BAL, saliva or subglottic aspirate(*3*), and a direct association of viremia with thrombotic complications was recently demonstrated in patients(*89*). Furthermore it may underscore the importance of PS as a factor enhancing SARS-CoV-2 entry(*16*). How the virus achieves this level of “asymmetry” for PS and PE in some strains is unknown since it doesn’t bud from the plasma membrane. Recent studies point to membrane asymmetry of ER during lipid droplet budding that relate to an excess of PL in one monolayer however, the full extent of this and how it might be regulated by scramblases or flippases is still completely unknown(*90,91*).

Four factors known to alter mammalian cell lipid metabolism that are directly relevant to COVID19 were tested for their ability to influence viral envelopes, namely IL-4, IL-6, dexamethasone and a series of sPLA2 isoforms. Of these, IL-4 led to dramatic upregulations of lipid along with significantly reduced PUFA content across all lipid categories, while all sPLA2s tested directly hydrolyzed the membrane. In direct support of our findings with IL-4, several studies found that this cytokine increases PL and FA biosynthesis in mammalian cells including endothelial and macrophages via SREBP1(*92–94*). Furthermore, SREBP1 activation (by M-CSF) dramatically increases FA synthesis while reducing PUFA containing PL during macrophage differentiation(*95,96*), while IL-13 (which signals via the same receptors as IL-4) elevates PC in mouse lung(*97*). Last, providing a mechanistic explanation for the changes in PUFA, IL-4 downregulates elongation and desaturation enzymes responsible for their generation from 18:2/18:3 FA in cultured epithelial and lung cells(*98,99*). These changes may have pathological significance for replication since SARS-CoV-2 infection leads to a dramatic increase in PUFA species across all lipid categories, and virus survival depends on de novo FA synthesis(*20*). Considering this, factors that disrupt the PUFA balance in host cells may have knock-on effects on viral envelope composition potentially impacting replication. The significant changes on IL-4 treatment of cells may also lead to changes in membrane fluidity or other biochemical parameters, long ascribed to long chain FA contained in PLs(*100*). These could include increasing curvature and flexibility, reducing lipid packing, and causing membrane thinning(*101–104*). Relating to sPLA2, theories have been proposed to explain its association with higher COVID mortality, relating to its ability to hydrolyze apoptotic cells releasing cellular content, exposing more PS and driving thrombosis, as well as providing substrates for eicosanoid and prostaglandin biosynthesis(*40,41*). Our study extends this by showing that the virus is a direct target for sPLA2, indicating that its impact on virus biology needs further investigation.

Clinical virus contained oxylipins, both free and PL-esterified, dominated by isoforms generated by LOX enzymes, particularly 12-, 15- and 5-LOXs, as based on both the pattern of positional isomers and their chirality. 12-HETE could be synthesized in humans by platelet 12-LOX (12S-HETE), or epithelial 12R-LOX (12R-HETE)(*62,63*). Consistent with this, 12R-LOX (ALOX12B) is present in the SARS-CoV-2 proteome of *in vitro* cultured SARS-CoV-2(*105*). For both 5- and 15-HETE, the dominance of S enantiomers, as well as 17-HDOHE, and a high 13-HODE:9-HODE ratio is consistent with generation by leukocyte-type 5- and either leukocyte or epithelial 15-LOXs. Oxidized phospholipids are widely considered as DAMPs generated at sites of injury to signal to the innate immune system. Several studies have shown that they are generated *in vitro* and *in vivo* in host cells or tissues following respiratory infection including influenza(*106–109*), but up to now this was assumed due to production of reactive oxygen species that generate non-enzymatically-derived species. However, previous studies used either antibody-based methods or measured truncated end products of PL oxidation only, and it was not determined whether enzymes were involved in their formation(*106–109*). Clarifying this, for SARS-CoV-2, using human clinical isolates, we demonstrate involvement of LOXs using lipidomics, and suggest that the origin of oxPL in other forms of enveloped viral infection should be re-evaluated. Furthermore, the presence of eoxPL in virus envelopes raises intriguing questions about potential function since these lipids are well known to be pro-coagulant through enhancing the ability of PS to drive thrombin generation(*110–112*). Considering that *ALOX15B* is widely expressed in epithelia, our findings that deletion of this isoform significantly reduces viral replication directly link its lipids with virus survival and are consistent with reports of lipid oxidation being also required for zika virus replication(*65*). In the zika study, which showed that free radical scavenging agent liproxstatin-1 prevents replication, the origin of the PL oxidation was not defined, and a role for LOXs is also possible. Our data suggest that local targeting of lipid oxidation, for example using ferroptosis inhibitors or downregulating LOX expression could be evaluated as a novel anti-viral strategy prior to vaccination.

While our studies focused on coronaviruses, mainly SARS-CoV-2, the UK Health Security Agency and WHO priority viral pathogens are almost all enveloped, including Ebola, Lassa, MERS, SARS, Rift Valley fever and zika, with currently nothing known about their membranes(*113,114*). This highlights a major information gap, and the importance of understanding this unique aspect of virus biology. Our studies reveal that lipid envelopes are not simply scaffolds supporting viral proteins and the genome, but actively participate in replication, integrity, and disease pathogenesis, and need to be better understood and repositioned as an important modifiable focus for anti-viral therapies across all enveloped strains, which could be targeted through host innate immunity.

## MATERIALS AND METHODS

### Cells and viruses

Human airway epithelial A549 and monkey kidney epithelial VeroE6 cells were gifts from the University of Glasgow/MRC Centre for Virology, UK. To enhance infectivity and produce a more sensitive cell line for detection of virus, both cell types were transduced with lentiviruses encoding ACE2 and TMPRSS2, two critical proteins for entry of the virus, and then drug selected as described creating the Vero A/T or AAT cell lines(*115*). Four different strains of SARS-CoV-2 were used: England2 (Wuhan), Alpha (Kent, B.1.1.7), Beta (South Africa, B.1.351) and Delta (India, B.1.617.2), provided by Public Health England. All viral were strains amplified in Vero A/T cells before being harvested from the supernatant and titrated in AAT (England2) or AA (Alpha, Beta, Delta) cells for subsequent experiments. All cells were grown in DMEM containing 10 % FCS until infected, after which they were maintained in 2 % FCS, and incubated at 37 °C in 5 % CO2.

### Cell culture and harvest of in vitro generated virus particles

Culture and infection of AAT cells with virus is outlined in full in Supplementary Methods, including assessment of AAT cells for IL-6 responsiveness, and treatment of cells with IL-6, IL-4, dexamethasone and sPLA2 isoforms.

### Collection of clinical samples, and isolation of virus

Clinical samples were collected under Research Ethics Committee approval (REC 21/PR/1189, IRAS 299479) with informed consent with Infection Services for Public Health Wales and University Health Boards across Wales (Aneurin Bevan, Betsi Cadwaladr, Cardiff and Vale, and Cwm Taf Morgannwg) in accordance with the Declaration of Helsinki. Inclusion/exclusion criteria, harvest and storage and isolation of live SARS-CoV-2 using human ACE2-coupled magnetic beads (Acro Biosystems) are outlined in Supplementary Methods.

### Lipid extraction for lipidomics profiling

Lipids were extracted from virus particles following addition of internal standards, as outlined in Supplementary Methods.

### Targeted LC/MS/MS analysis of virus lipids

Several targeted assays were used to analyze lipids from *in vitro* cultured, or *ex vivo* harvested virus, including for PL, SM, lysoPL, cholesterol, CE, TGs, external facing PE and PS, oxPL and oxylipins. Chiral analysis was used to determine S/R ratio of oxylipins. All methods are detailed in full in Supplementary Methods.

### RNA extraction

Nucleic acids were extracted from saliva using a protocol adapted from the automated Bio-On-Magnetic-Beads (BOMB) COVID-19 protocol(*116*), with full details in Supplementary Methods.

### Plaque assay

Viral titers were determined using a standard plaque assay, as outlined in Supplementary Methods.

### RT-qPCR analysis of SARS-CoV2

Reverse transcriptase-polymerase chain reaction (RT-PCR) was used for detection of three gene regions, SARS-CoV-2 nucleocapsid N1 and N2, and the small envelope protein (E) gene, as outlined in Supplementary Methods.

### Virus propagation, and infection of ALOX15B-silenced macrophages

For virus propagation, Caco-2 or Vero cells (DSMZ, Braunschweig, Germany) were used as outlined in Supplementary Methods. Human peripheral blood mononuclear cells were isolated from commercially obtained buffy coats of anonymous donors (DRK-Blutspendedienst Baden-Württemberg-Hessen, Institut für Transfusionsmedizin und Immunhämatologie, Frankfurt, Germany) using Ficoll density centrifugation, differentiated and infected as outlined in Supplementary Methods. Infection was determined by intracellular viral RNA by qPCR at different time points, with primers listed in Supplementary Methods. In some experiments, sterol intermediates were included, as outlined in Supplementary Methods. Live cell imaging and measurement of accessible cholesterol is described in full in Supplementary Methods.

### Statistics

A full description of statistical approaches and strategies used is provided in Supplementary Methods.

## ACKNOWLEDGEMENTS

We gratefully acknowledge assistance from Linda Chan (Morriston Hospital) and Ameerah Azmil (Royal Glamorgan Hospital) and Bethan Gibson (Cwm Taf Morgannwg) for sample collection.

## AUTHOR CONTRIBUTIONS

Conceptualization: VOD, RS, SAJ, PG, BB, DWT, Methodologies: VJT. Reagents: GL, LR, CAB, JH, Investigation: VJT, GL, AZ, MYE, GV, FM, RGS, MBP, EHS, BB, YB, JK, DB, RG,

Funding acquisition: VOD, DT, RGS, PG, SAJ, Project administration: VOD, WP, Supervision: VOD, DWT, RS, BB, Writing – original draft: VOD, BB, Writing – review & editing: VJT, AZ, WP, MYE, VG, FM, MBP, EHS, SAJ, GL, BB, RGS, DB, JK, YB, DWT, RJS, VOD.

## DATA AND MATERIALS AVAILABILITY

All data are available in the main text or the supplementary materials file.

## SUPPLEMENTARY MATERIALS, METHODS AND DATA

### Cell culture and harvest of in vitro generated virus particles

70% confluent AAT cells were infected for 1h with SARS-CoV-2 at MOI=0.05 for the England2 or MOI=0.01 for the VOCs (Alpha, Beta, Delta). Flasks were rocked for 1 hour, and then the inoculum was removed and replaced with fresh DMEM/2% FBS. For virion purification, supernatants were harvested, cellular debris pelleted (2,000 g, 5 min), then virus pelleted through a 5 ml of 30 % sucrose cushion (25,000 rpm, 2.5 h, 4 °C, in a SW28 rotor (112,398 × g)). Pellets were resuspended in 1 ml PBS and purified by density gradient centrifugation on a 20 – 60 % sucrose or a 15 – 40% OptiPrep (Iodixanol, Sigma DI556) gradient (25,000 rpm, 16 h, 4 °C, in a SW41 rotor (106,882 × g)), before being pelleted (25,000 rpm, 2.5 h, 4 °C, in a SW41 rotor (106,882 × g)), and resuspended in 0.5 ml of PBS for subsequent lipid extraction as outlined below. Preparations were analyzed for purity and abundance by nanoparticle tracking analysis using the NanoSight-NS300® (Malvern Panalytical), and by Western blot. For PE and PS externalization, samples were used immediately as fresh. For lipidomic profiling they were used immediately or stored for a few days at -80°C as snap frozen pellets. Samples were processed together, with n=3 biological replicates per variant. During integration of peaks, it was noted that one isolate of England2 had lower levels of lipids overall reflecting less recovery of virus. This unfortunately led to a higher number of lipids being below LOD/LOQ for some lipid categories (TG, LPE, PE, PE-O, PS) and these have been omitted from analysis. No other lipid categories or samples were affected with n = 3 measures generated for the majority of species in this comparison.

Before testing the impact of IL-6 stimulation on viral envelope, the expression of gp130 and IL-6Rα were assessed on AAT cells as follows: Uninfected cells were harvested using TrypLE Express (Gibco, Thermo Fisher Scientific), washed in cold PBS and stained with LIVE/DEAD Fixable Aqua (Thermo Fisher Scientific), PE anti-human CD130/gp130 (clone 2E1B02, cat no. 362003, BioLegend), PE anti-human CD126/IL-6Rα (clone UV4, cat no. 352803, BioLegend), PE Mouse IgG1, κ isotype (clone MOPC-21, cat no. 400112, BioLegend), or PE Mouse IgG2a, κ Isotype (clone MOPC-173, cat no. 400212, BioLegend), followed by a secondary Alexa Fluor 488 goat anti-mouse IgG (cat. no. A-11029, Thermo Fisher Scientific; 1:500). All the above were done as single stains for the cell markers along with the Live/Dead staining. After staining, cells were fixed with 4 % PFA for 30min at 4 °C and washed with PBS before analysis. Data were acquired using an Attune NxT Flow Cytometer (Thermo Fisher Scientific) and analyzed with Attune NxT software or FlowJo software, version 10 (Tree Star). Particles were counted using NanoSight.

To test the impact of IL-6 on viral envelope composition, ‘England2’ strain was generated from AAT stimulated with i) IL-6 (30 ng/ml, BioTechne, 206-IL-010) alone or ii) IL-6 (30 ng/ml) and sIL-6Rα (200 ng/ml, BioTechne, 227-SR-025) together, added to cells 1 hr after virus infection (MOI 0.05) in 2% FCS. Virus was harvested after 72 hrs. To test the impact of IL-4, AAT cells were cultured in the presence of 10 ng/ml IL-4 (rhIL-4, BIO-TECHNE) for 5 days (with 10% human AB serum, Cambridge Biosciences) prior to infection with ‘England2’ strain virus at MOI 0.05, at 2% FCS. Cells were then cultured for 72 hrs before virus was harvested.

For dexamethasone (Dex) treatment, two separate protocols were used: pre- and post-infection. For pre-infection treatment, AAT cells were first cultured for 3 days in DMEM containing FBS which had been charcoal-absorbed to remove glucocorticoids. Then they were cultured with or without 100 nM Dex (Merck Life Science) at low FCS (2%, charcoal stripped) for 24 hrs. Next, AAT were infected with SARS-CoV-2 England2 at MOI 0.05, in 2% FCS, and cultured for 48h before virus was isolated as detailed below. For post-infection, AAT cells were first cultured for 3 days in DMEM containing FBS which had been charcoal-absorbed to remove glucocorticoids. Then, 100 nm Dex was added 1 hr after inoculation of virus and then again once each day during culture prior to virus harvest.

### Incubation of virus with sPLA2 isoforms

Eight sPLA2 were tested for their hydrolytic activity towards SARS-CoV2 envelopes, as listed in Supplementary Table 1), including both recombinant human and purified venom isoforms. Virus (England2) isolated from AAT supernatants using density gradient centrifugation as described above was resuspended in 100 mM NaCl, 20 mM Tris-HCl, 2 mM CaCl2, 0.05% BSA (pH 7.4). Virus was incubated in 10 nM sPLA2 for 6 hr at 37 °C. 20 mM CaCl2 was added, the samples snap frozen to -80 prior to addition of internal standards, and lipid extraction and analysis performed as described below.

### Collection of clinical samples

Following Research Ethics Committee approval (REC 21/PR/1189, IRAS 299479) and informed consent, saliva samples were obtained from 225 individuals (1 per donor), within 5 days of a positive test for COVID19 (via PCR, lateral flow or licensed point of care testing; according to UKHSA recommendations). Samples were collected working with Infection Services for Public Health Wales and University Health Boards across Wales (Aneurin Bevan, Betsi Cadwaladr, Cardiff and Vale, and Cwm Taf Morgannwg) in accordance with the Declaration of Helsinki. Inclusion criteria included: Age 18-75 yrs, ability to provide consent, not having used topical antimicrobial oral therapies in the preceding 12 hours. Exclusion criteria included: known pregnancy or breastfeeding, requirement for ventilation and ingestion of food in the 1 hour prior to donating saliva. Unstimulated saliva (<3-5 ml) was collected into sterile 30 mL universal tubes, which were then, immersed in 70 % (v/v) alcohol, enclosed in 2 sterile ziplock bags ("double-bagged"), treated with alcohol and transported to HTA designated -80 °C freezers in rigid HPA-approved containment. Following transfer to the analytical laboratory, samples were stored at -80 °C until lipidomic analysis was performed as outlined below.

### Isolation of SARS-CoV-2 from clinical samples

Prior to further analysis, samples were tested for SARS-CoV-2 positivity using RTqPCR as described below. Following this, an optimization experiment was undertaken on 10 positive samples using human ACE2-coupled magnetic beads (Acro Biosystems) to purify virus as follows: beads were washed to remove trehalose then reconstituted in PBS, as per manufacturer’s instructions. 5 ml beads were added to 20 ml pooled saliva which was placed on a rotator at 37 °C for 1 hr before being placed on a 3D printed magnetic separator for 2 min. Beads with bound virions (DB1) were recovered separately to the remaining supernatant. Supernatant was then added to another 5ml beads. As above, this was placed on a rotator at 37 °C for 1 hr before being placed on the magnetic separator rack for 2 min. Beads with virions were again recovered (DB2). DB1 and DB2 were stored at -80°C before being tested for infectivity using plaque assay as described later. Next, for lipidomics of SARS-CoV-2, 26 RTqPCR positive isolates were selected, with volumes ranging from 0.2-14 ml (Supplementary Table 2). To each, washed ACE2-beads were added at ratio of 1:5 (beads:sample). Magnetic separation was then carried out as described for the optimization experiment above. Samples were stored at at -80 °C prior to lipidomics. Recovered virus from DB1 and DB2 were separately extracted and analyzed using LC/MS/MS, then the two datasets from each individual were pooled to provide a single lipidomics dataset per patient.

### Lipid extraction for lipidomics profiling

Isolated virus particles were resuspended in 0.5 ml of PBS, which was then spiked with 10 μl EquiSplash mix (Avanti Polar Lipids), containing d18:1-18:1(d9) SM (100 ng), 15:0-18:1(d7) phosphatidylcholine (PC) (100 ng), 15:0-18:1(d7)PE (100 ng), 15:0-18:1(d7) phosphatidylglycerol (PG) (100 ng), 15:0-18:1(d7) phosphatidylinositol (PI) (100 ng), 18:1(d7) Lyso PC (100 ng), 18:1(d7) Lyso PE (1000 ng), cholesterol-d7 (100 ng), cholesteryl ester (CE) 18:1-d7 (100 μg), triglyceride (TG) 15:0/18:1-d7/15:0 (100 ng), 15:0-18:1(d7) PS (100 ng), and 100 ng of 17:1 Lyso PG, PS and PI. Samples were also spiked with 5 ul of Cer/sphingoid internal standard mixII (Avanti Polar Lipids) containing 56.99 ng of d18:1/12:0 Cer. Samples were then extracted using an acidified Bligh and Dyer method. Briefly, 1.9 ml of solvent mixture, 2:1 methanol:chloroform v:v, was added to 0.5 ml sample acidified with 10 ul glacial acetic acid. Samples were vortexed for 30 s, and then 0.625 ml of chloroform added. Samples were vortexed again (for 30 s), and 0.625 ml of water added. Samples were vortexed for 30 s and centrifuged at 500 g, at 4°C, for 5 min. Lipids were recovered from the lower layer and evaporated to dryness using a Labconco RapidVap® (Labconco). Extracted lipids were reconstituted in 200 ul methanol and stored at −80°C until analysis.

### Targeted LC/MS/MS analysis of native lipids

Targeted assays were performed on culture preparations of gradient-purified SARS-CoV-2 virus, from A549 cells, and clinical isolates. A full list of all lipids analyzed is shown in Supplementary Table 3. with data on extraction efficiency and instrument coefficient of variation. We note that PEs annotated as plasmalogen (vinyl ether) could also comprise isobaric ether lipids. For the analysis of the clinical samples, PC species were monitored as [M-CH3]^-^. For all other experiments, PCs were monitored as acetate adducts. Standard curves were run at the same time in all instances to account for any variation. Hydrophilic interaction liquid chromatography (HILIC) LC-MS/MS was used for PLs and SLs on a Nexera liquid chromatography system (Shimadzu) coupled to an API 6500 qTrap mass spectrometer (Sciex). Liquid chromatography was performed at 35°C using a Phenomenex Luna, 3 μm NH2 column, 2 × 100 mm, at a flow rate of 0.2-0.7 ml/min over 24 min. Mobile phase A was DCM/acetonitrile (7/93; v/v and 2 mM ammonium acetate), and mobile phase B was water/acetonitrile (50/50; v/v and 2 mM ammonium acetate). The following linear gradient for B was applied: 0.1% B–50% B over 11 min, 50–70% B over 0.5 min, 70–100% B over 1 min, and held at 100% B for a further 2.5 min before returning to 0.1% B. At 2 min, the flow rate changed from 0.2 to 0.7 ml/min and remained at that flow rate until 15 min where it returned to 0.2 ml/min. Source conditions for positive mode were ionization voltage (IS) 5.5 kV, curtain gas (CUR) 30, temperature (TEM) 550°C, source gas 1 (GS1) 50, and source gas 2 (GS2) 60. Negative-mode source conditions were IS −4.5 kV, CUR 30 psi, TEM 550°C, GS1 50 psi, and GS2 60 psi. Dwell time was calculated in Analyst (V1.6, AB Sciex) automatically based on the number of multiple reaction monitoring (MRMs). This is a scheduled method with pos/neg switching throughout. PLs were quantified using an external calibration with internal standards as listed above. Here, they were quantified from standard curves containing two primary standards each (with the exception of Lyso PE and Lyso PG, which had one primary standard each) (SM C12, PC 16:0-18:1, PC 18;0-22:6, PE 16:0-18:1, PE 18:0-20:4, PG 16:0-18:1, PG 18:0-22:6, PI 16:0-18:1, PI 18:0-20:4, Lyso PC 16:0, Lyso PC 18:0, Lyso PE 16:0, and Lyso PG 16:0, Lyso PS 16:0 and 18:0). For confirming the absence of serum contamination of lipids in purified virus cultured from A549 cells, blank isolates (medium + 2% serum) were extracted and then analyzed using direct injection precursor scanning MS/MS for the presence of PE (precision 196, negative ion mode), PC (precision 184, positive ion mode), and CE (precision 369, positive ion mode), comparing with virus lipid extracts.

LC-MS/MS for free cholesterol and CEs and LC-MS analysis of TGs was performed on a Nexera liquid chromatography system (Shimadzu) coupled to either an API 4000 or 7500 qTrap mass spectrometer (Sciex). Liquid chromatography was performed at 40°C using a Hypersil Gold C18 (Thermo Fisher Scientific) reversed phase column (100 × 2.1 mm, 1.9 um) at a flow rate of 0.4 ml/min over 11 min. Mobile phase A was water/solvent B (95/5; v/v and 10 mM ammonium formate + 0.1% formic acid), and mobile phase B was acetonitrile/isopropanol (60/40; v/v and 10 mM ammonium formate + 0.1% Formic acid). The following linear gradient for B was applied: 90% for 1 min, 90–100% from 1 to 5 min and held at 100% for 3 min followed by 3 min at initial condition for column re-equilibration. Samples were spiked with cholesterol-d7 (1 ug), CE 18:1-d7 (100 ng), and TG 15:0/18:1-d7/15:0 (100 ng) prior to extraction. TGs were analyzed in selected ion monitoring positive mode, covering a range from TG 32:0 up to TG 56:0 including also unsaturated TGs. MS conditions were as follows: TEM 450°C, GS1 35 psi, GS2 50 psi, CUR 35 psi, IS 5 kV, declustering potential 60 V, and entrance potential 10 V for the 4000 QTRAP. 7500 conditions were as follows: TEM 150°C, GS1 70 psi, GS2 40 psi, CUR 44psi, IS 5.5 kV, and entrance potential 10 V. Dwell time was 10 ms. Triacylglycerides (TAGs) were quantified using an external calibration with TG 15:0/18:1-d7/15:0. Free cholesterol and CEs were analyzed in MRM mode monitoring the precursor to product transitions of 12 CEs and free cholesterol, as [M + NH4]^+^. MS conditions for the 4000 QTrap were as follows: TEM 150°C, GS1 25 psi, GS2 50 psi, CUR 35 psi, IS 5 kV, declustering potential 70 V, entrance potential 10 V, collision energy 20 V, and collision cell exit potential 25 V. 7500 conditions were as follows: TEM 150°C, GS1 70 psi, GS2 40 psi, CUR 44psi, IS 5.5 kV, and entrance potential 10 V Dwell time was 100 ms for each transition. Cholesterol and CEs were quantified using external calibration curves against the internal standards, with the following primary standards: cholesterol, CE 14:0, CE 16:0, CE 18:0, CE 18:1, CE 20:4, and CE 22:6. For all targeted assays, inclusion criteria for peaks were those at least 5:1 signal-to-noise ratio and with at least seven points across the peak.

### Identification and quantitation of external facing PE and PS on the surface of SARS-CoV-2

Total and external PE and PS were derivatized and analyzed using LC/MS/MS as described previously(*111*). Briefly, virus particles were suspended in 0.2 ml PBS and incubated with 20 μl of 20 mM NHS-biotin (total PE/PS) or 86 μl of 11 mM EZ-Link Sulfo-NHS-biotin (external PE/PS) for 10 min at room temperature before addition of 72 μl of 250 mM l-lysine. Volumes were increased to 0.4 ml using PBS. Vials containing 1.5 ml chloroform:methanol (1:2) solvent with 10 ng of internal standards (biotinylated 1,2-dimyristoyl-PE and 1,2-dimyristoyl-PS) were used for lipid extraction. The solvent:sample ratio was 3.75:1 as a modified Bligh/Dyer technique(*14*). Following vortexing and centrifugation (400 *g*, 5 min), lipids were recovered in the lower chloroform layer, dried under vacuum, and analyzed using LC-MS/MS. Samples were separated on an Ascentis C-18 5 μm 150 mm × 2.1 mm column (Sigma-Aldrich) with an isocratic solvent (methanol with 0.2% w/v ammonium acetate) at a flow rate of 400 μl/min. Products were analyzed in MRM mode on a Q-Trap 4000 instrument (Applied Biosystems, UK) by monitoring transitions from the biotinylated precursor mass (Q1 *m/z*) to product ion mass (Q3 *m/z*) in negative ion mode. The area under the curve for the analytes was integrated and normalized to internal standards. The ratio of external to total PE/PS was calculated for each molecular species and expressed as a fraction (%) externalized. MRM transitions monitored are provided in Supplementary Table 4.

### LC/MS/MS analysis of oxidized phospholipids

Lipid extracts were separated using reverse-phase HPLC on a Luna 3 µm C18 150 × 2 mm column (Phenomenex, Torrance, CA) with a gradient of 50 – 100 % B over 10 min followed by 30 min at 100 % B (A: methanol:acetonitrile:water, 60:20:20 with 1 mM ammonium acetate + 0.1% glacial acetic acid, B: methanol, 1 mM ammonium acetate + 0.1% glacial acetic acid) with a flow rate of 200 µl min^−1^. Lipids were analyzed in MRM mode on a 7500 Q-Trap (Sciex, Cheshire, United Kingdom), monitoring transitions from the precursor to product ion (dwell 20 ms) with TEM 550°C, GS1 60, GS2 70, CUR 40, IS − 3500 V, EP − 10 V, CE − 44 V and CXP at − 19 V. The peak area was integrated and normalized to the internal standard. For quantification of HETE-PEs, standard curves were generated with PE 18:0a/5-HETE, PE 18:0a/8-HETE, PE 18:0a/11-HETE, PE 18:0a/12-HETE and PE 18:0a/15-HETE synthesized as described previously(*112*). Information on MRM transitions and m/z values are presented in Supplementary Table 5. HETE-PEs were quantified using standard curves with DMPE used as internal standard, with LOQ at signal:noise 5:1. Due to the limited standards available, identifications for some lipids are putative, based on the presence of characteristic precursor and product ions, and retention times.

### Oxylipin method

Samples were spiked with 2.1-2.9 ng of 5-HETE-d8, 12-HETE-d8, 15-HETE-d8, 13-HODE-d4, 20-HETE-d6, TXB2-d4, PGE2-d4 and LTB4-d4 standards (Cayman Chemical). Lipids were separated by liquid chromatography (LC) using a gradient of 30-100% B over 20 minutes (A: Water:Mob B 95:5 + 0.1% Acetic Acid, B: Acetonitrile: Methanol – 80:15 + 0.1% Acetic Acid) on an Eclipse Plus C18 Column (Agilent), and analyzed on a Sciex QTRAP^®^ 7500 LC-MS/MS system. Source conditions: TEM 475°C, IS -2500, GS1 40, GS2 60, CUR 40. Lipids were detected using MRM monitoring with all parameters listed in Supplementary Table 6. Chromatographic peaks were integrated using Sciex OS 3.3.0 software (Sciex). Peaks were only selected when their intensity exceeded a 5:1 signal to noise ratio with at least 7 data points across the peak. The ratio of analyte peak areas to internal standard was taken and lipids quantified using a standard curve which was run at the same time as the samples. Supplementary Table 6.

### Chiral analysis of oxylipins

Lipid extracts (50 μl) were hydrolyzed to release free oxylipins from PL, as follows: following evaporation under N2, 1.5 ml isopropanol was added, the sample vortexed, then 1.5 ml 0.2 M NaOH (or water for controls) added before incubation at 60 °C for 30 min. Samples were acidified to pH 3.0 with 140 ml conc HCl (or water added to controls), 5 μl oxylipin internal standard mix added, and then oxylipins were extracted twice using 3 ml of hexane with 10 μl glacial acetic acid added to controls to aid extraction efficiency. The combined hexane layers were dried under N2 flow, resuspended in 100 μl of methanol, and stored under argon at 80 °C until analysis by LC/MS/MS as below. Lipids were separated using a Chiralpak IA-U column (50×3.0 mm, Diacel) in reverse phase mode, with flow rate 300 μl/min, at 40 °C, according to(*113*), on a 6500 Q-Trap (Sciex, Cheshire, United Kingdom). Mobile phase A was water:0.1 % acetic acid, and B was acetonitrile:0.1% acetic acid, and the gradient was 10 % B raised to 100 % B over 20 min followed by a 2 min hold then decrease to starting conditions over 2 min. MRM transitions and instrument parameters were as used for oxylipin reverse phase analysis. Control samples were first used to determine S/R ratios of free oxylipins, with their levels subtracted from those detected post-hydrolysis to specifically calculate the esterified pools.

### RNA extraction

Nucleic acids were extracted from saliva using a protocol adapted from the automated Bio-On-Magnetic-Beads (BOMB) COVID-19 protocol(*110*). A combination of manual pipetting and automation was used, with all automated steps done on a GEN2 OT-2 pipetting robot (Opentrons). 100 ul of saliva sample was manually added to a 96 deep-well-plate (Applied Biosystems), and 240 µl of magnetic bead solution added (Magnacell Ltd) with 270 µl guanidine isothiocyanate (GITC) based lysis buffer(*110*). To aid adsorption of nucleic acids onto the magnetic beads, the 96 deep-well-plate was placed onto an IKA MS 3 orbital shaker (Fisher Scientific) for 10 min at 2,000 rpm. The plate was centrifuged for 1 min at 3,000 rpm (Eppendorf 5920R centrifuge) and then placed on the Opentrons magnetic rack for 10 minutes to allow the beads to form pellets on the side of the wells. Using the Opentrons, the magnetic beads were then washed three times with 200 µl 80 % (v/v) ethanol, with nucleic acids adsorbed onto the magnetic bead surface eluted in 50 µl of RNAse-free water (Severn Biotech Ltd). To avoid transfer of any magnetic beads, the magnetic rack was used to separate the beads from the nucleic acid solution, with the nucleic acid solution subsequently transferred to a 96 well plates for downstream analysis or stored at -80°C for future use.

### Plaque assay

Cells were seeded in 12-well plates at approximately 1 × 10e5 cells per well and incubated for 24 hours where the monolayer was ∼ 80% confluent. Ten-fold dilution of clinical isolates were done on 48-well plates using DMEM. Prior to infection, cells were washed with 1 × phosphate buffered saline (PBS). Each dilution Virus inoculum in 250 μL was added to each well (in duplicates). Inoculum was removed after one hour of incubation. Viral titers were determined by Avicel plaque assay. Avicel Overlay was prepared by mixing 3% Avicel solution with an equal volume of 2 × DMEM and adding Gentamicin (50 ug/ml) +/- Amphotericin-B (2.5 ug/ml). Then 1 mL of the mixture were immediately added to each well. The plaque assay plates were incubated without disturbing at 37°C and 5% CO2. Overlays were removed on day three and cells were fixed with 1 ml of absolute methanol. After 5 min the methanol was removed, and cells were stained with 0.5 ml of a 0.2% crystal violet solution/well. After 3 days, cytopathic effect (CPE) of the clinical virus samples was identified visually. The positions of CPE foci were marked and counted with a permanent marker. Mean viral titers were then determined at the highest virus dilution that shows CPE as plaque forming unit (pfu)/ml.

### RT-qPCR analysis of SARS-CoV2

Reverse transcriptase-polymerase chain reaction (RT-PCR) was used for detection of three gene regions, SARS-CoV-2 nucleocapsid N1 and N2, and the small envelope protein (E) gene. The human control gene ribonuclease P (RPPH1) was used to identify viability of the samples tested and to evaluate efficiency of RNA extraction. We ran RT-PCRs in two duplex reactions: E gene and RNase P, and N1 and N2. All primers were unlabeled, with the E and N2 probes labelled with the hexachloro-fluorescein (HEX) fluorophore, with RNAse P and N1 probes labelled with the fluorescein amidite (FAM) fluorophore. We performed RT-qPCR using a Luna® Universal Probe One-Step RT-qPCR Kit (New England BioLabs) on a QuantStudio^TM^ 7 real-time PCR system (Applied Biosystems). Samples were run in a MicroAmp Endura optical 384 well clear plate (Applied Biosystems) with a reaction volume of 10 μl. The mastermix contained Luna® Universal One-Step Reaction Mix, 10 pmol of each primer, 5 pmol of each probe, Luna^®^ WarmStart^®^ RT Enzyme Mix, and molecular grade water. A total of 4 μl of sample was added to each well containing 6ul of mastermix. All plates were set up using a GEN2 OT-2 pipetting robot (Opentrons). Samples were tested against all four markers, with positive controls (heat-inactivated SARS-CoV-2) and no template controls (dH20) randomly placed within the plate. All primers and probes were ordered from Integrated DNA Technologies (IDT, Coralville, IA, USA). Primer and probe sequences of N1, N2, and RPPH1 were taken from(*114*), with primer and probe sequences of the E gene taken from(*115*). A complete list of primers, probes, and their respective sequences can be found in Supplementary Table 7.

### Virus production for macrophage experiments

For virus propagation, Caco-2 or Vero cells (DSMZ, Braunschweig, Germany) were seeded in T175 tissue culture flasks in minimal essential medium (MEM) supplemented with 10% fetal bovine serum (FBS) and containing 100 IU/ml penicillin and 100 ug/ml streptomycin and incubated at 37 °C with 5 % of CO2. Cell lines were then infected with HCoV-229E (ATCC no. CCL-137) or MERS-CoV (BEI Resources (EMC/2012, NR-44260), respectively. Cell culture supernatants were harvested after appearance of cytopathogenic effect (CPE), centrifuged to remove cell debris and cell free viral stocks were stored at -80°C. Virus titers were determined as TCID50/ml in 96-well microtiter plates in confluent cells.

### Viral infection of ALOX15B-silenced macrophages

Human peripheral blood mononuclear cells were isolated from commercially obtained buffy coats of anonymous donors (DRK-Blutspendedienst Baden-Württemberg-Hessen, Institut für Transfusionsmedizin und Immunhämatologie, Frankfurt, Germany) using Ficoll density centrifugation. Primary human macrophages were generated and transfected using siRNA as outlined here to achieve a knockdown of *ALOX15B*(*70*). Monolayers of control and ALOX15B KD macrophages were infected with either MERS-CoV at MOI (multiplicity of infection) of 1 or HCoV-229E at MOI of 0.1, respectively. Virus inoculum was removed after 2 hours, the cells were washed three times with PBS and supplemented with RPMI 1640 medium containing penicillin (100 U/ml), streptomycin (100 ug/ml) and 1% (v/v) FCS. Infection rates were determined by detection of intracellular viral RNA by qPCR at different time points. Primer sequences were used as follows: MERS-CoV UpE (Fwd: 5’-gca acg cgc gat tca gtt-3’, Rev: 5’-gcc tct aca cgg gac cca ta-3’)(*116*), MERS-CoV N2 (Fwd: 5’-ggc act gag gac cca cgt t-3’, Rev: 5’-ttg cga cat acc cat aaa agc a-3’)(*117*), HCoV-229E M (Fwd: 5’-ttc cga cgt gct cga act tt-3’, Rev: 5’-cca aca cgg ttg tga cag tga-3’)(*118*).

### Viral infection of ALOX15B-silenced macrophages with addition of sterol intermediates

To analyze the effects of sterol intermediates on post-entry processes, macrophages were infected with MERS-CoV as described in the section above. After 2 hours, virus inoculum was removed by three washes with PBS. RPMI 1640 medium containing penicillin (100 U/ml), streptomycin (100 ug/ml) and 1% (v/v) FCS containing either lipid solvent ethanol or sterol intermediates was added to the macrophages for the following 46 hours. Subsequently, RNA isolation was performed using TRIzol reagent (cat. no. L34355, Thermo Fisher Scientific). Sterols were used in the following concentrations: mevalonate (300 µM), geranylgeraniol, lathosterol, and desmosterol (each 15 µM). Lathosterol (cat. number Cay9003102) was purchased from Cayman Chemical, desmosterol (cat. number 700060P) from Avanti Polar Lipids, mevalonolactone (cat. number 68519), and geranylgeraniol (cat. number G3278) from Sigma-Aldrich.

### Accessible cholesterol measurements using streptolysin O

For live-cell imaging, naïve macrophages were incubated in serum-free media containing propidium iodide (PrI, 1 µg/ml) and vehicle (dithiothreitol (DTT), 20 µM) or γ-irradiated streptolysin O (SLO, 29 nM)(*119*). After one hour, media was aspirated and replaced with fresh serum-free media and red fluorescence was measured in cells using the Incucyte® S3 Live cell Analysis System (Essen BioScience, Ann Arbor, Michigan, USA). For flow cytometric PrI fluorescence measurements, siRNA transfected, or inhibitor treated cells were incubated with PrI, vehicle, or SLO as indicated above. Following treatment, PrI fluorescence in macrophages was measured by flow cytometry (FACSymphony™ A5 SE, BD Life Sciences, Heidelberg, Germany). In some experiments, macrophages were pretreated for 72 hours with water-soluble cholesterol (10 µg/ml). ML-351 (10 uM) was added for 24 hours prior to SLO treatment. In control cells, media was supplemented with DMSO (ML-351 solvent). ML-351 (cat. no. 6448) was purchased from Tocris (Wiesbaden-Nordenstadt, Germany), water-soluble cholesterol (cat. no. C4951), γ-irradiated streptolysin O from Streptococcus pyogenes (cat. no. S0149) and propidium iodide (cat. no. P4170) from Sigma Aldrich.

### Statistics

For in vitro comparisons, one-way ANOVA with Tukey Post Hoc test was used where there were 3 or more separate conditions (https://astatsa.com/OneWay_Anova_with_TukeyHSD/), or unpaired Student t-test (Excel) where there were only 2 comparisons. Where a lipid was below LOQ in 50 % or more of samples, the lipid was removed from analysis in that experiment. Where any individual sample had over 50 % of the lipids below LOQ, the sample was removed from analysis in that experiment (this issue only impacted the 4-strain comparison experiment). Supplementary Data provides red entries to denote imputed values. Although all in vitro experiments were conducted using n = 3 biological replicates, in the 4-strain comparison, one isolate of England2 had a low yield for several lipid categories, and another from the Beta strain had low yield for some lipids, due to low recovery of virus. For this experiment, each “n” represents a different culture of virus, obtained around 1 week apart using different inoculations of stock virus, and due to this, overall yields showed higher variability. For all other *in vitro* experiments (IL-6, IL-4, Dex), samples were established at the same time. In all experiments, missing values were replaced with 20 % of the lowest detected value (ng/sample). Comparison across lipid categories was done following conversion of lipidomics data to mol/sample using an average mass value for each lipid category. Comparison within lipid categories instead used ng/sample data. Where n = 3, mean +/- SEM is shown on bar charts. Where n = 2 for any samples in a comparison (applies to 4 strain comparison only), SD is shown instead for all bars, even though most samples in the experiment will have had n = 3. Figure legends provide specific information on each experiment and number of replicates analyzed.

## SUPPLEMENTARY FIGURE LEGENDS

**Supplementary Figure 1.**
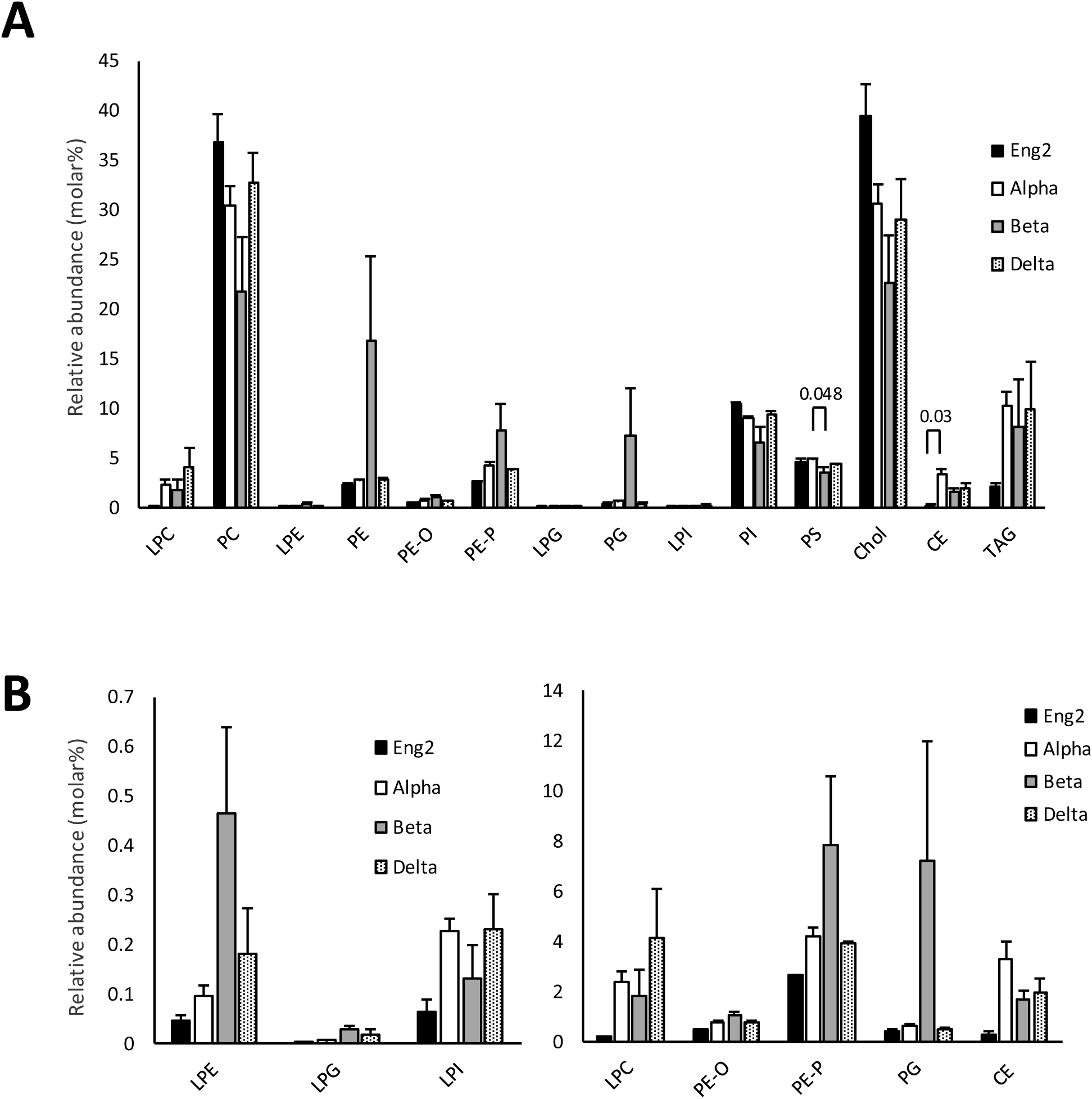
Lipidomics of four SARS-CoV-2 pandemic strain viruses reveals their overall composition to be relatively conserved. Lab grown virus from AAT cells was isolated using density gradient centrifugation, lipids extracted and then analyzed using targeted LC/MS/MS as outlined in Methods. Each isolate was from a different culture obtained at least one week apart. Alpha, Beta, Delta (n=3 isolates mean +/- SEM), England2 (n=2 isolates, mean +/-SD). Relative abundance was calculated from ng values of all lipid species totaled, converted to molar amount, then related to a mean mass value for each category. Statistics used one way ANOVA with Tukey Post Hoc test. Panel B shows zoomed in data for lower abundance species shown in A.

**Supplementary Figure 2.**
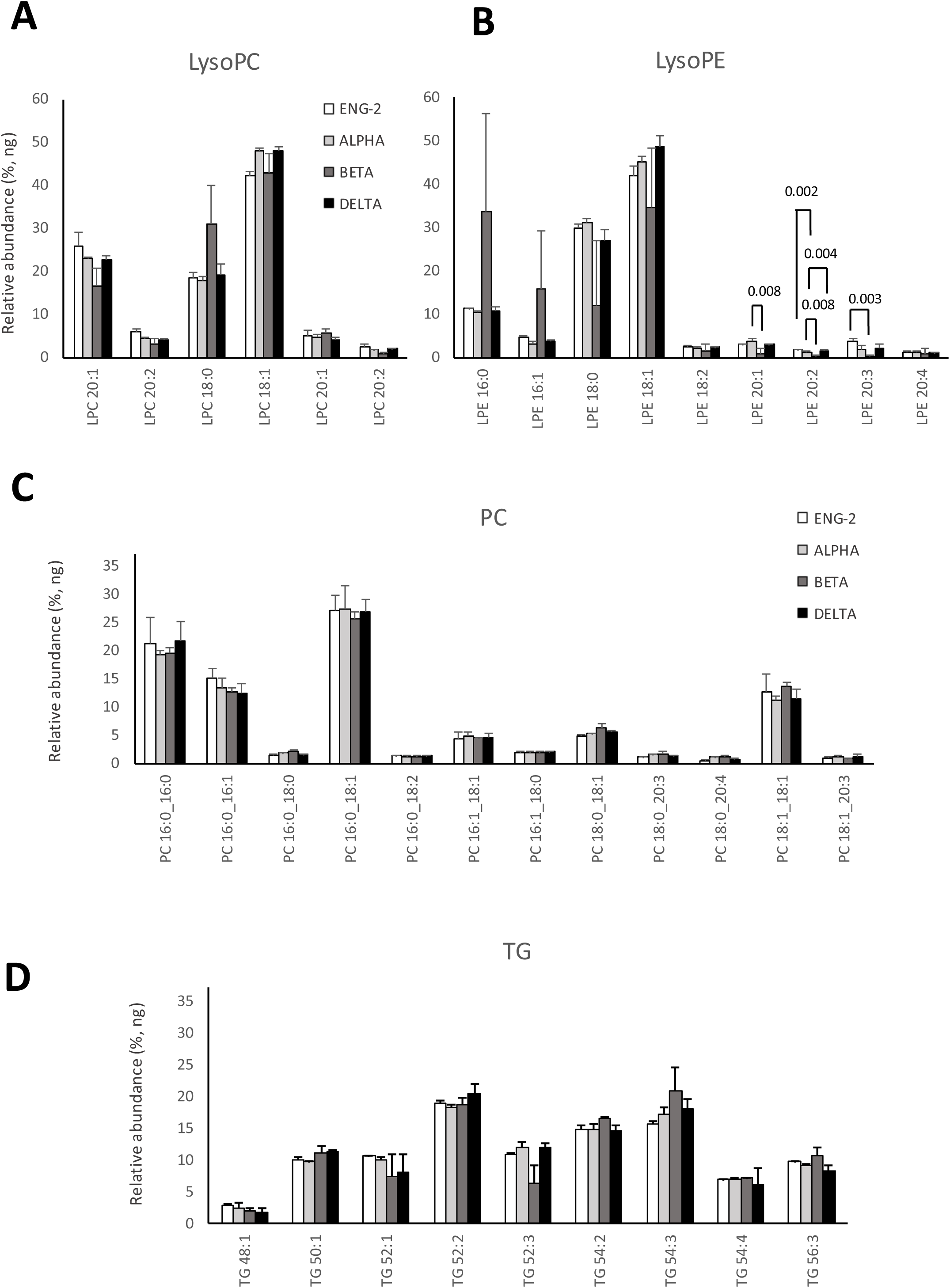
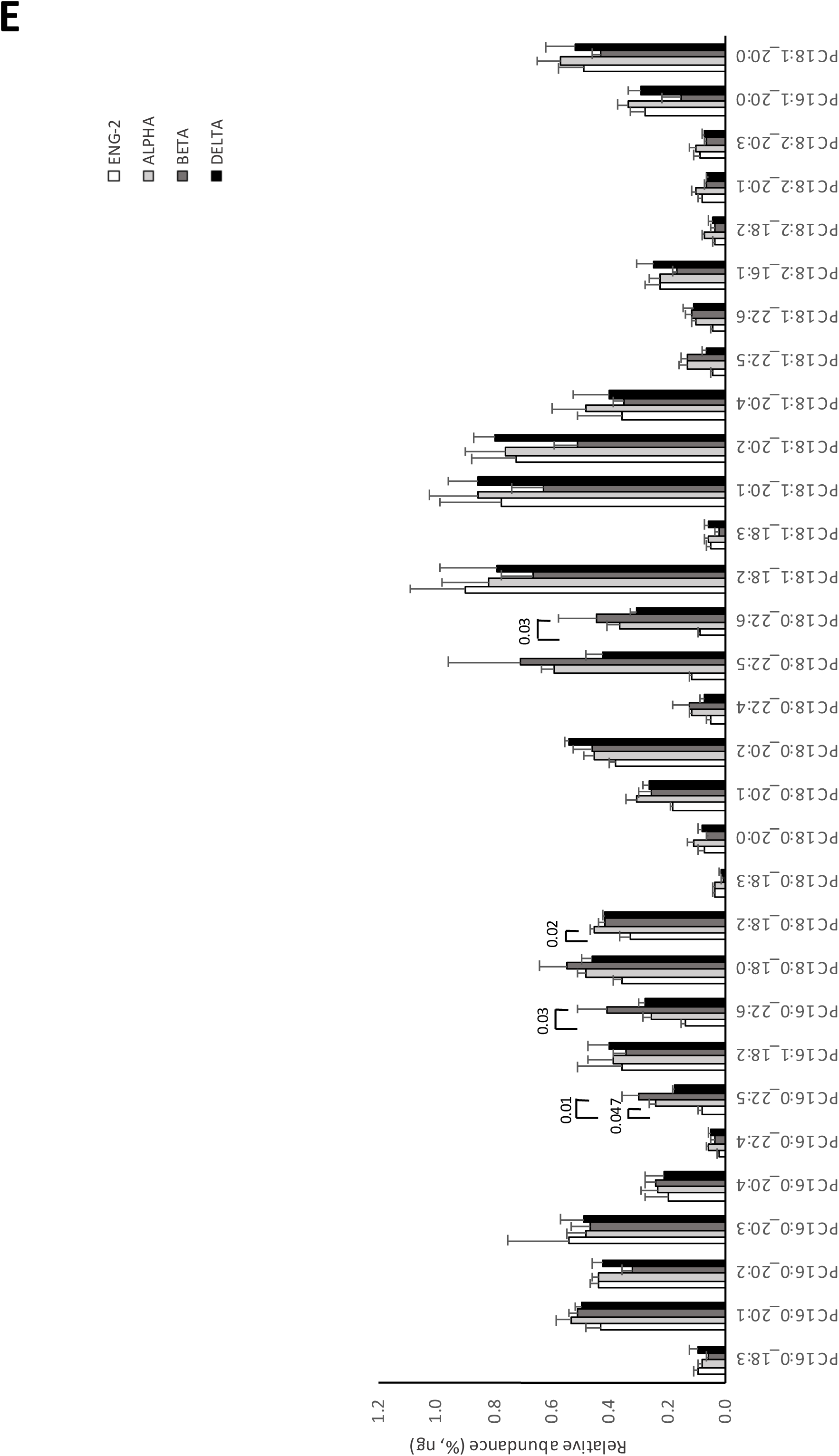
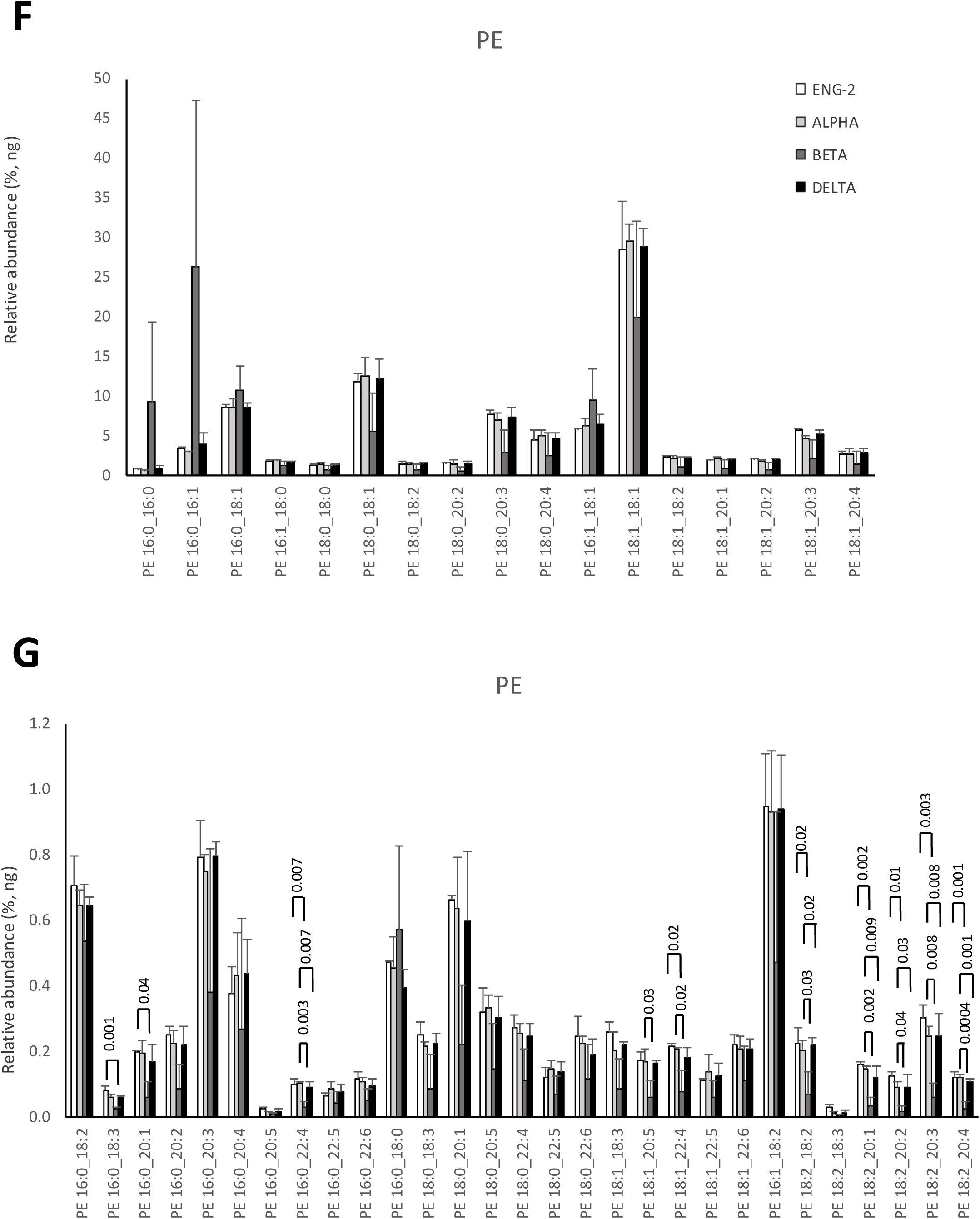
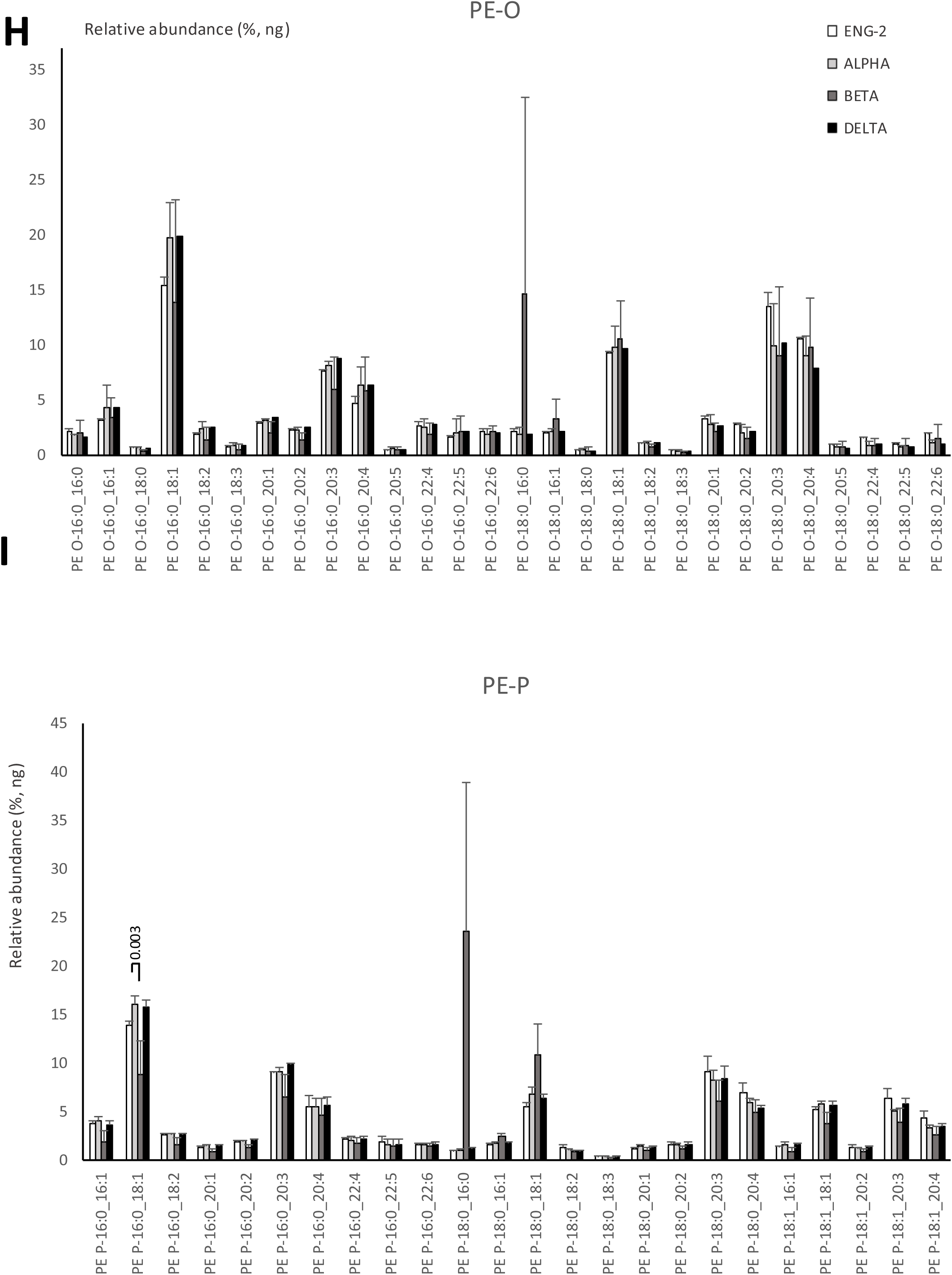
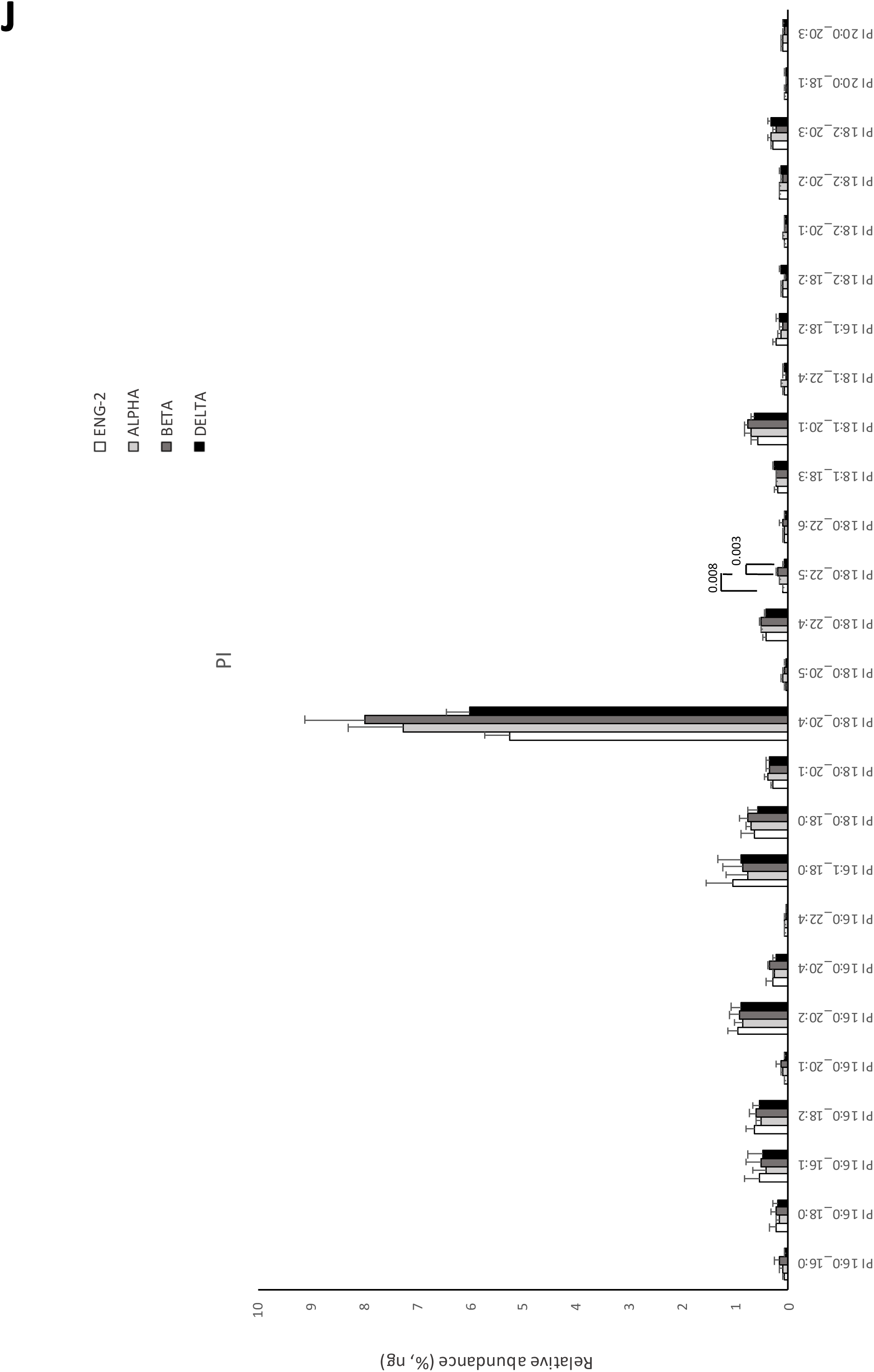
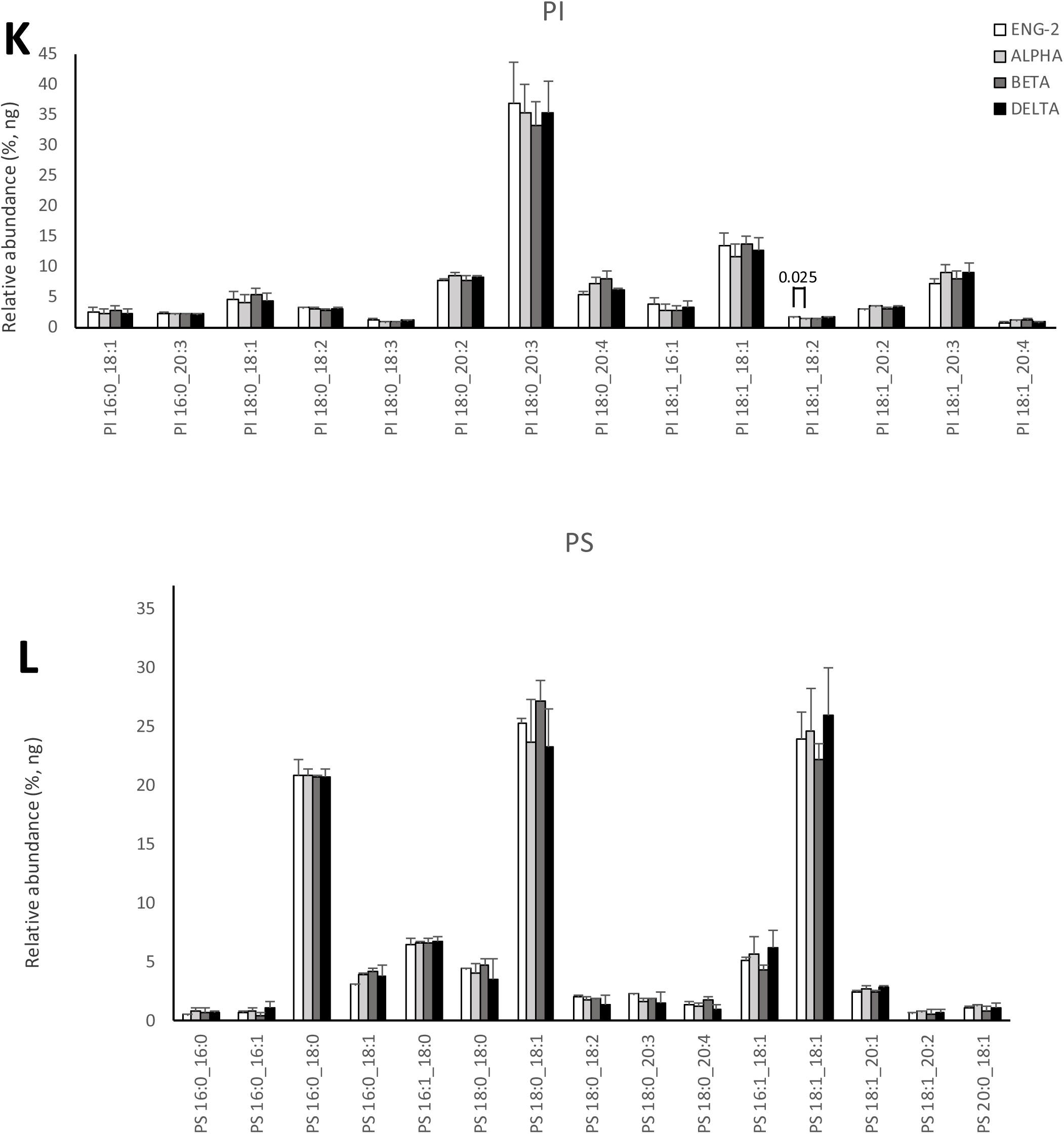
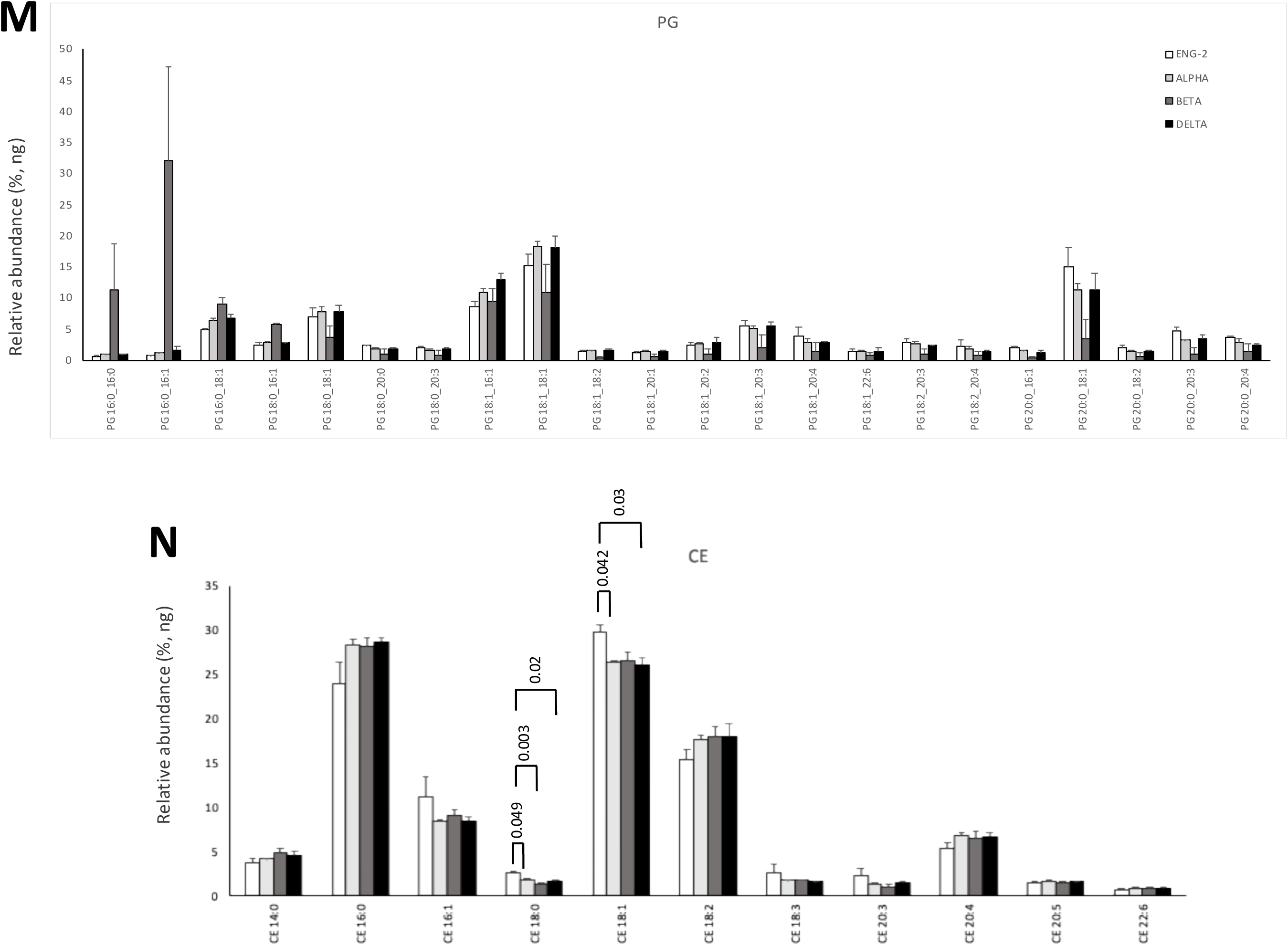
Lipidomics of four SARS-CoV-2 pandemic strain viruses reveals their molecular species to be relatively conserved. Lab grown virus from AAT cells was isolated using density gradient centrifugation, lipids extracted and then analyzed using targeted LC/MS/MS as outlined in Methods. Each isolate was from a different culture obtained at least one week apart. All panels are n=3 for all strains except for lysoPE, PE, PE-O and PS, which have n=2 for England2 only. TG is n=3 for Alpha and Delta and n=2 for England2 and Beta strain. Where all samples have n=3 isolates, mean +/- SEM is shown. Where any have n=2, mean +/- SD is shown. Relative abundance was calculated from ng values of all lipid species totaled. Statistics used one way ANOVA with Tukey Post Hoc test.

**Supplementary Figure 3.**
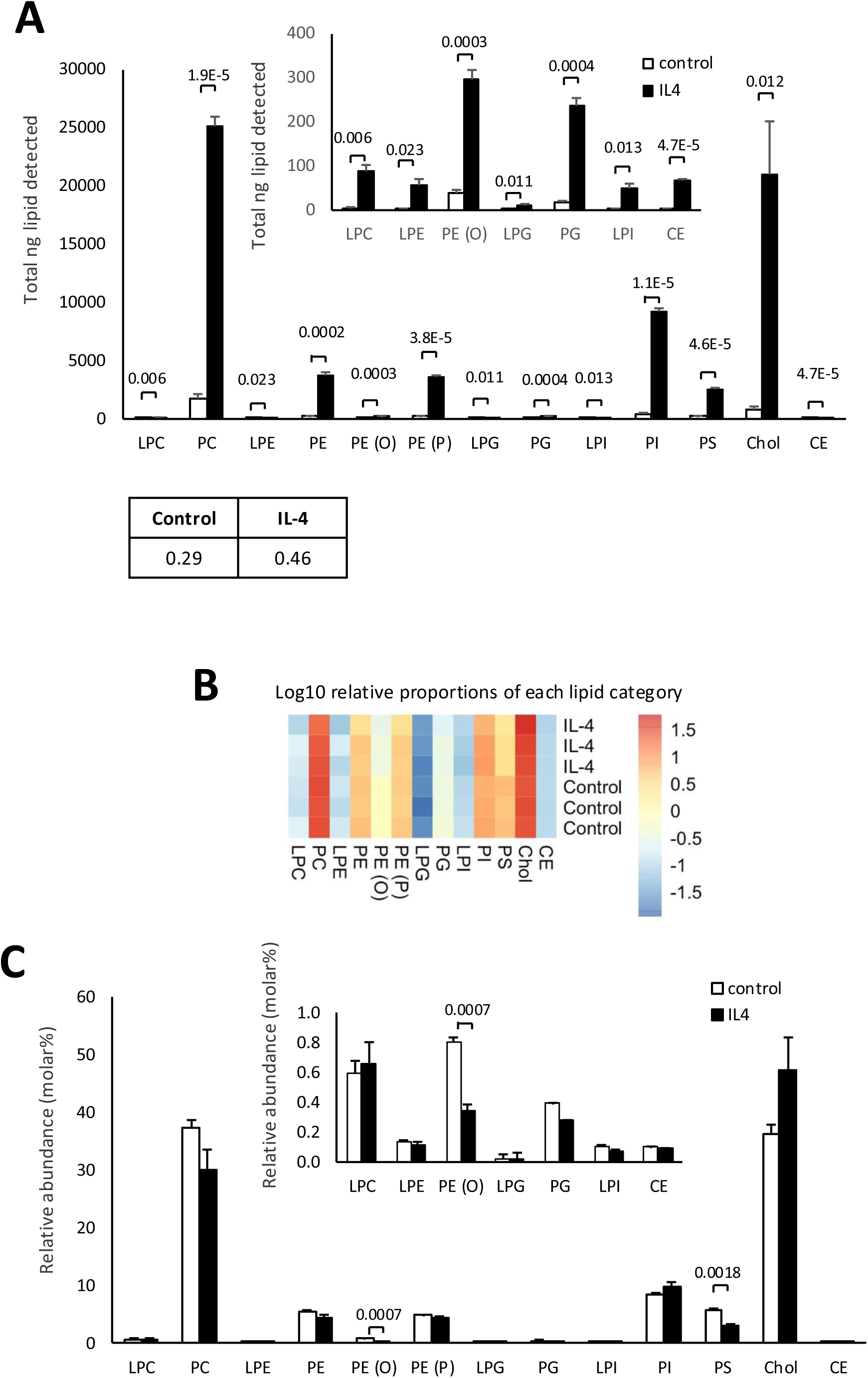
Lipid amount, but not composition of virus particles is significantly increased by IL-4 treatment. *Panel A. Total virus lipid recovered is significantly increased from IL-4 treated host cells.* IL-4 AAT cells were pretreated with IL-4 for 5 days as outlined in Methods, before infection with SARS-CoV-2 England2 strain. Virus was harvested using density gradient centrifugation and lipids extracted and quantified using LC/MS/MS as outlined in methods. Heatmap shows ng values as total for each PL category, n=3 isolates mean +/- SEM, Statistics used one way ANOVA with Tukey Post Hoc test. *Panel B. IL-4 treatment leads to little change in overall lipid composition of shed particles.* Data from Panel A was expressed as relative molar proportions, by normalizing to an average mass value per category and shown as log10 in a heatmap (Panel B), or as a bar chart (Panel C), n=3 isolates mean +/- SEM, Statistics used one way ANOVA with Tukey Post Hoc test.

**Supplementary Figure 4.**
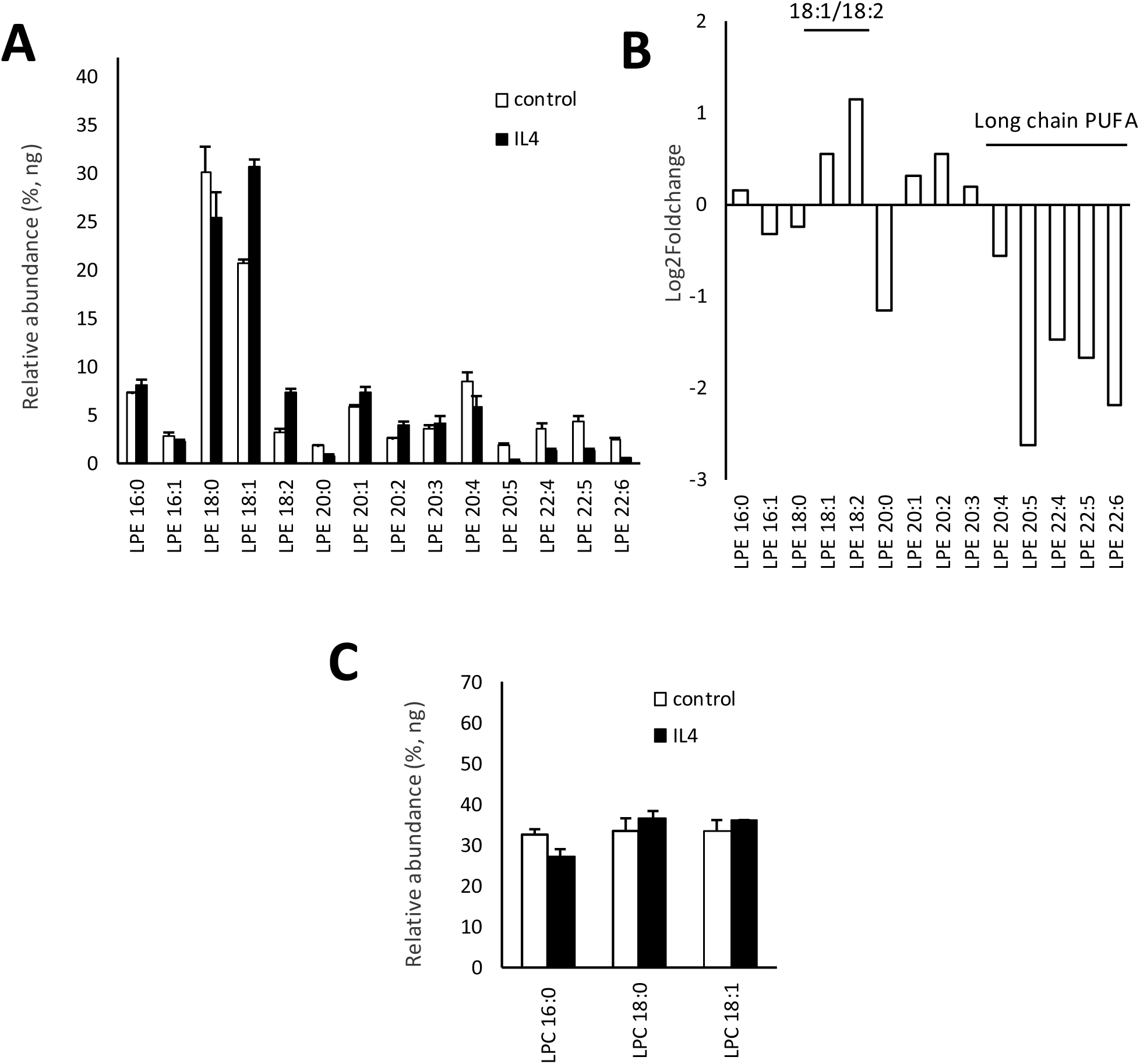
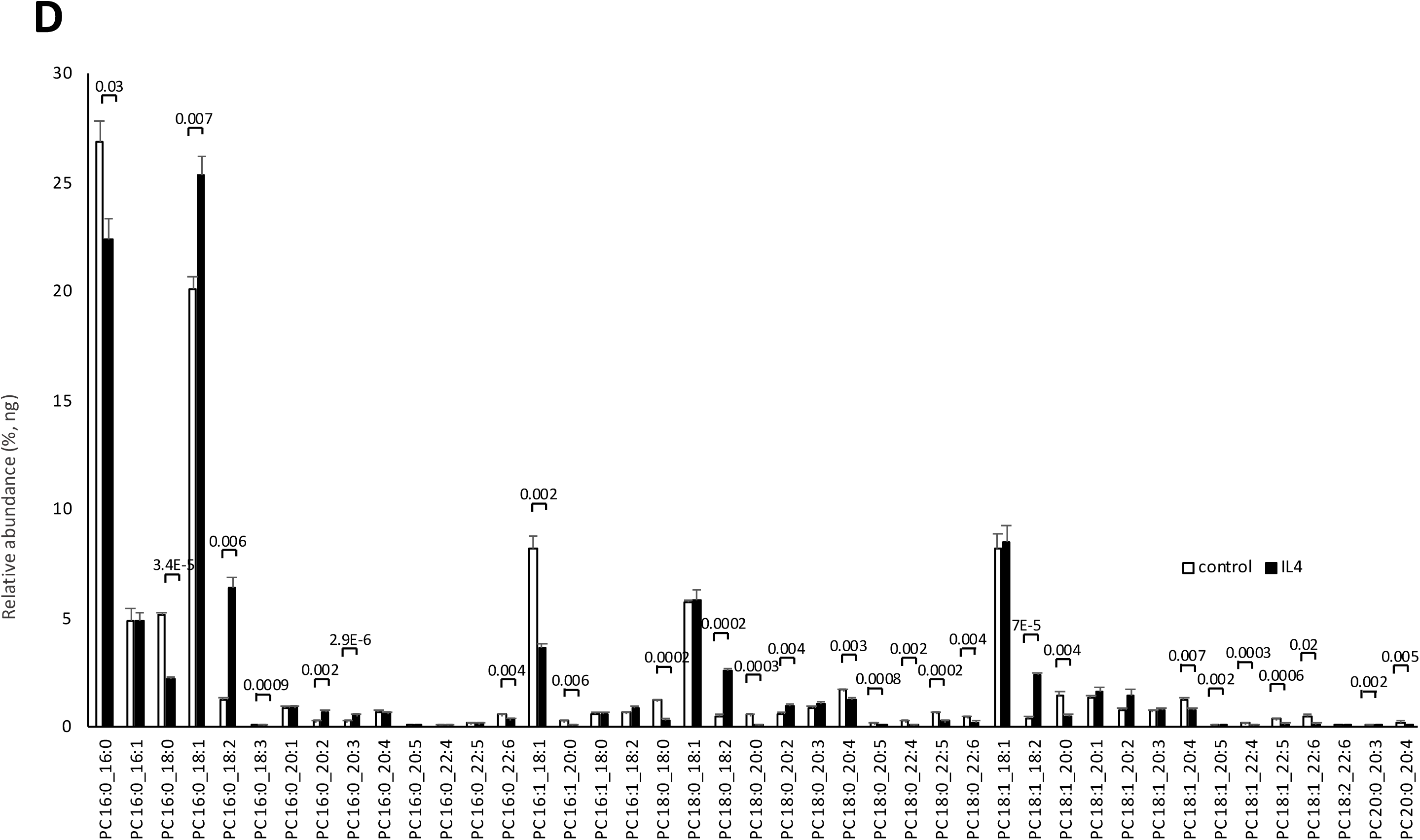
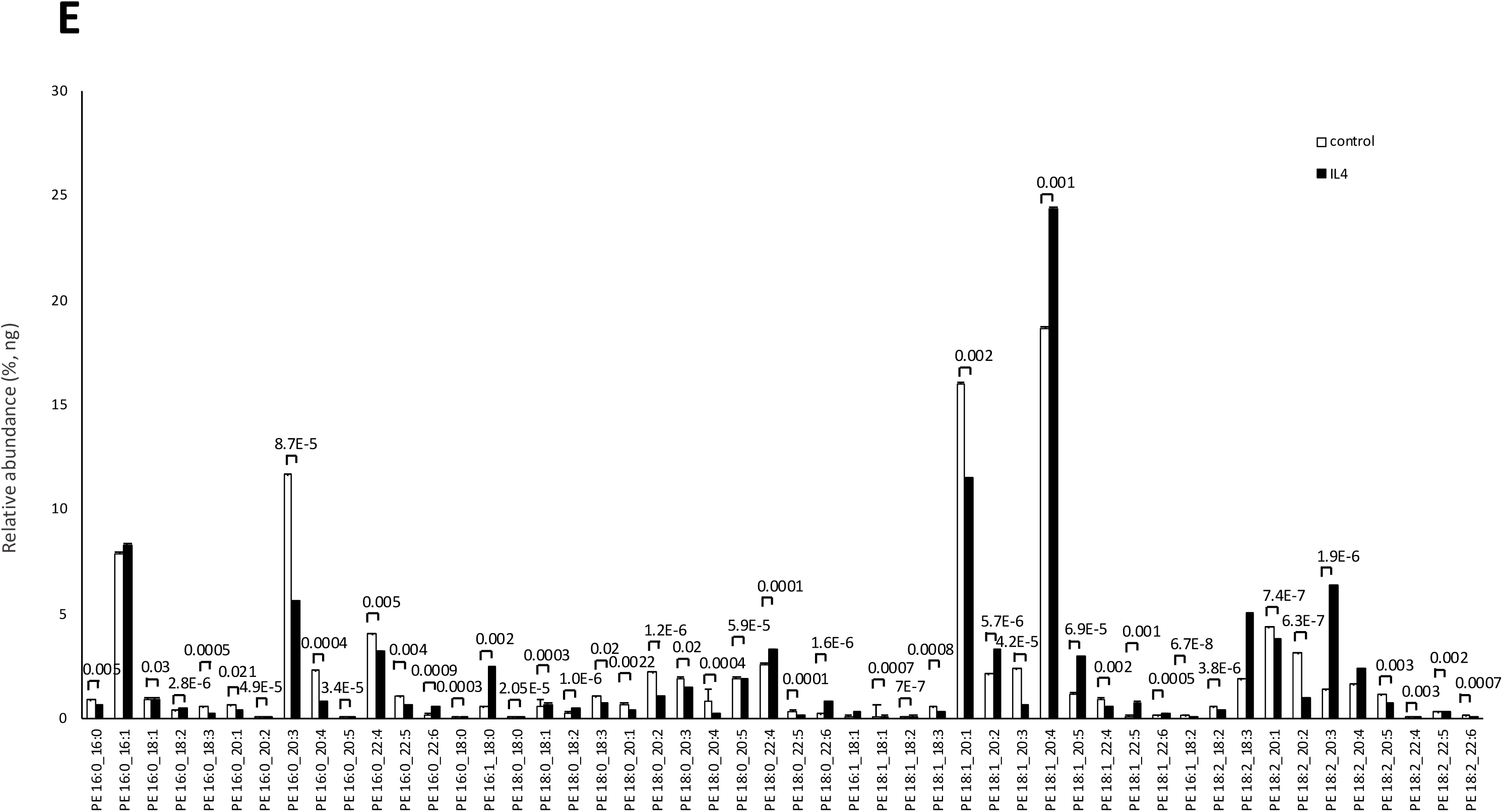
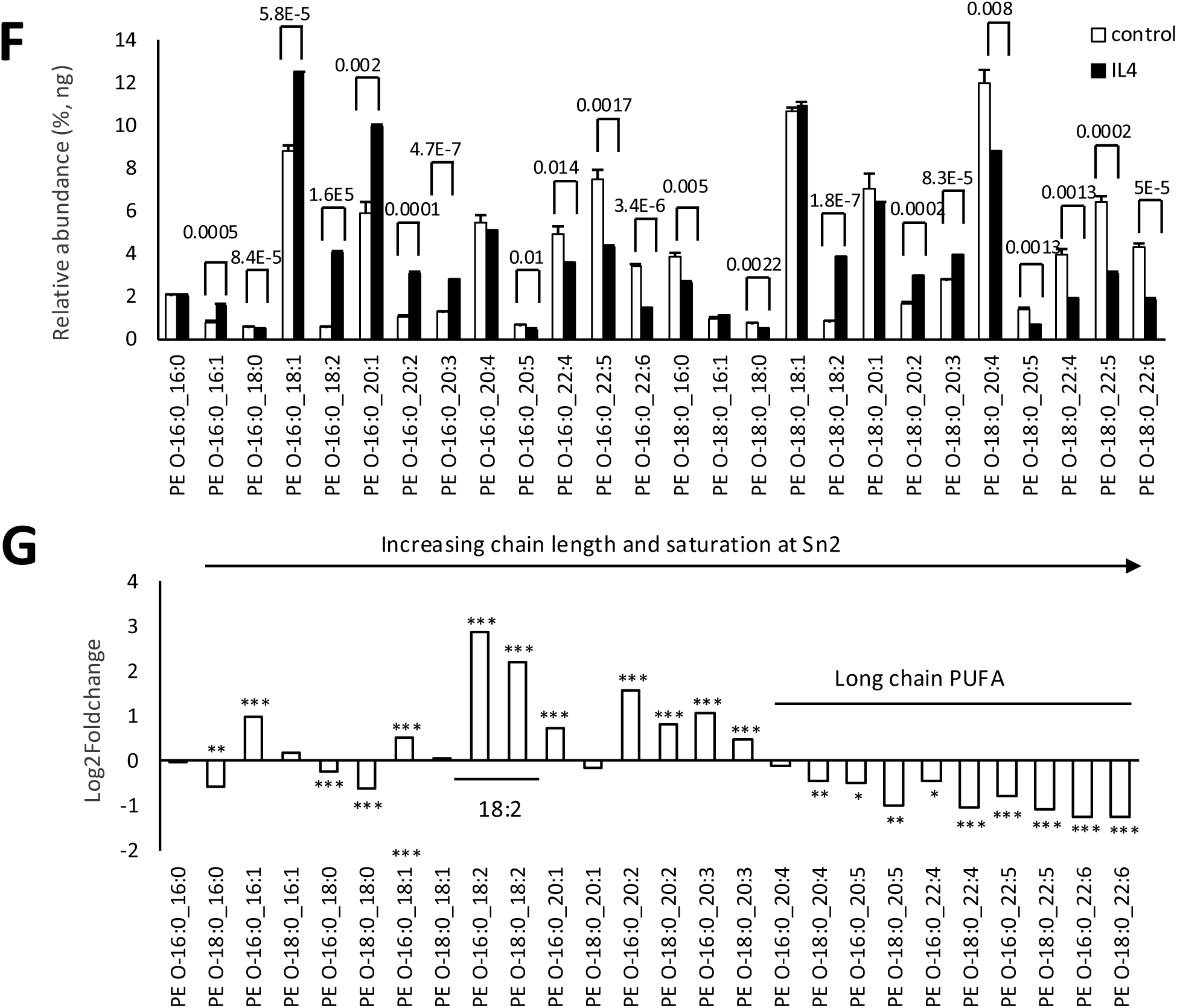
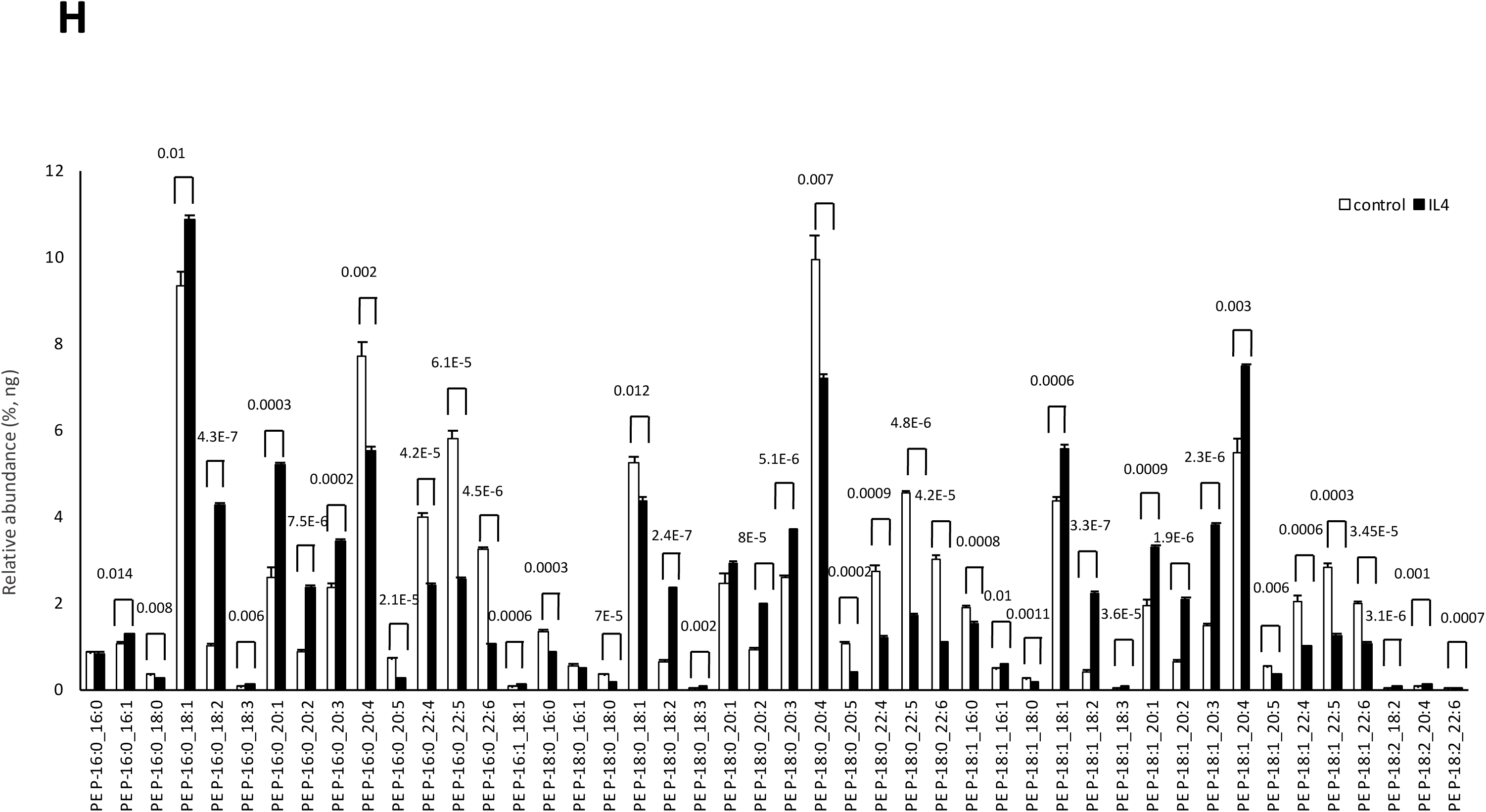
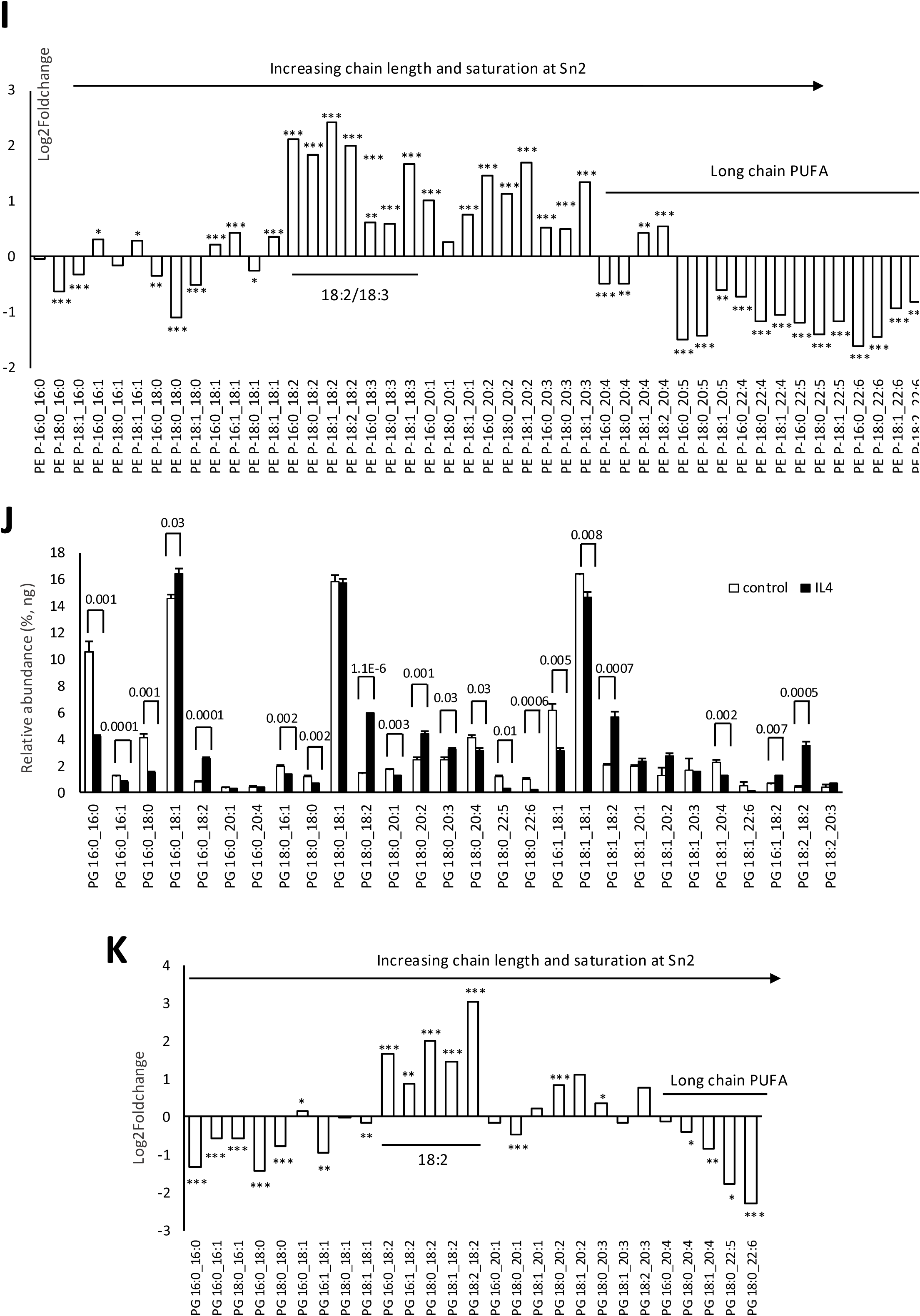
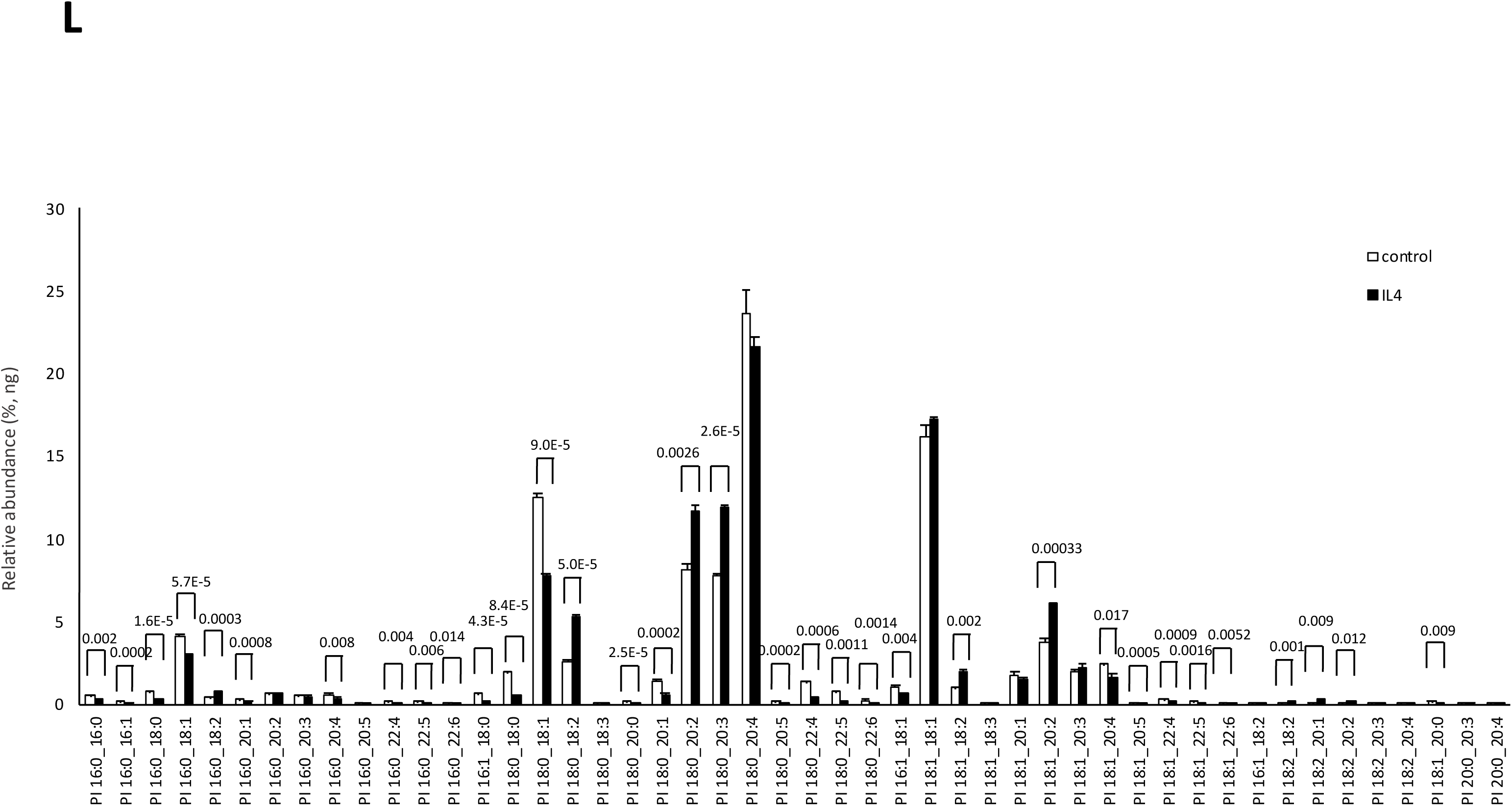
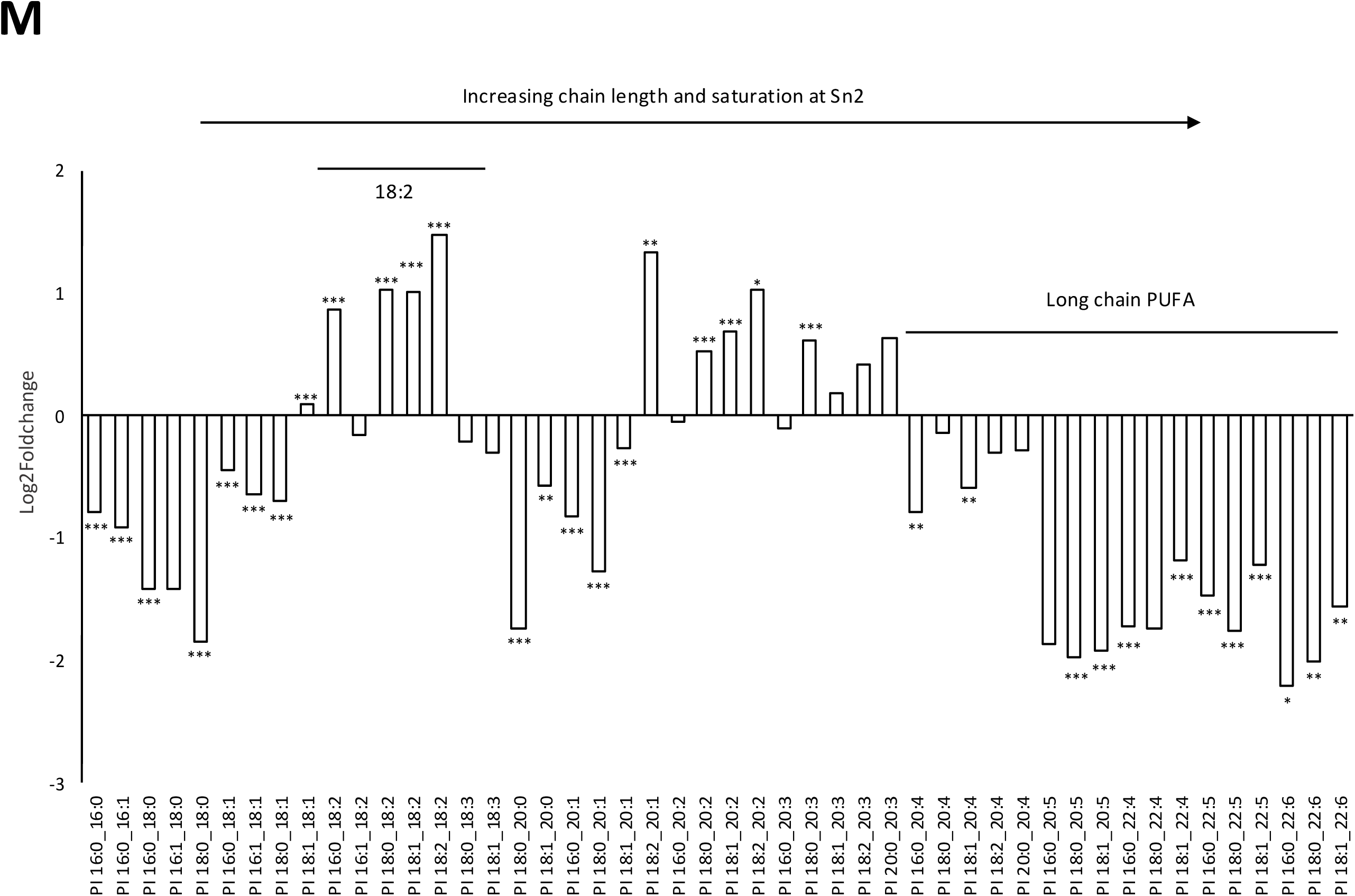
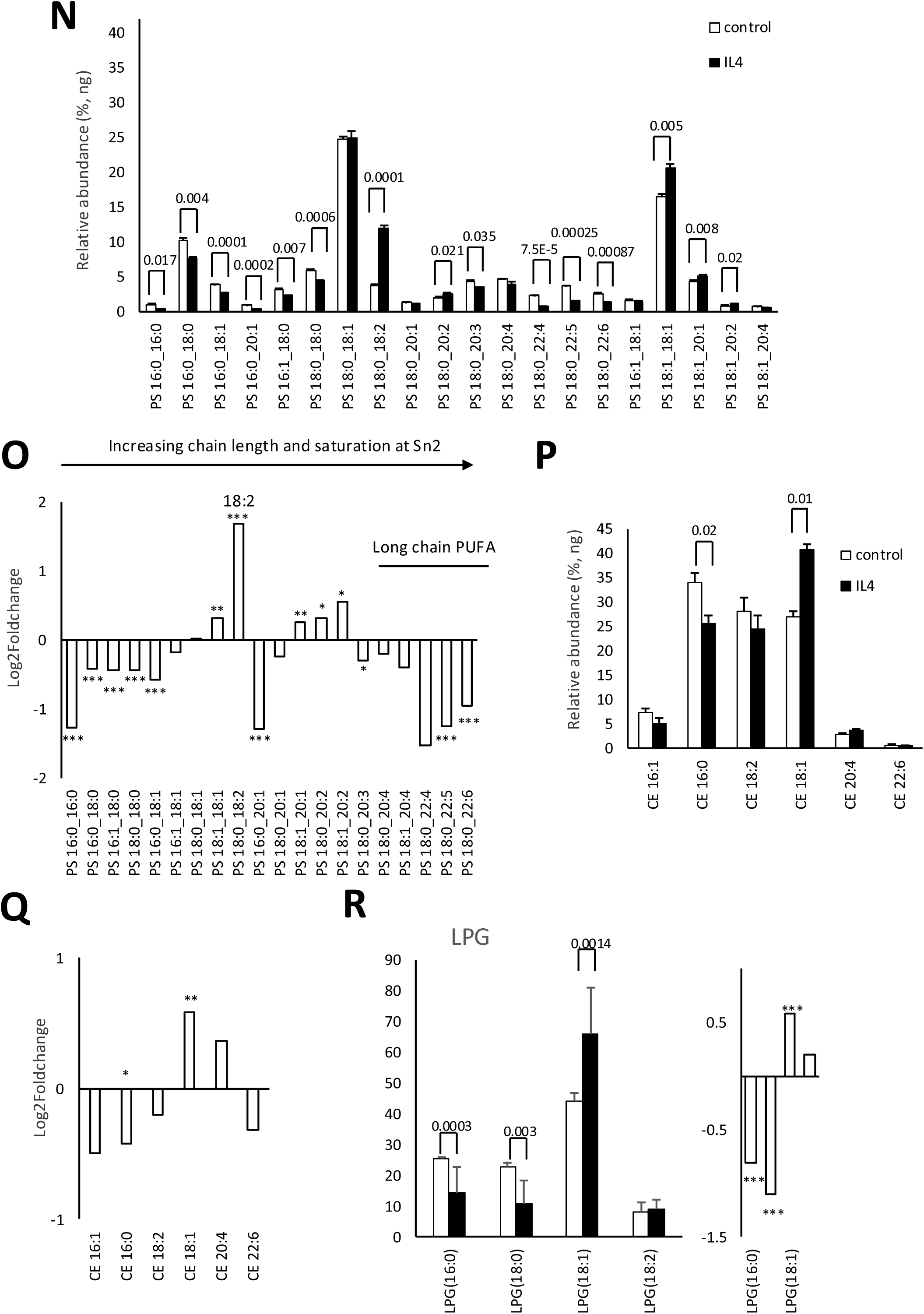
IL-4 pretreatment of host AAT cells leads to reduced PUFA and saturated FA containing PL, but increased PL with 18:2/18:3. AAT cells were pretreated with IL-4 for 5 days as outlined in Methods, before infection with SARS-CoV-2 England2 strain. Virus was harvested using density gradient centrifugation and lipids extracted and quantified using LC/MS/MS as outlined in methods (n=3 isolates, mean +/- SEM, unpaired Student t-test). Amounts of individual PL species detected were averaged and compared for an impact of IL-4 treatment, and are shown as either bar charts, or log2foldchange. A decrease means reduction in the species by IL-4, (n=3 isolates, mean +/- SEM), unpaired Student t-test, * P<0.05, ** P<0.01, ***P<0.005.

**Supplementary Figure 5.**
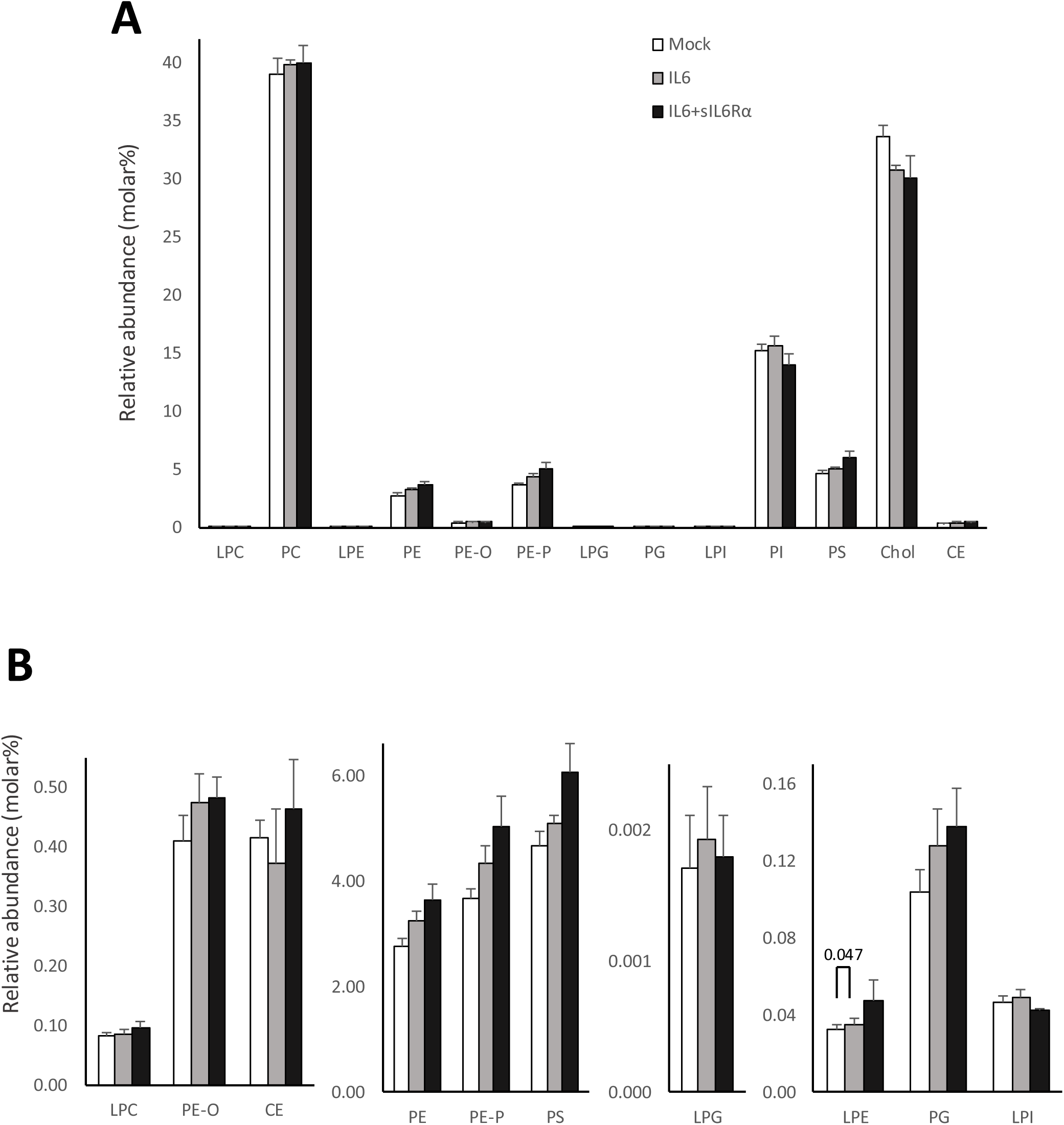
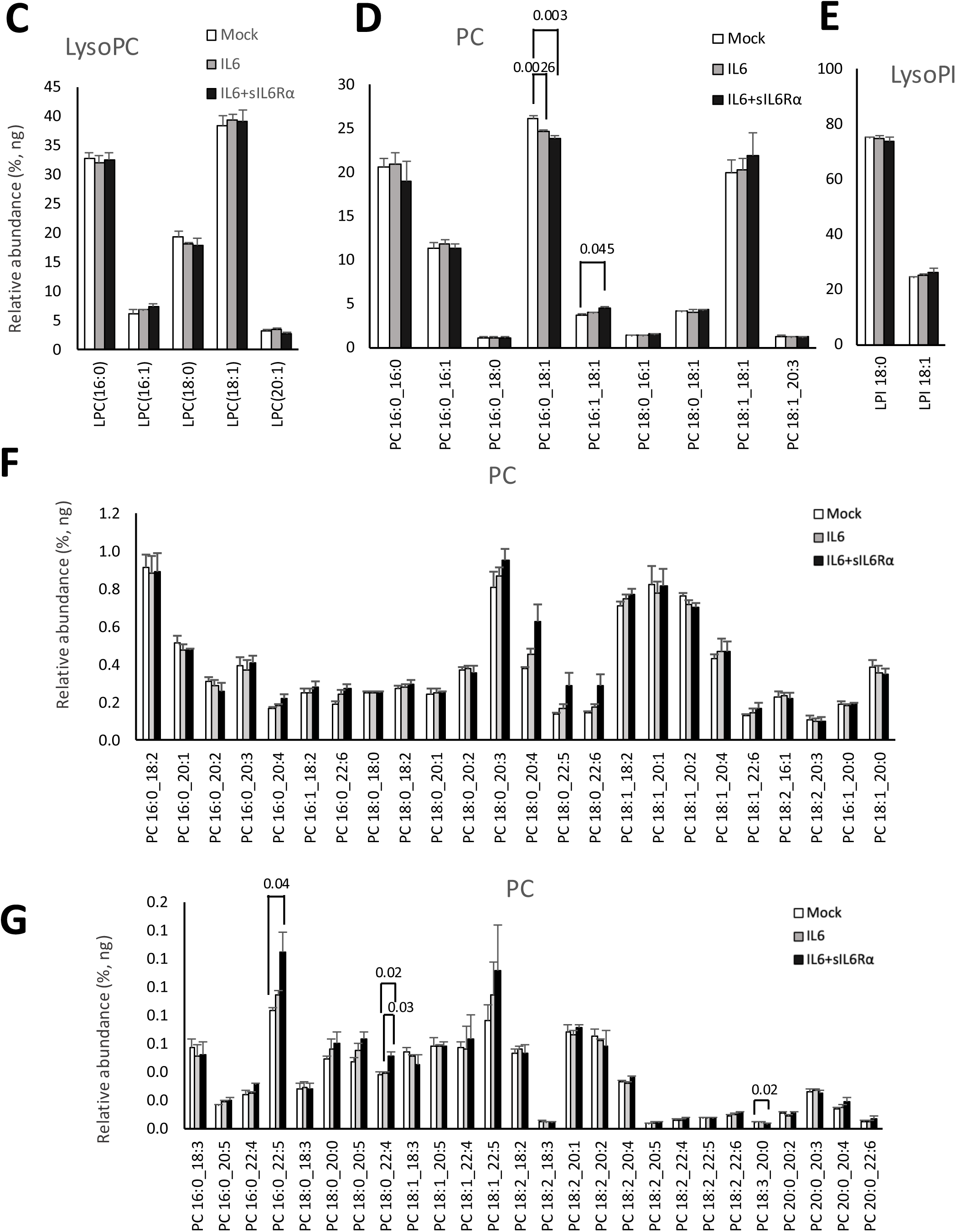
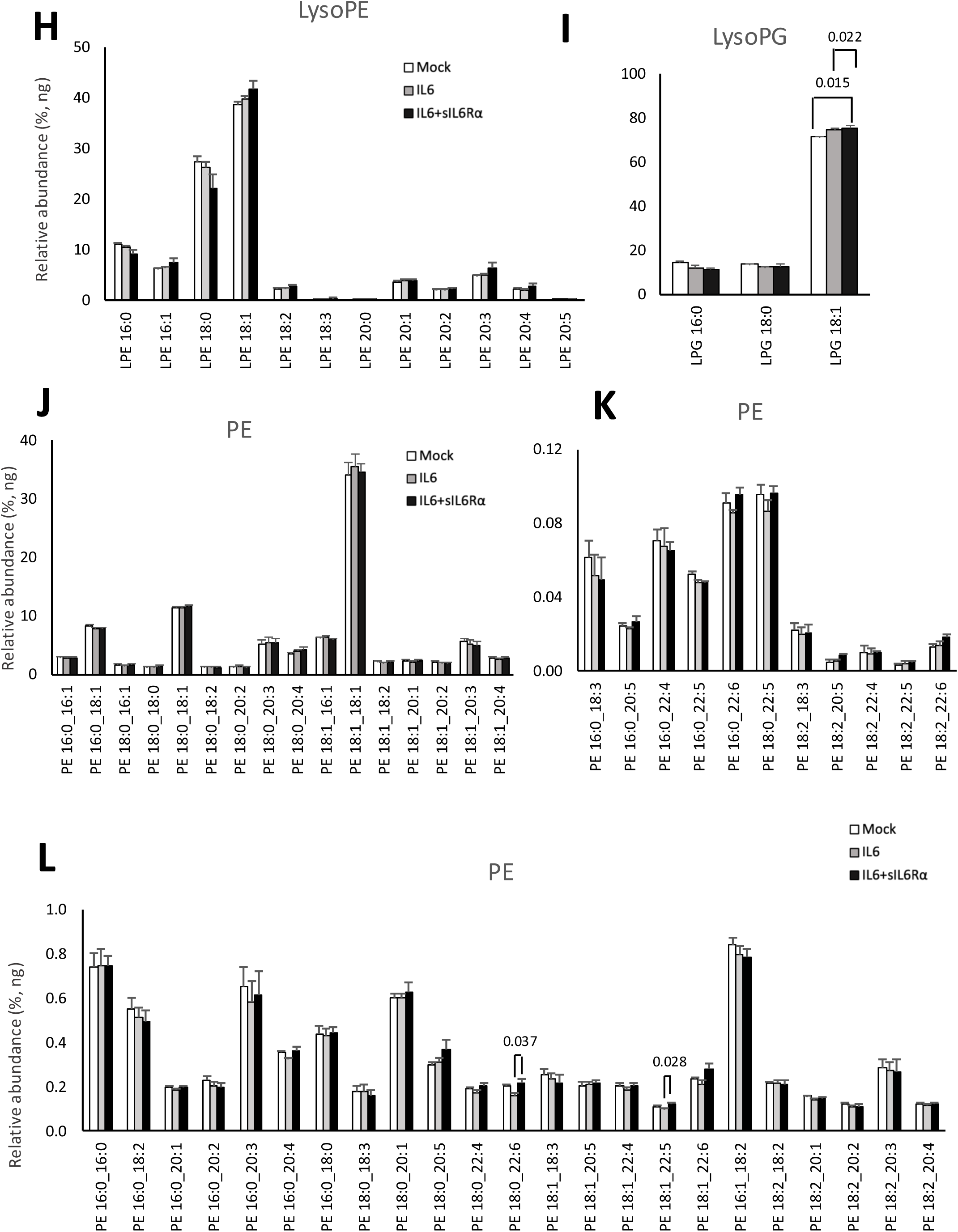
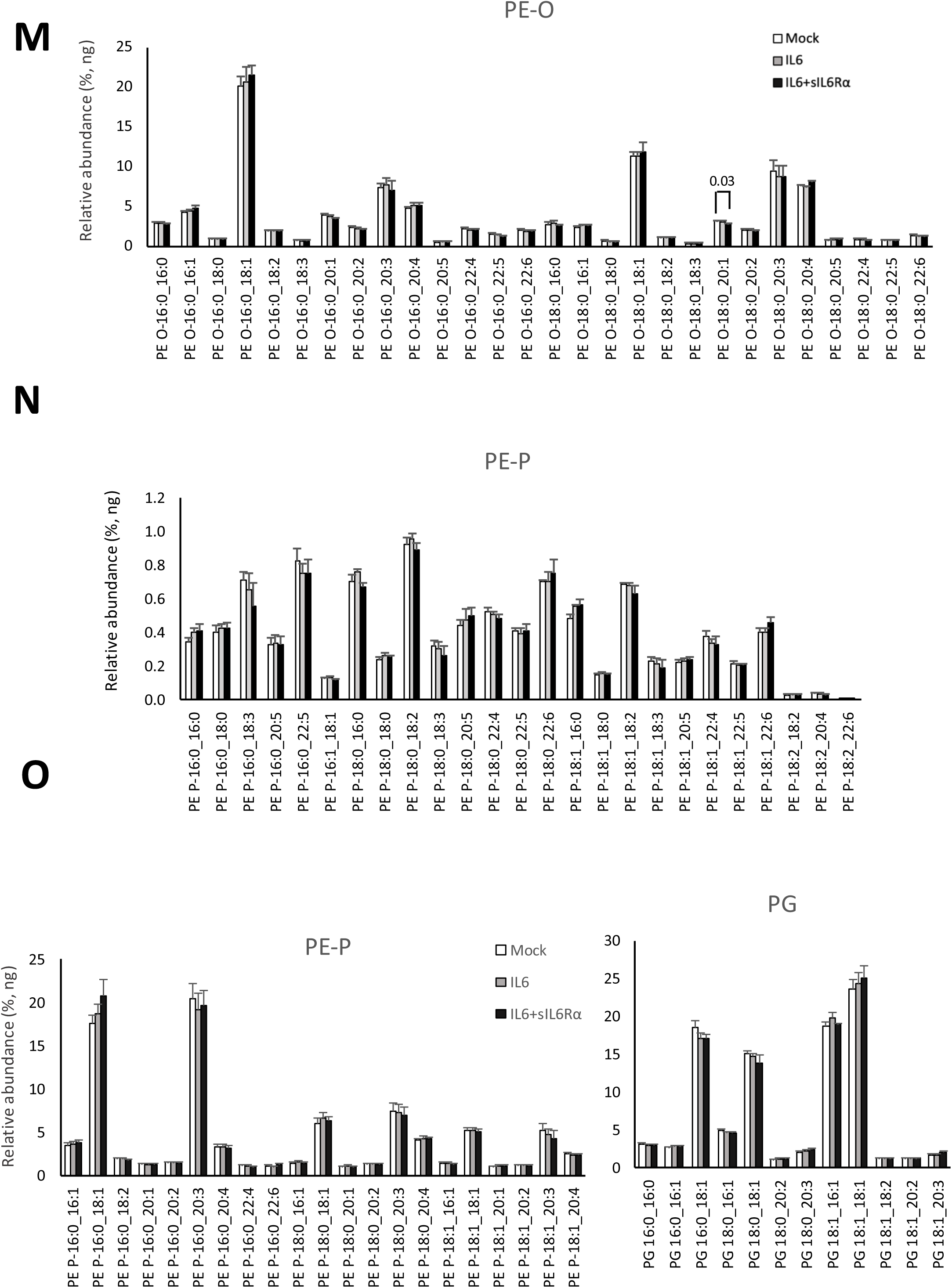
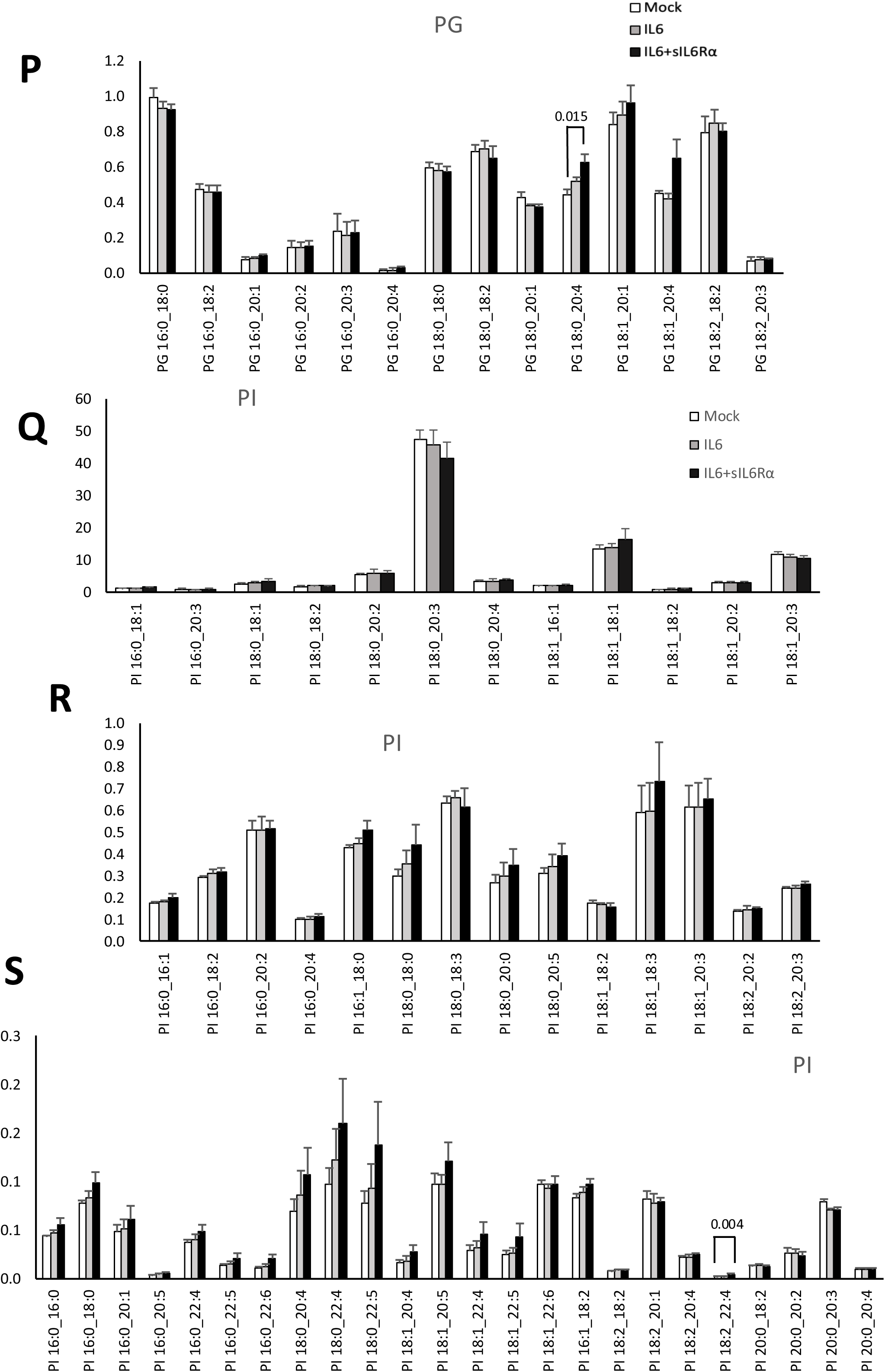
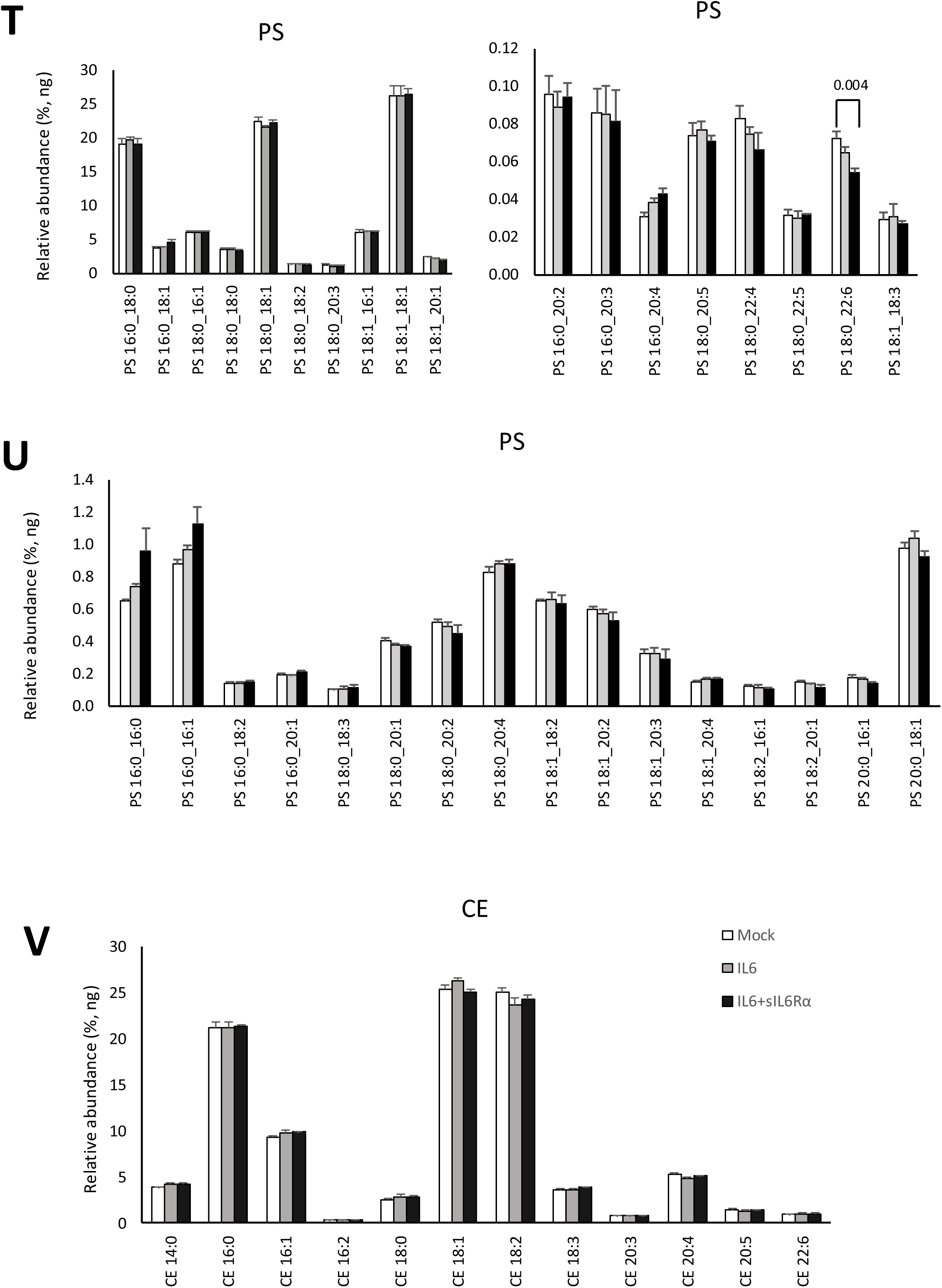
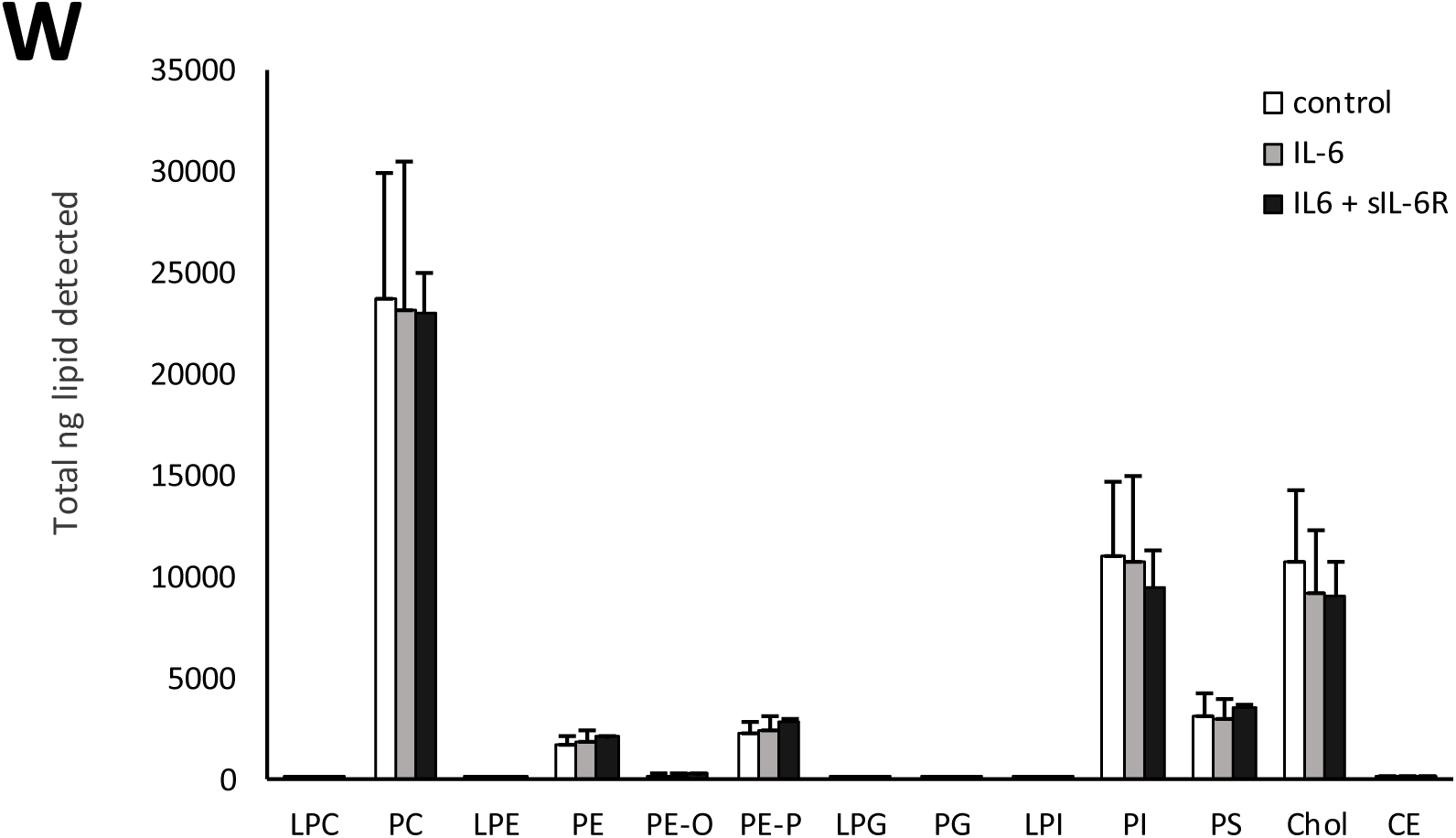
IL-6 treatment of host cells has no impact on SARS-CoV-2 membrane composition. *Panels A,B.* Lipids were extracted from virus from IL-6 or IL-6/sIL-6Rα-treated AAT cells then analyzed using LC/MS/MS as outlined in methods. Lipid amounts are expressed as relative molar%, by totaling all species then normalizing to an average mass value per category (n=3 isolates, mean +/- SEM), unpaired Student t-test. *Panels C-W.* Lipids were extracted from virus from AAT cells treated with IL-6 or IL-6/sIL-6Rα, then analyzed using LC/MS/MS as outlined in methods. Lipid amounts are expressed as relative abundance, ng% (n=3 isolates, mean +/- SEM), unpaired Student t-test.

**Supplementary Figure 6.**
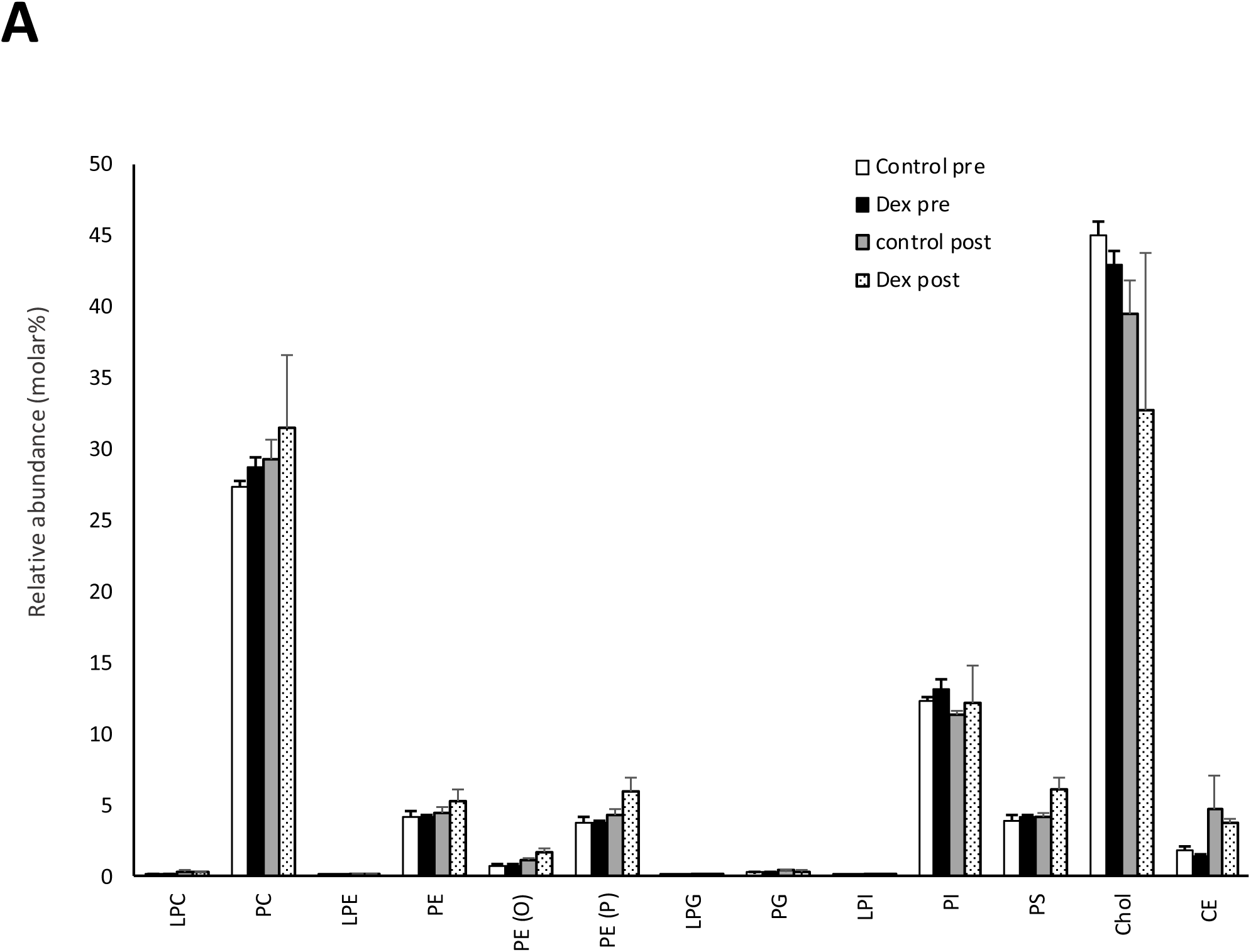
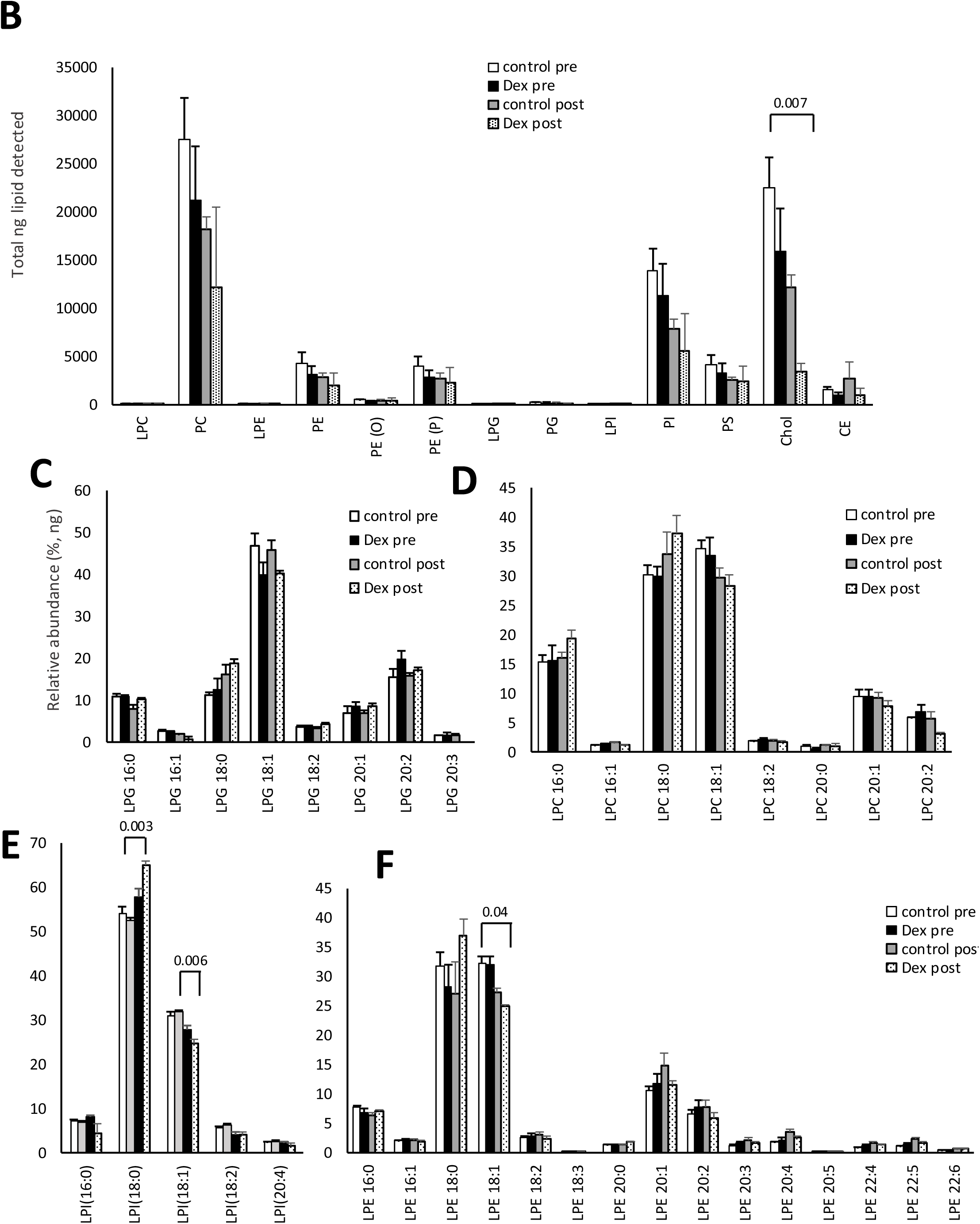
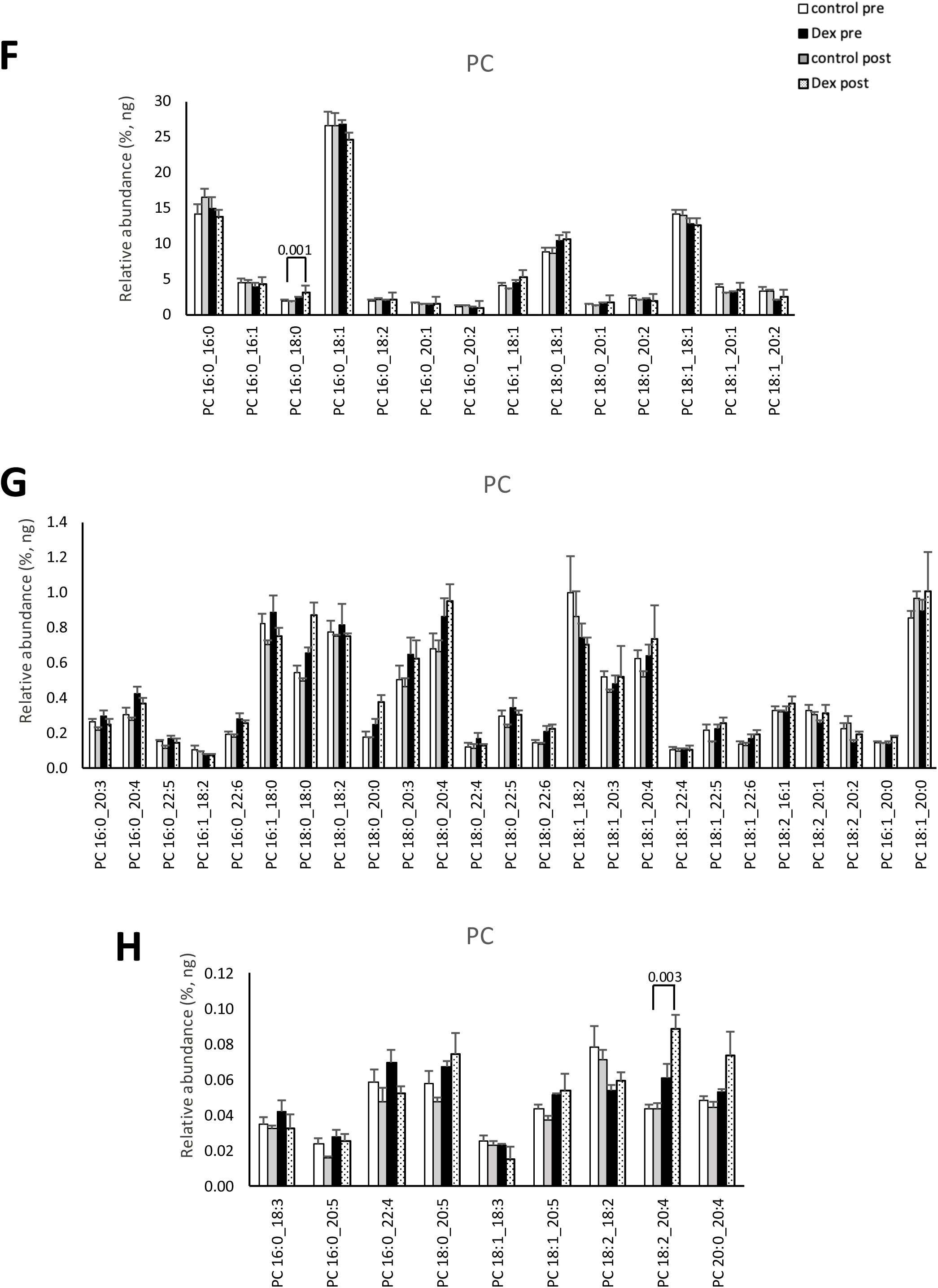
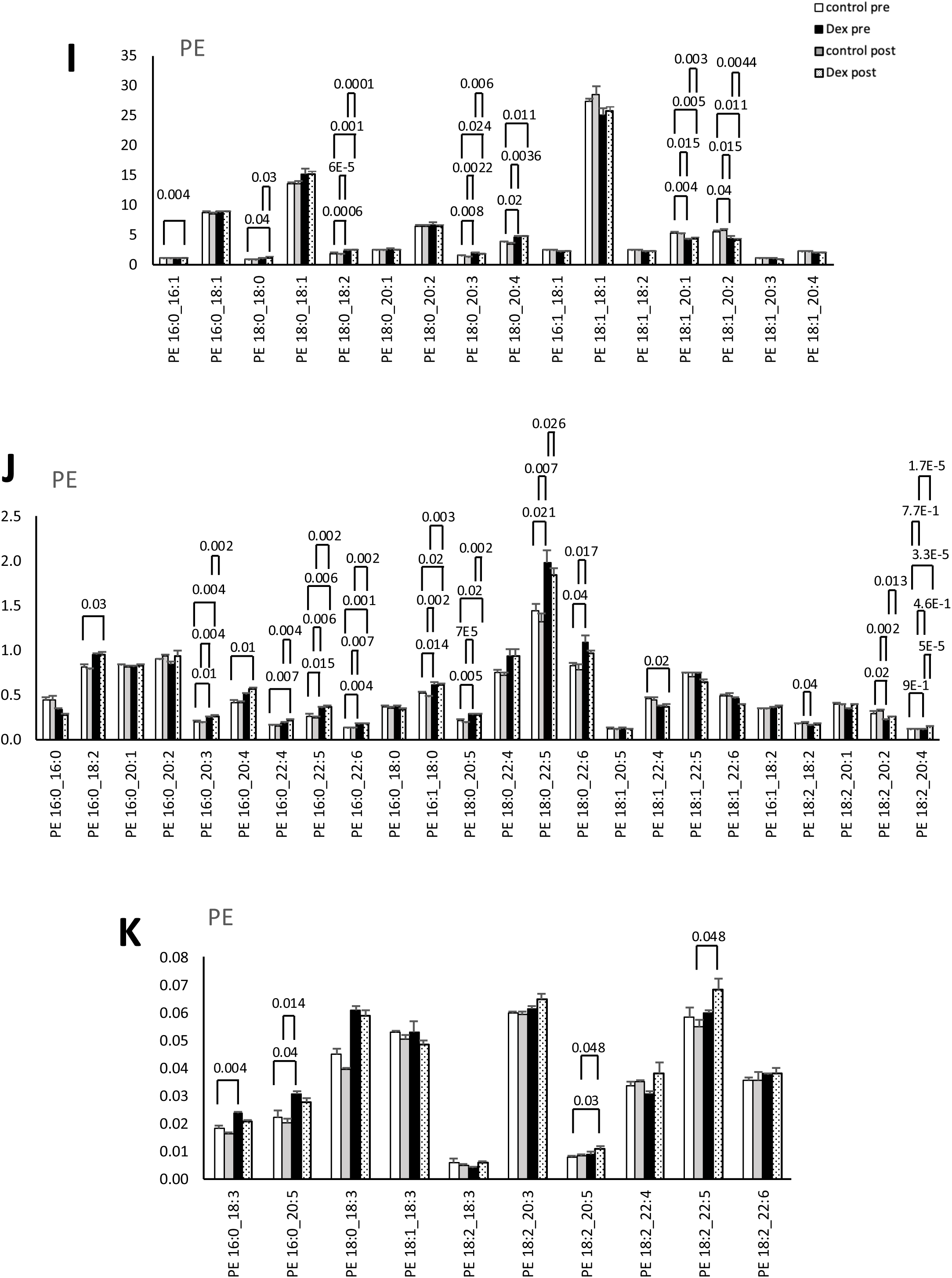
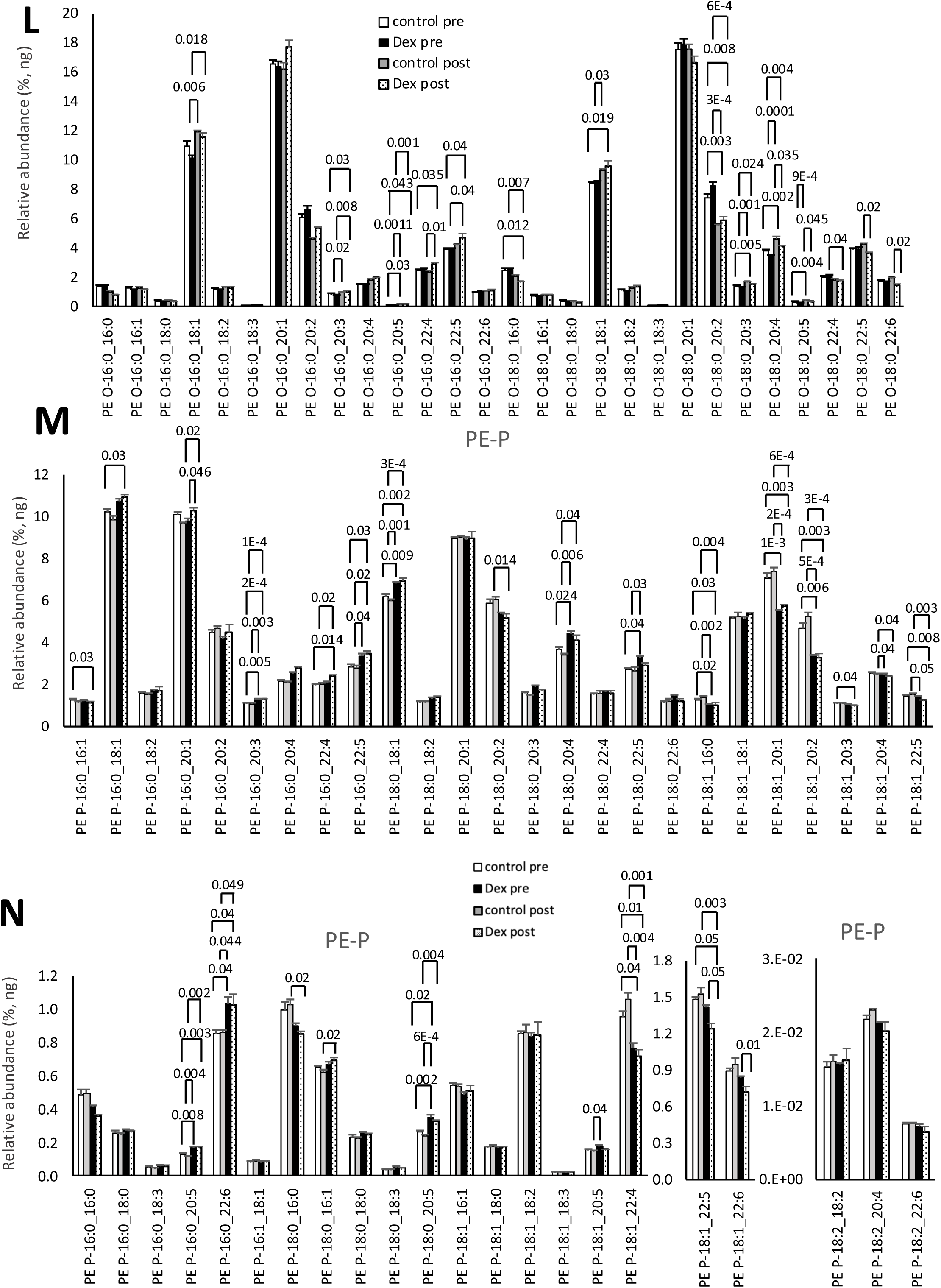
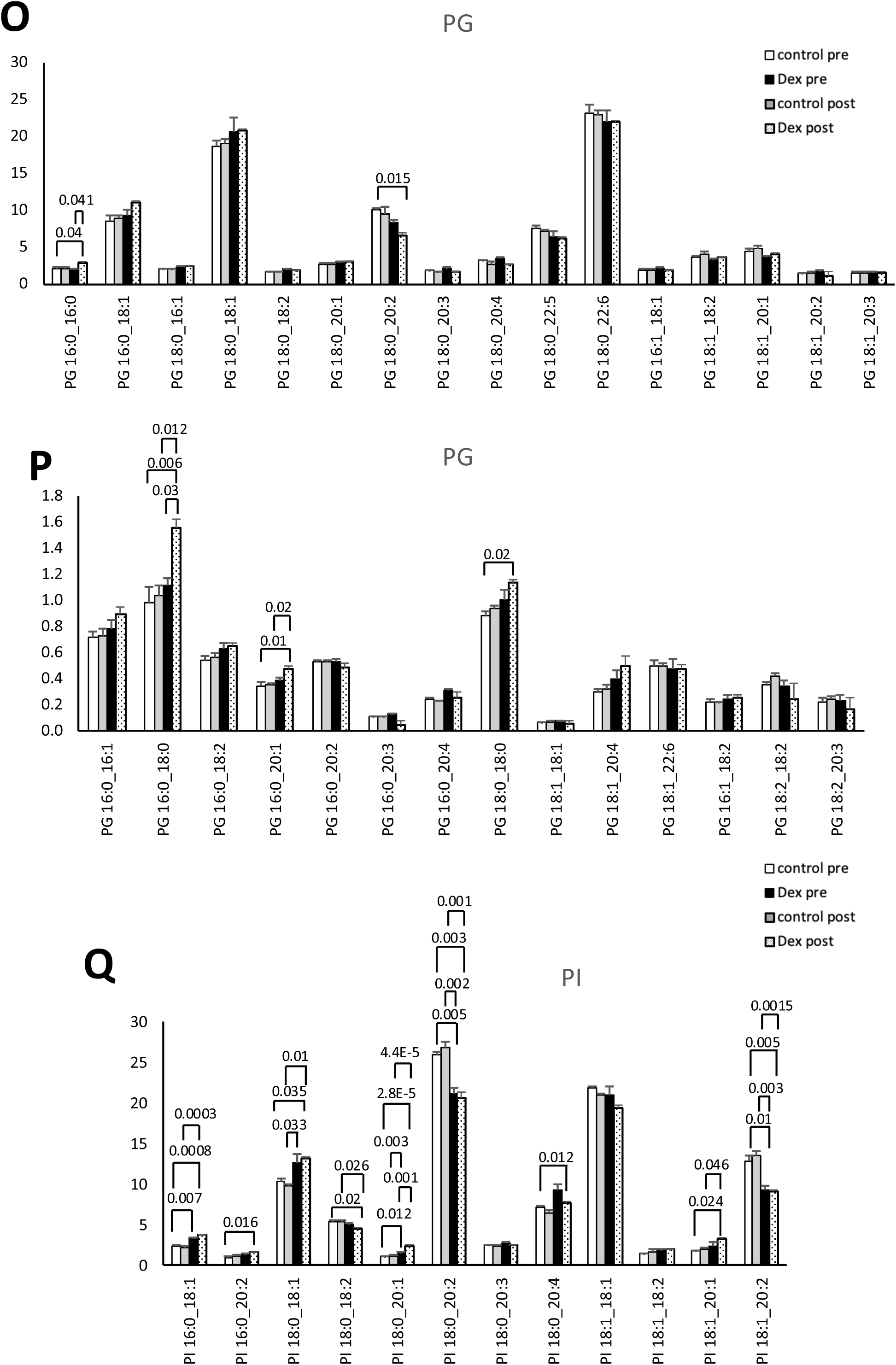
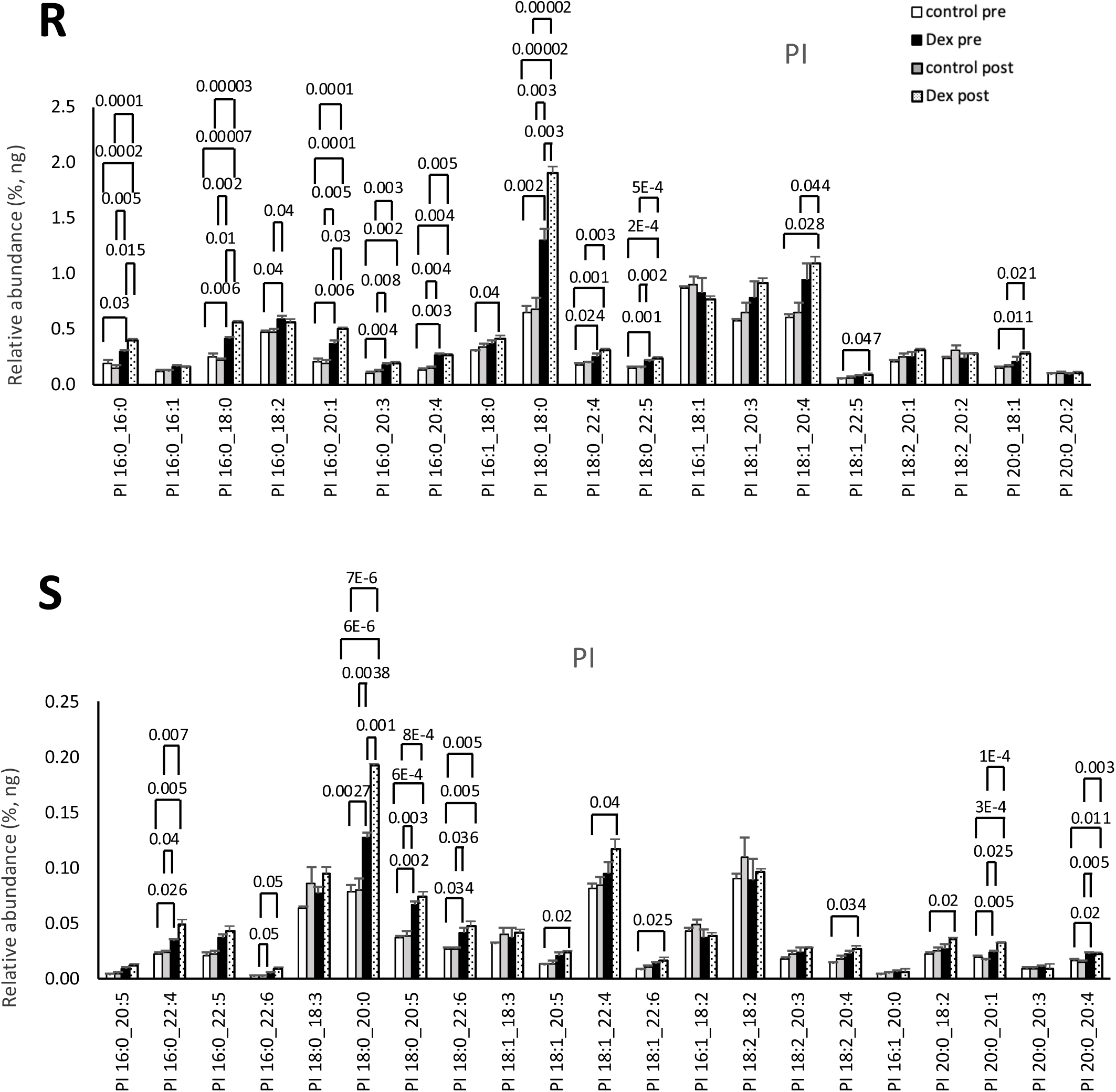
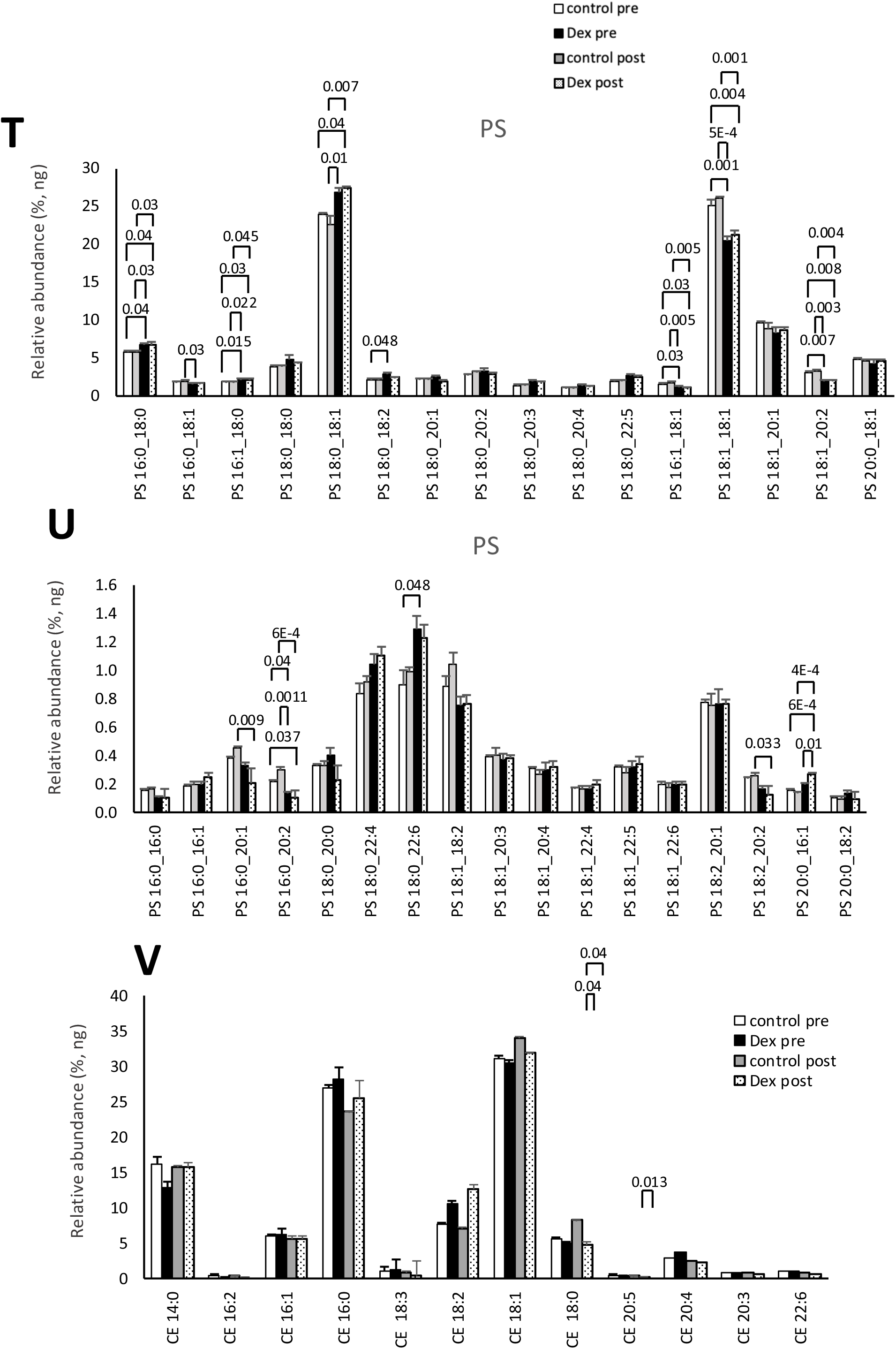
Dexamethasone treatment of host cells has no impact on SARS-CoV-2 membrane composition. *Panels A,B.* Lipids were extracted from virus from dexamethasone treated AAT cells then analyzed using LC/MS/MS as outlined in methods. Lipid amounts are expressed as ng recovered, or relative molar%, by totaling all species then normalizing to an average mass value per category (n=3 isolates, mean +/- SEM), unpaired Student t-test. *Panels B-V.* Lipids were extracted from virus from AAT cells treated with dexamethasone, then analyzed using LC/MS/MS as outlined in methods. Lipid amounts are expressed as relative ng% (n=3 isolates, mean +/- SEM), one-way Anova, Tukey post hoc test.

**Supplementary Figure 7.**
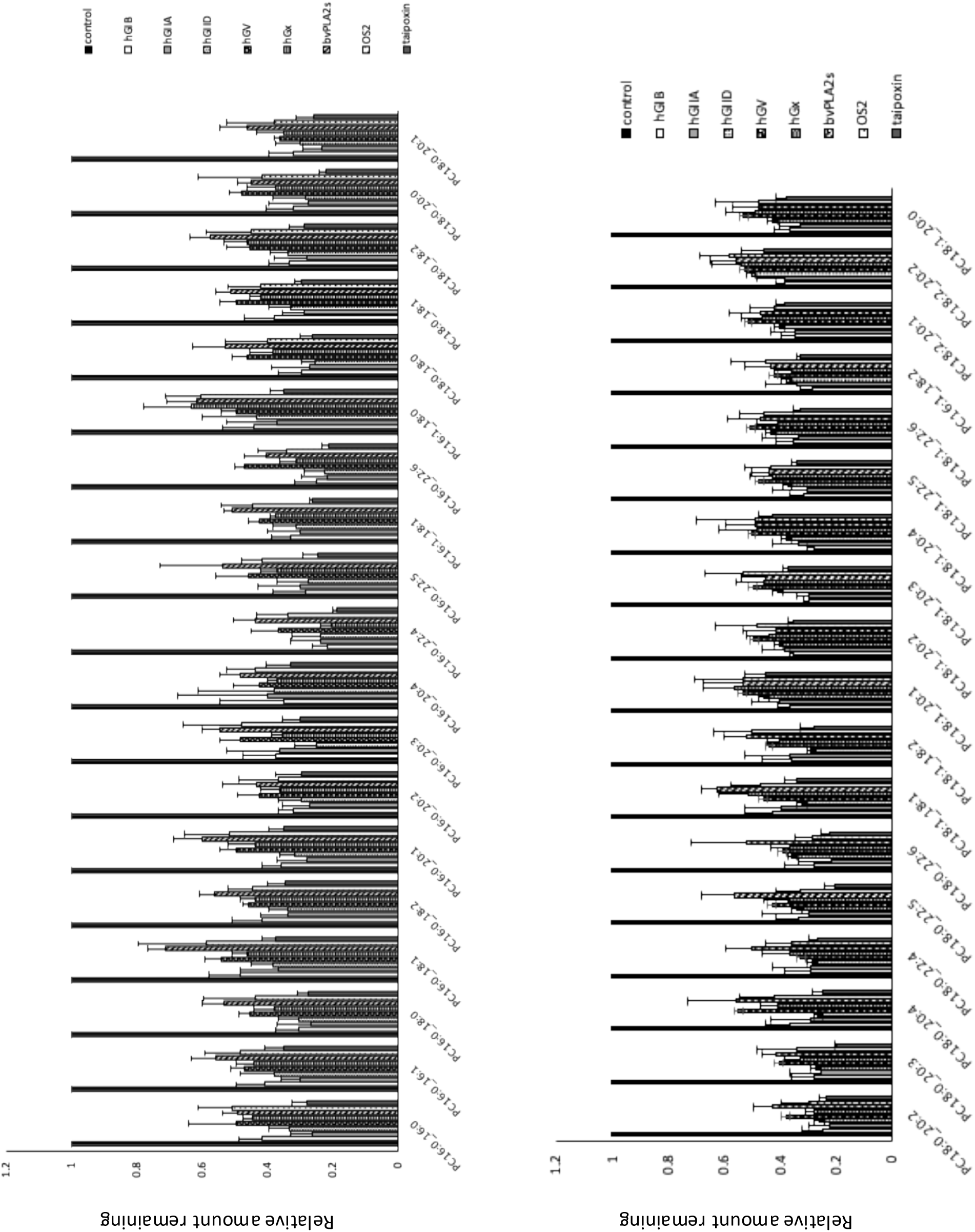
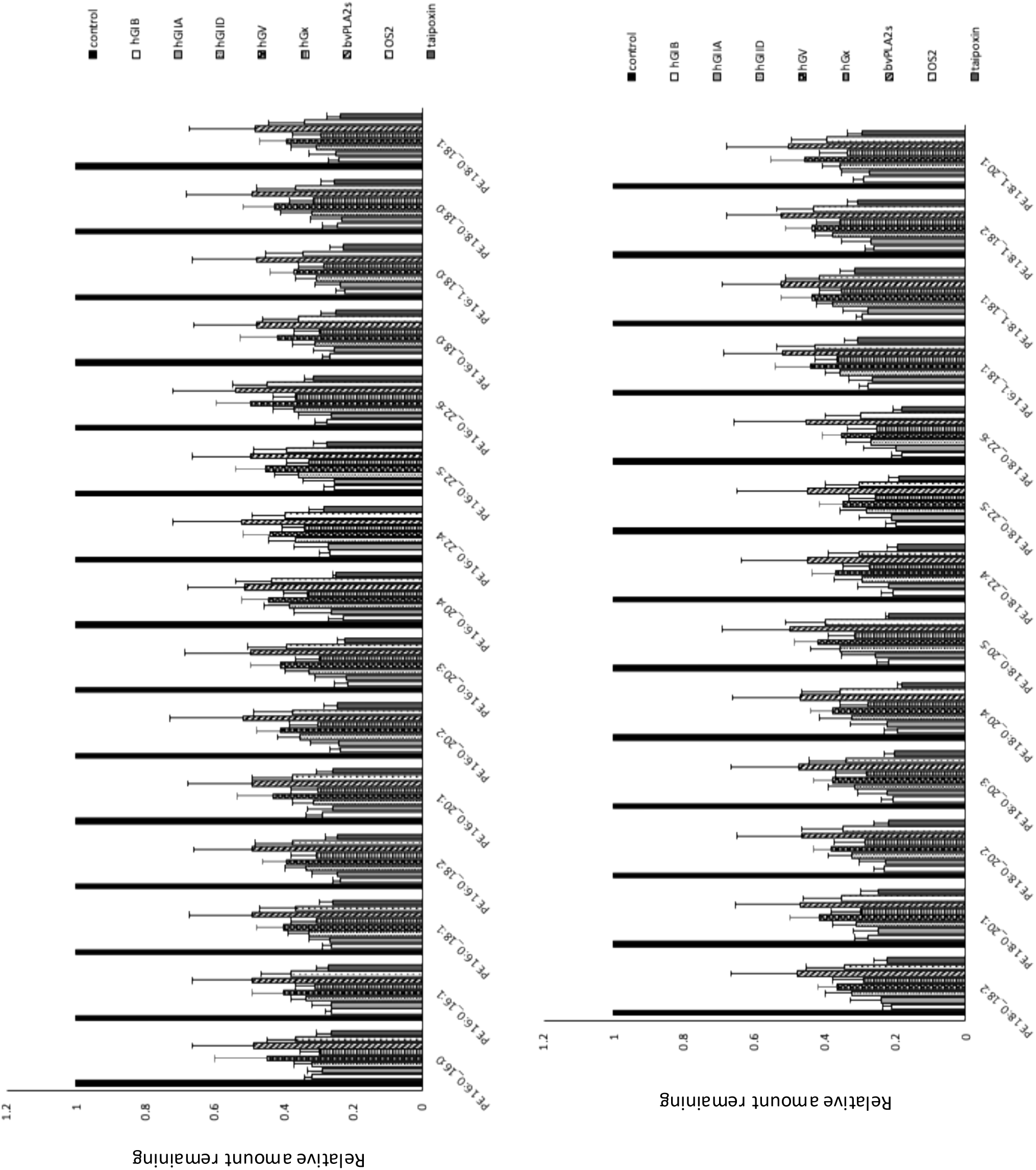
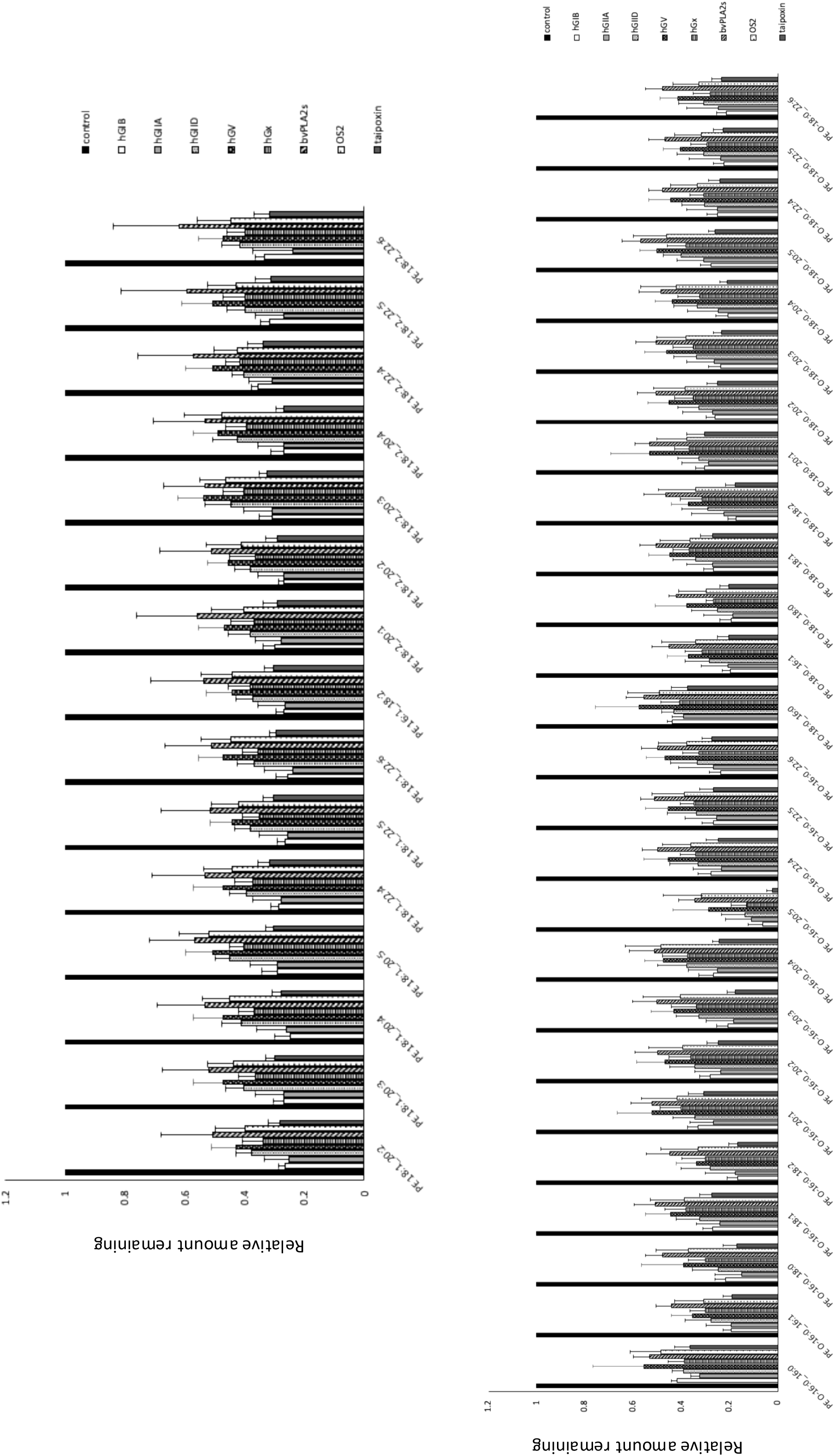
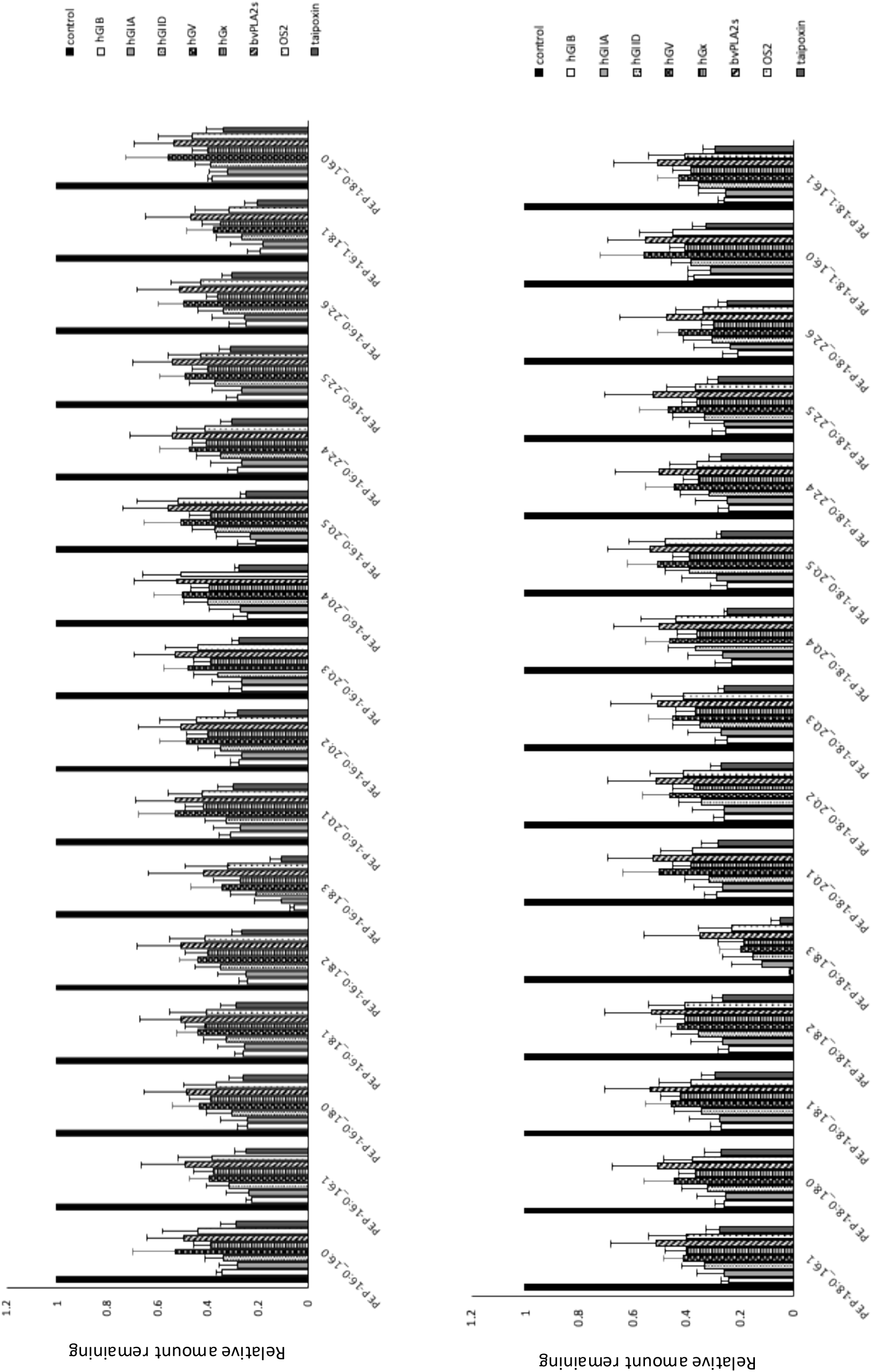
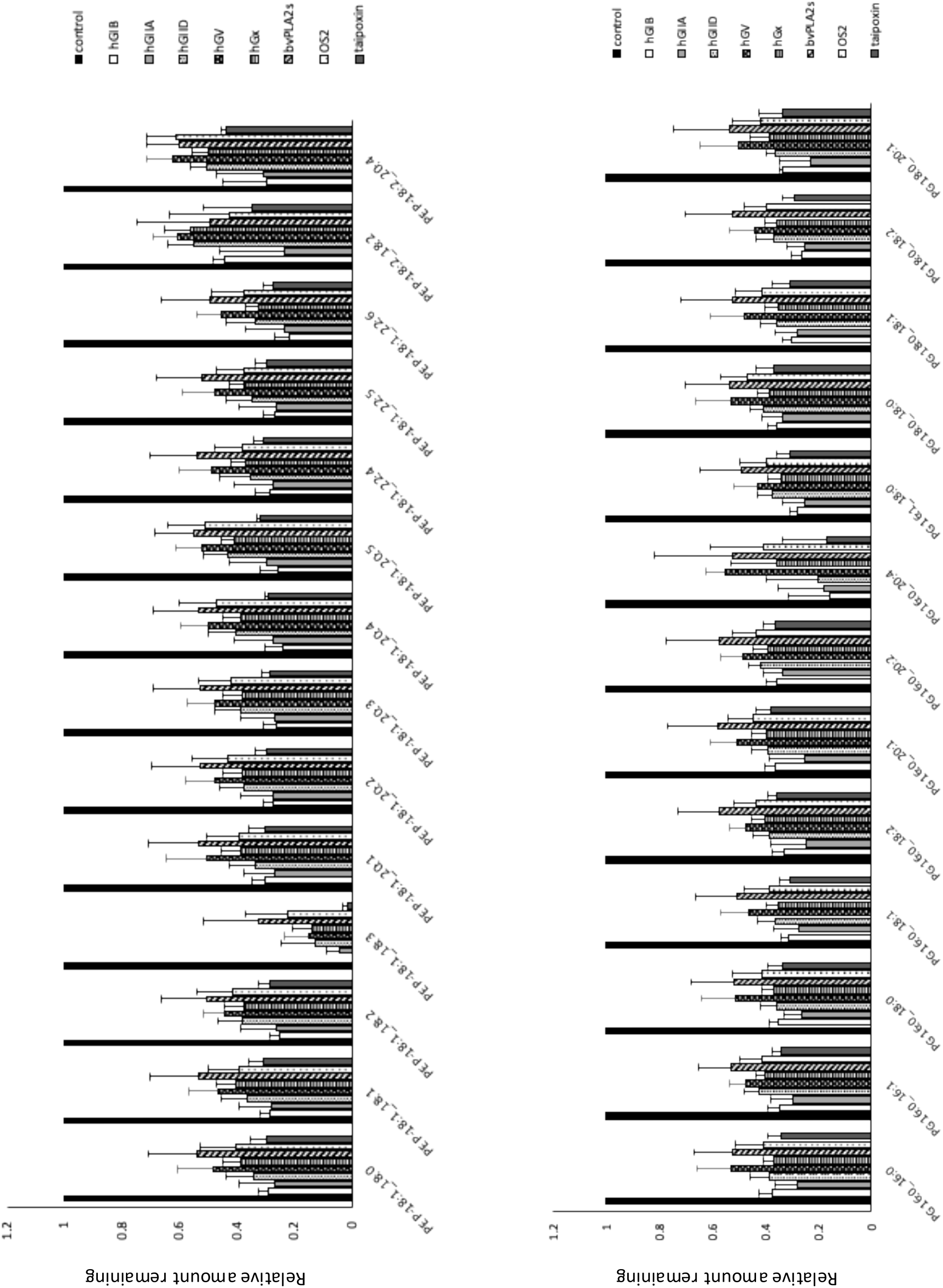
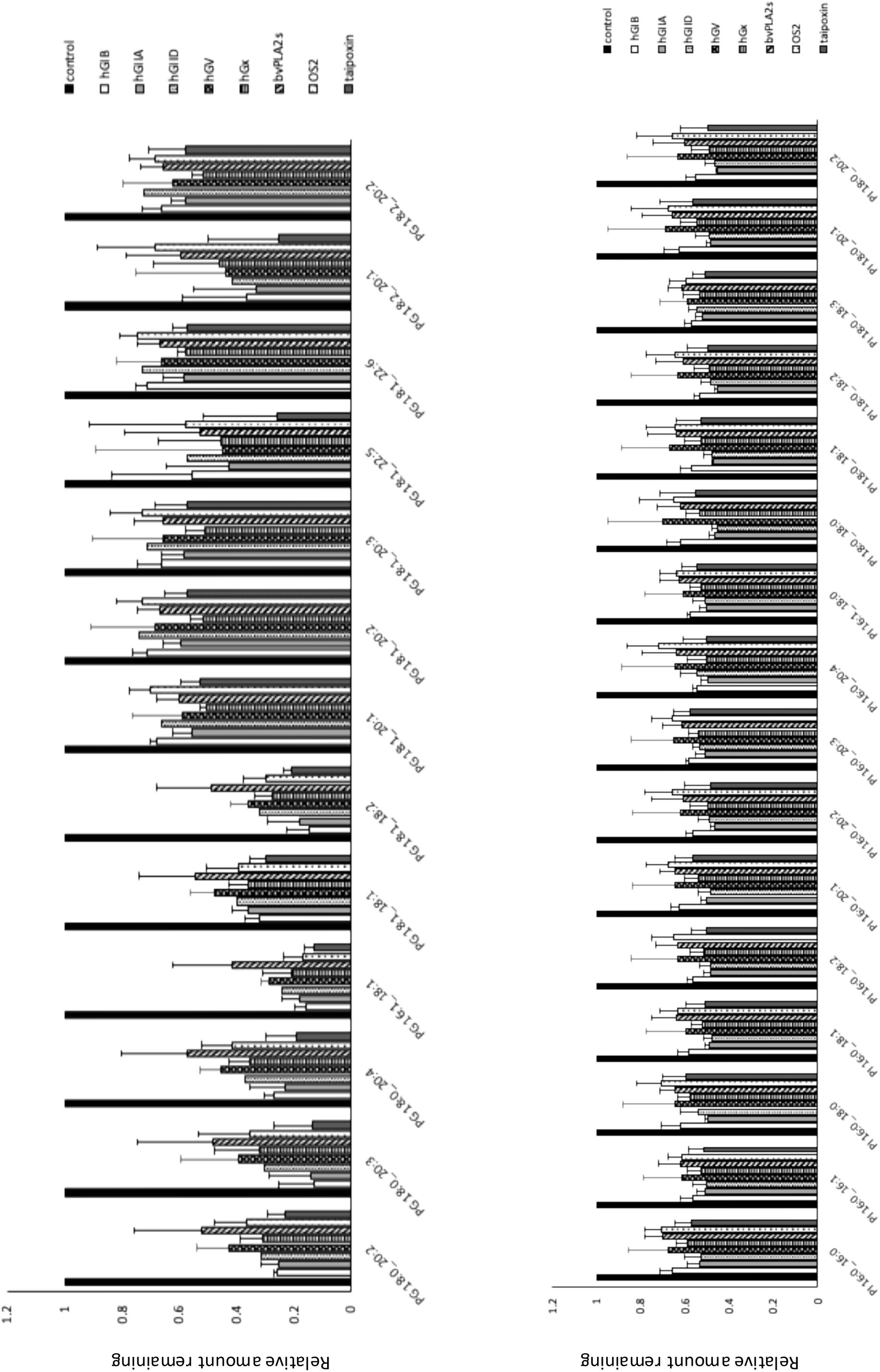
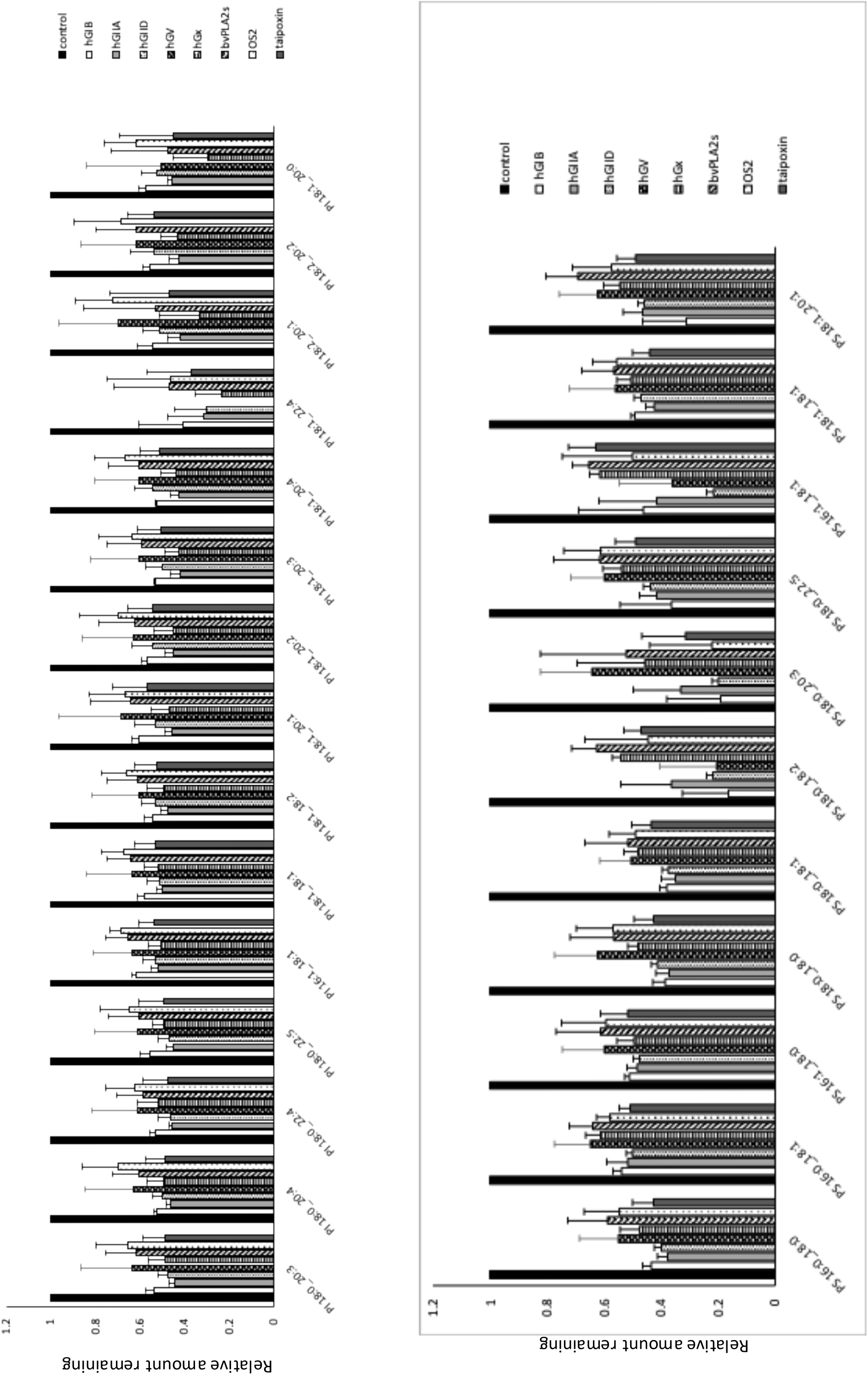
SARS-CoV-2 is sensitive to sPLA2 hydrolysis by several isoforms. Virus was exposed to several sPLA2 isoforms as outlined in Methods, then lipids extracted and analyzed using LC/MS/MS. The relative amounts of each molecular species remaining was determined, and totaled per lipid category, then normalized to untreated control virus PL levels (n=3 isolates, mean +/- SEM).

**Supplementary Figure 8.**
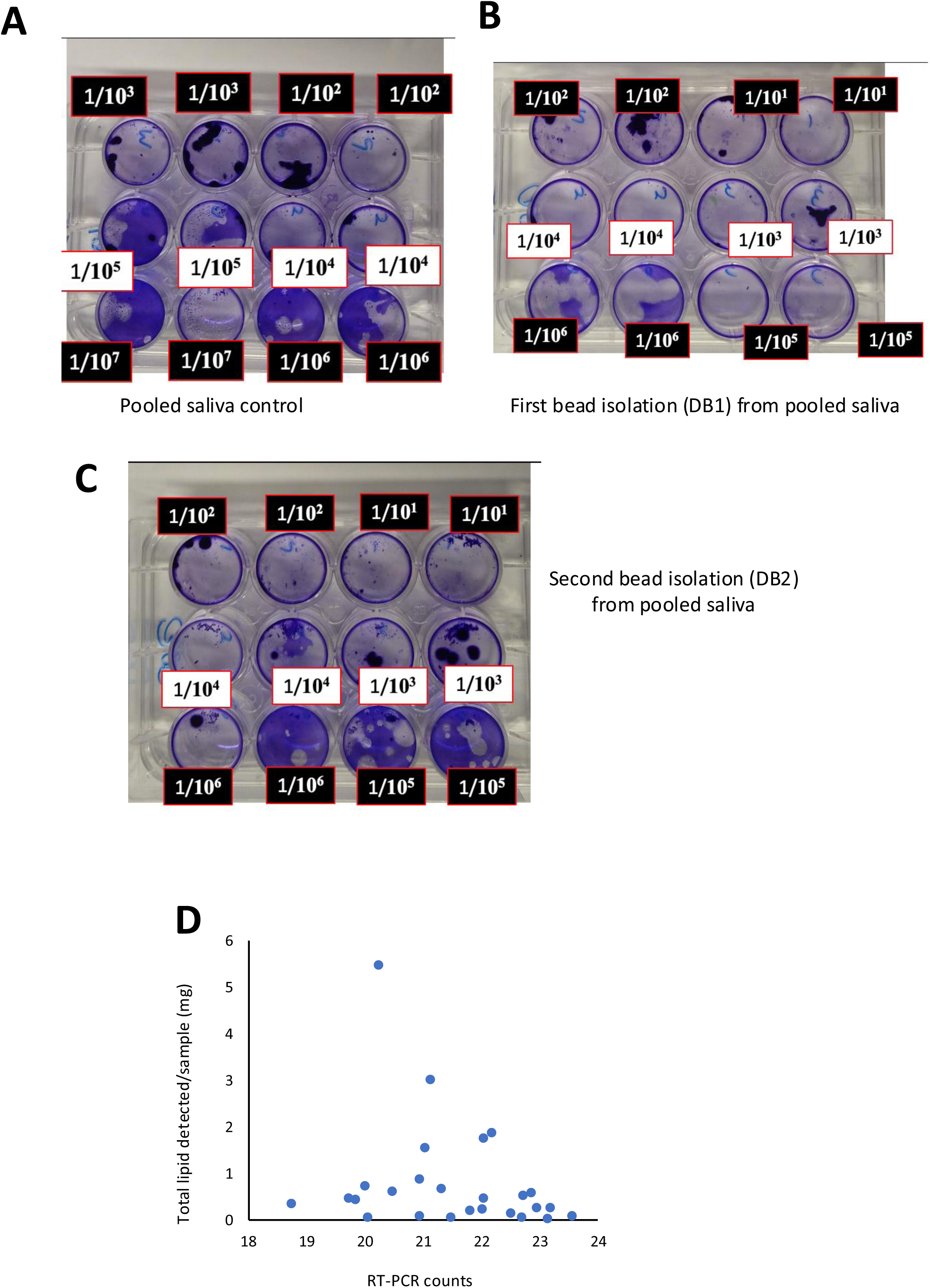
Plaque assays show the presence of live SARS-CoV-2 in pooled saliva before and after isolation using ACE2-coupled magnetic beads. *Panel A. Live virus in pooled sample of SARS-CoV-2*. Ten PCR positive clinical isolates were pooled and tested in a plaque assay at several indicated dilutions. *Panel B. Live virus is detected in the sample obtained after the first round of bead isolation*. The pooled sample was extracted using ACE2 coupled beads as outlined in Methods. The recovered sample was re-tested in a plaque assay at several indicated dilutions. *Panel C. Live virus is detected in the sample obtained after a second round of bead isolation*. The pooled sample was again extracted using ACE2 coupled beads as outlined in Methods, for a second time. The second recovered sample was re-tested in a plaque assay at several indicated dilutions. *Panel D. Total lipid measured doesn’t correlate with RT-PCR counts.* Clinical isolates (n=26) were extracted using ACE2-coupled beads as outlined in Methods, with both isolates per sample pooled to generate one isolate/sputum. Lipids were extracted and analyzed using LC/MS/MS as indicated in methods, and the total mg/sample compared with RC-qPCR counts obtained prior to virus isolation.

**Supplementary Figure 9.**
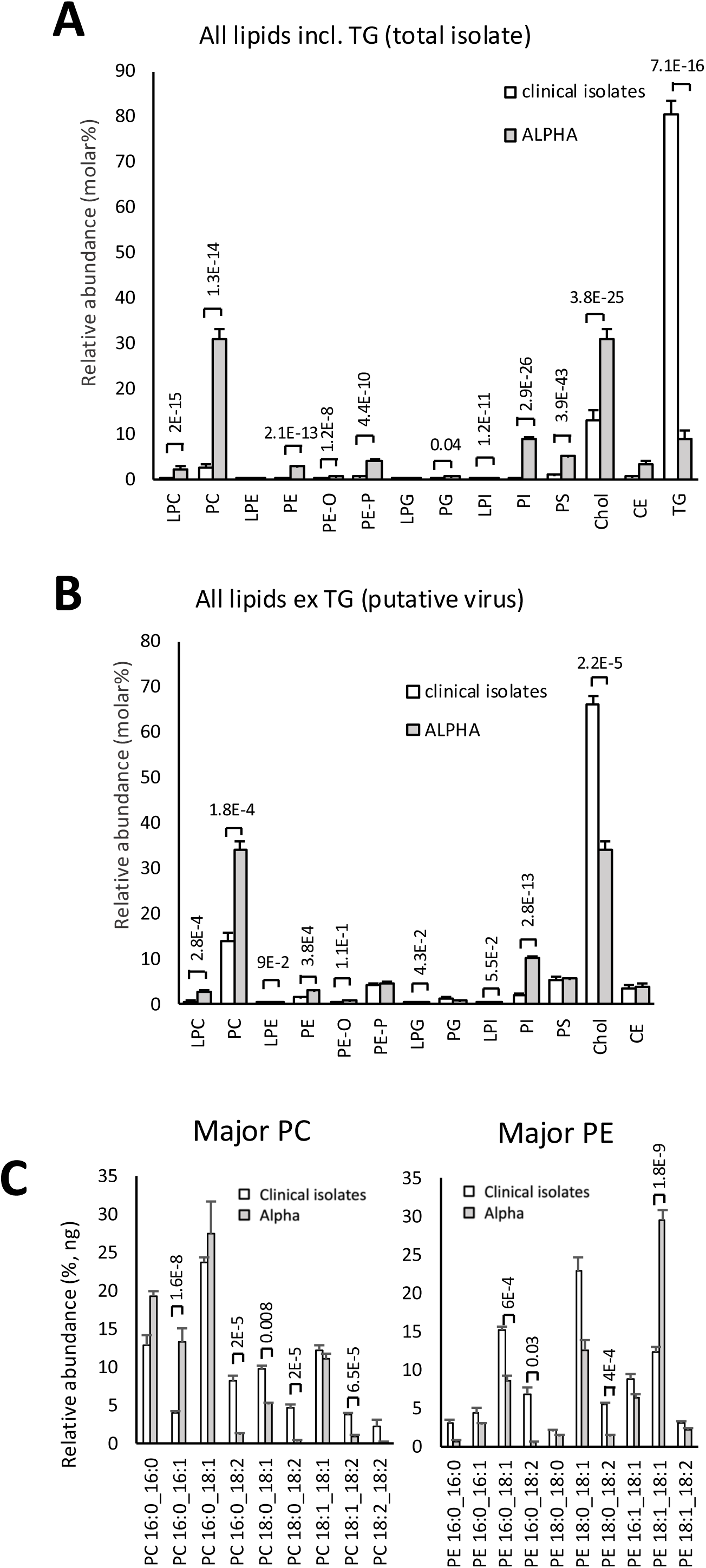
Lipid composition of recovered particles reveals major enrichment with TGs in clinical isolates, as compared with Alpha strain, but when excluding TG, particles become similar but show cholesterol enrichment. Lipids were extracted from clinical isolates or Alpha strain then analyzed using LC/MS/MS as outlined in methods. *Panel A. Lipid composition of extracted virus samples is highly TG enriched.* Lipid amounts (ng/sample) were converted to molar, totaled and expressed as a % total lipid using an average mass value per category (n=26 or 3, for clinical isolates and lab grown Alpha strain respectively, unpaired Student t-test). *Panel B. Lipid composition of recovered particles shows higher levels of cholesterol in clinical isolates, as compared with Alpha strain*. Lipid amounts (ng/sample), omitting TG, were converted to molar, totaled and expressed as a % total lipid using an average mass value per category (n=26 or 3, for clinical isolates and lab grown Alpha strain respectively, unpaired Student t-test). *Panels C,D. Within lipid categories some significant differences for PE and PC species are seen between clinical and lab grown SARS-CoV-2*. Lipids were converted to ng% within specific categories and compared (n=26, clinical, n=3 lab grown, mean +/- SEM, unpaired Student t-test).

**Supplementary Figure 10.**
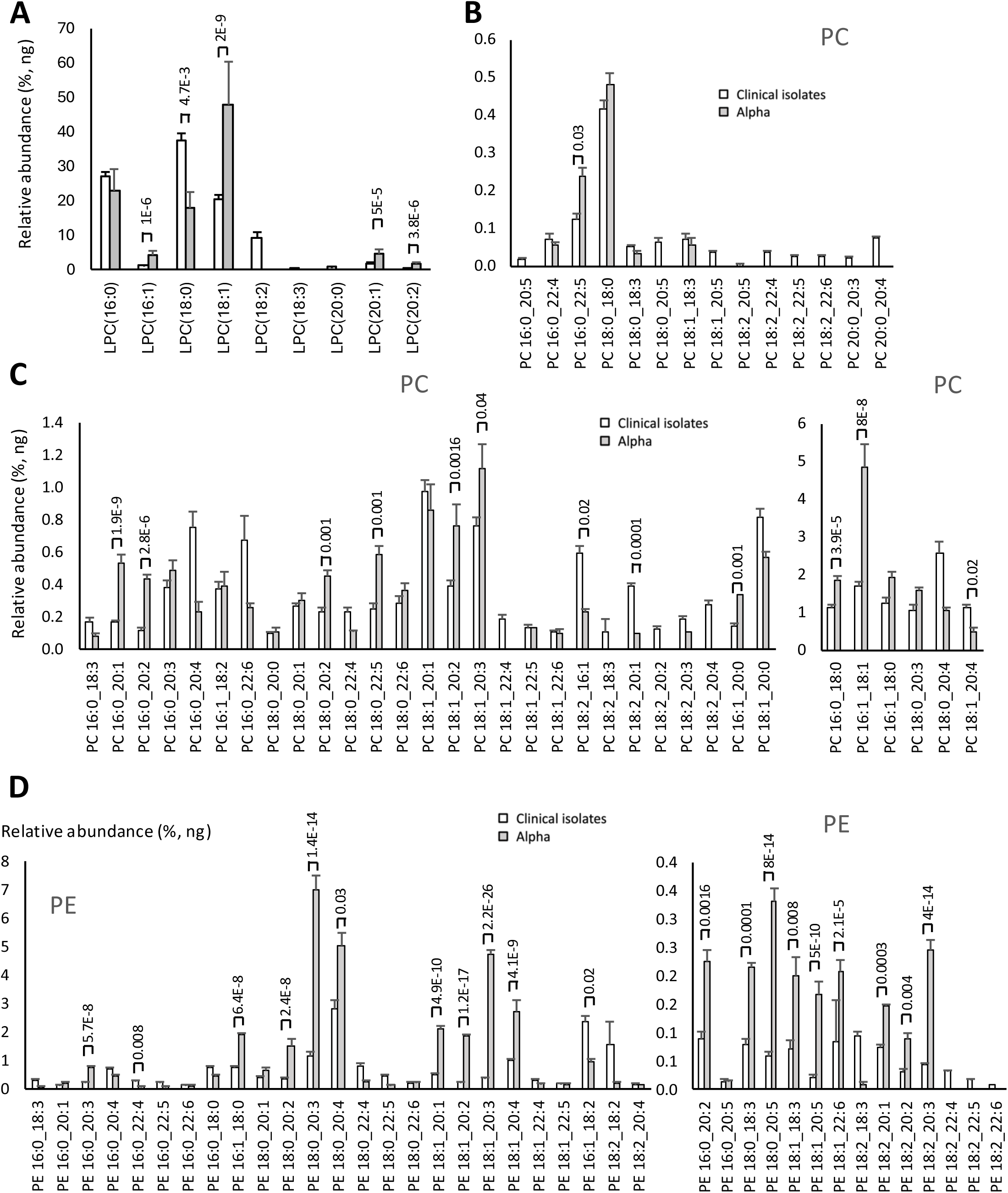
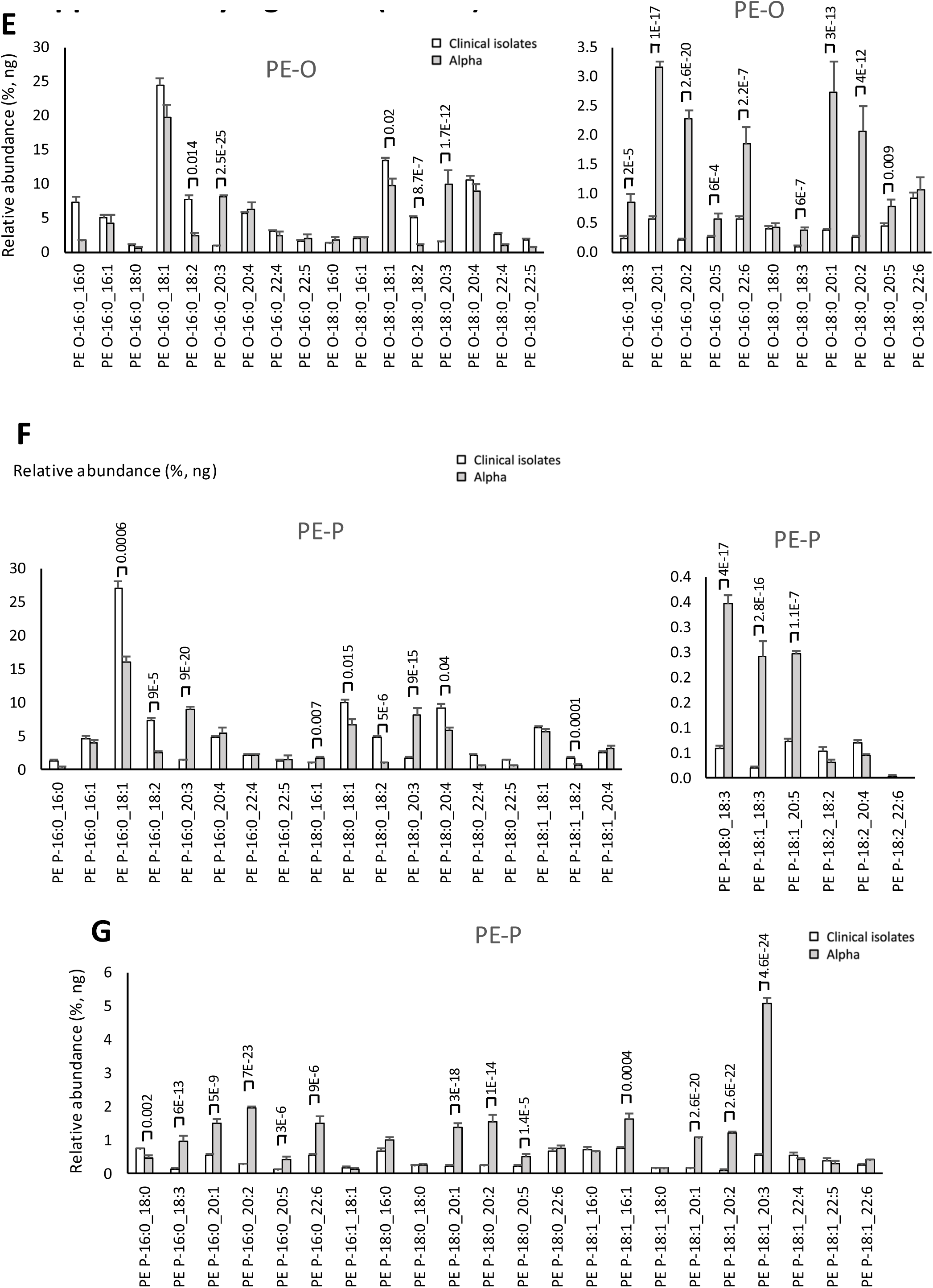
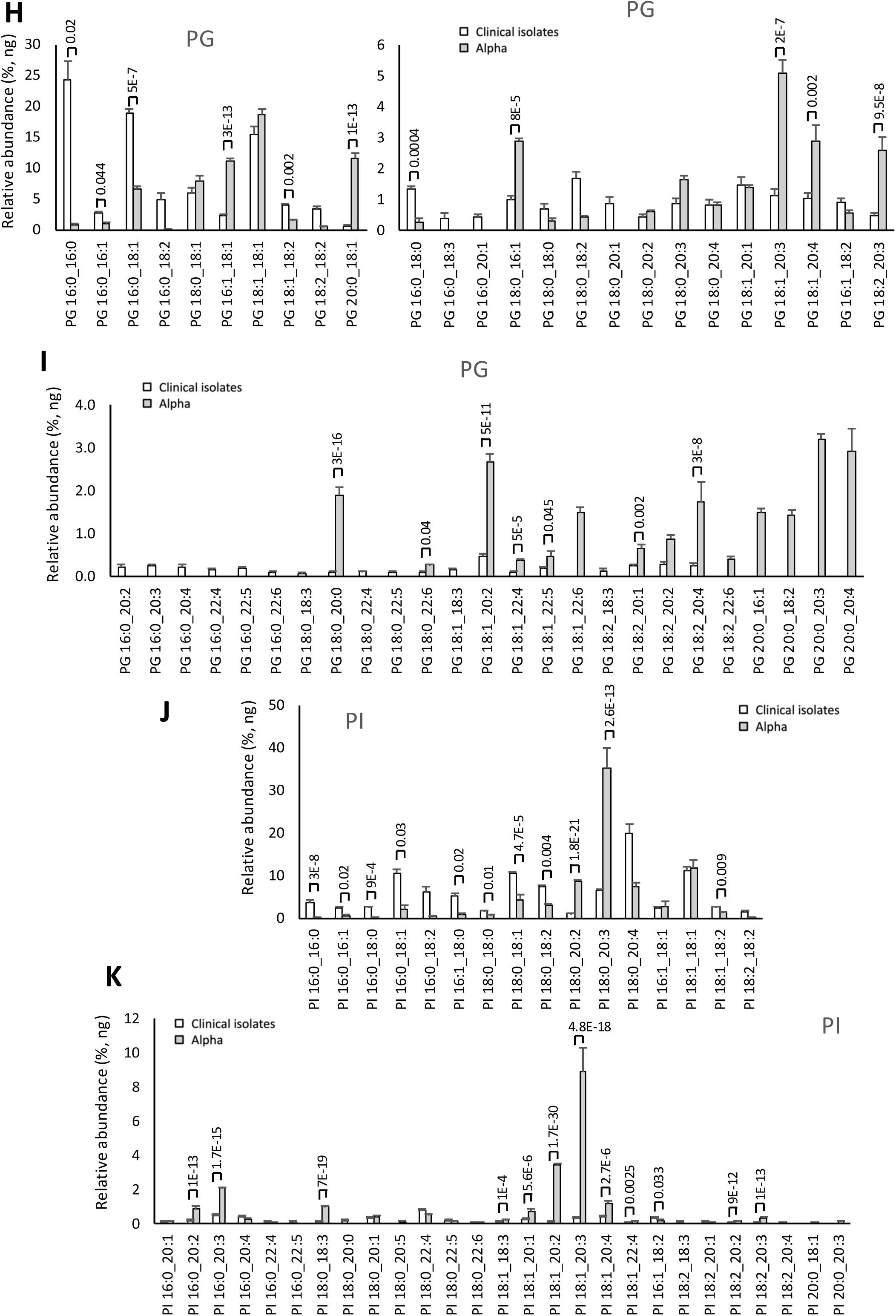
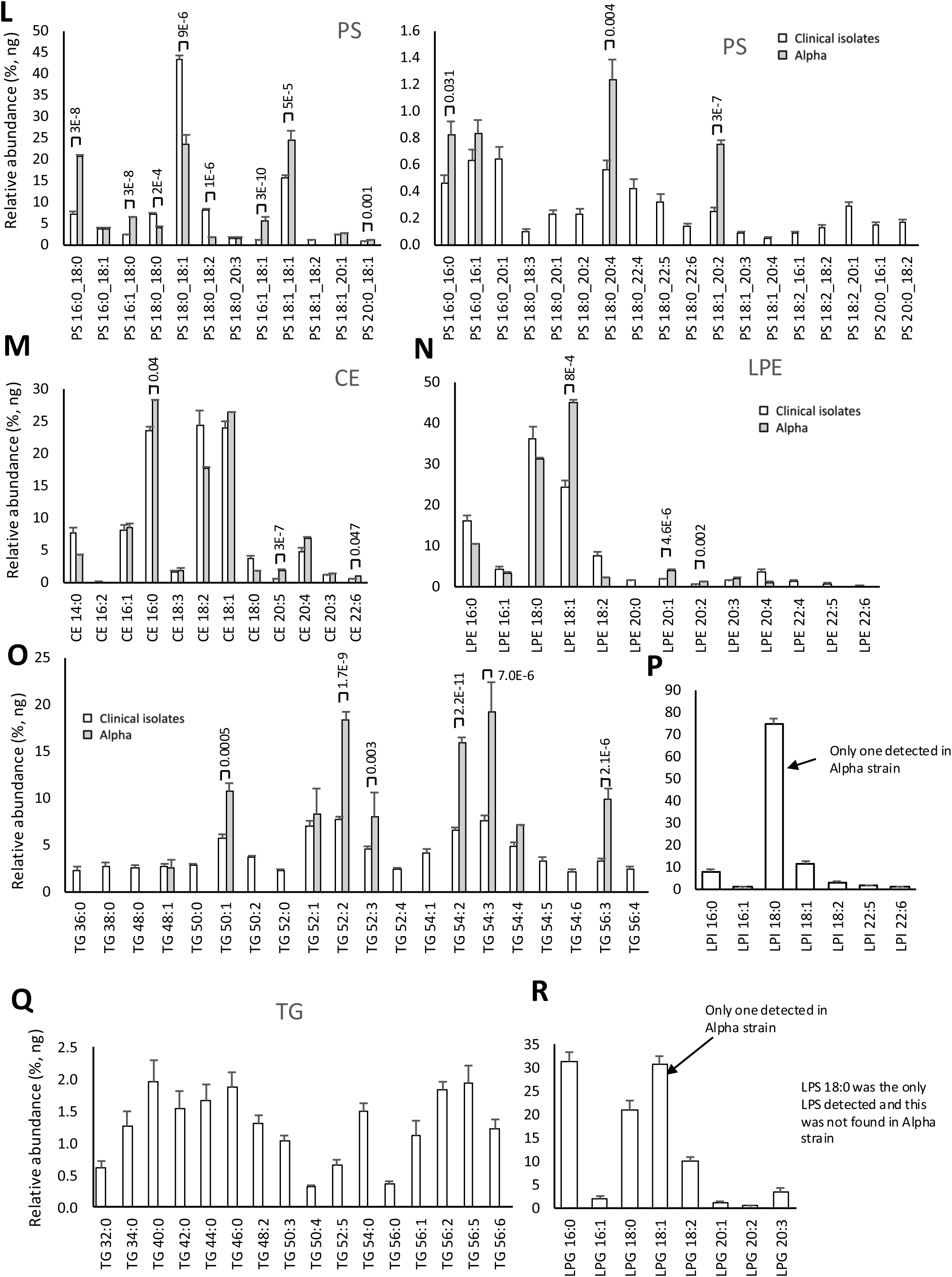
Comparison of clinical virus with Alpha strain reveals some differences in FA composition in individual PL categories. *Panels A-R.* Lipids were extracted from clinical isolates or Alpha strain then analyzed using LC/MS/MS as outlined in methods. Lipid amounts are expressed as relative abundance, ng% (n=26, clinical isolates, n=3 Alpha, mean +/-SEM), unpaired Student t-test.

**Supplementary Figure 11.**
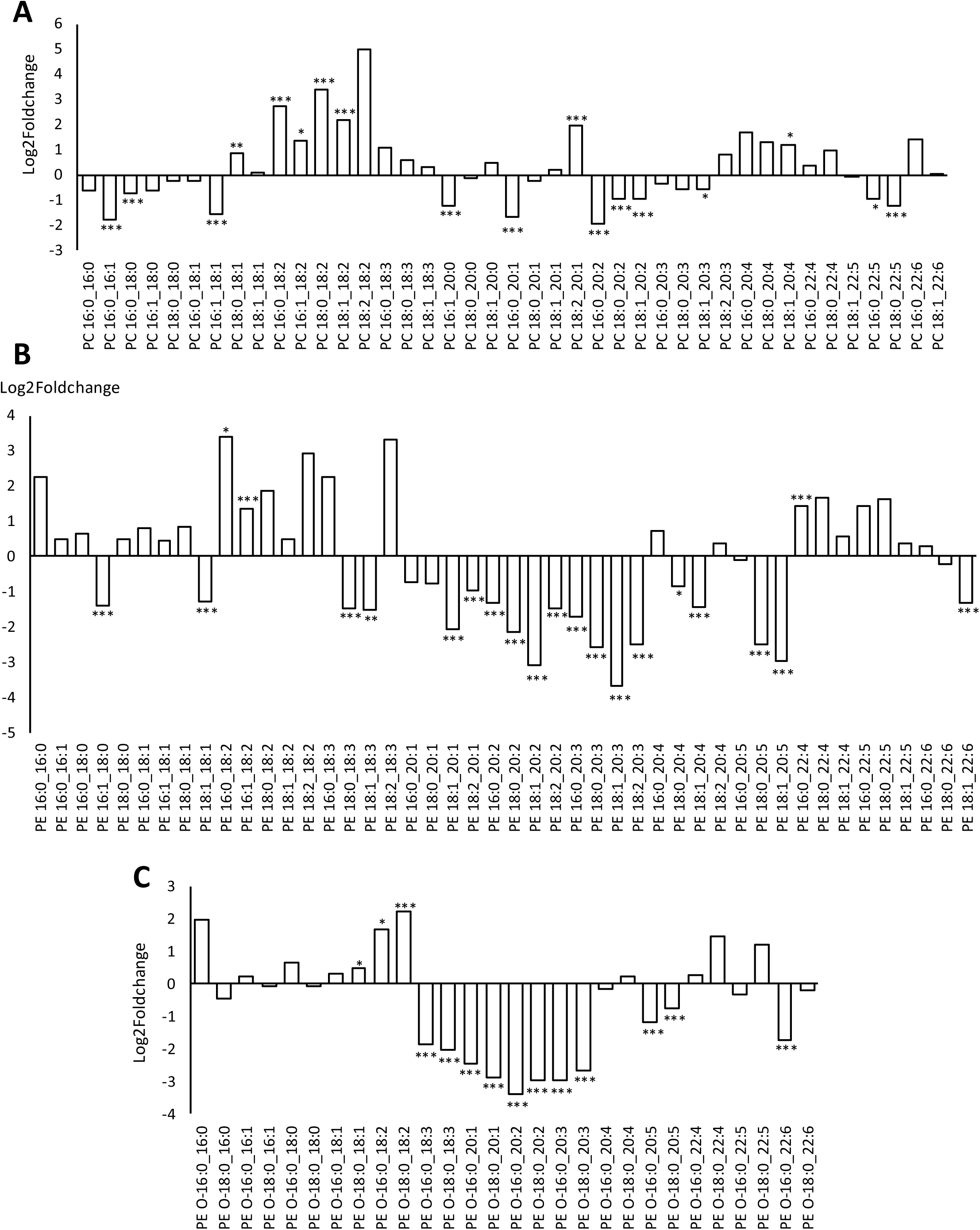
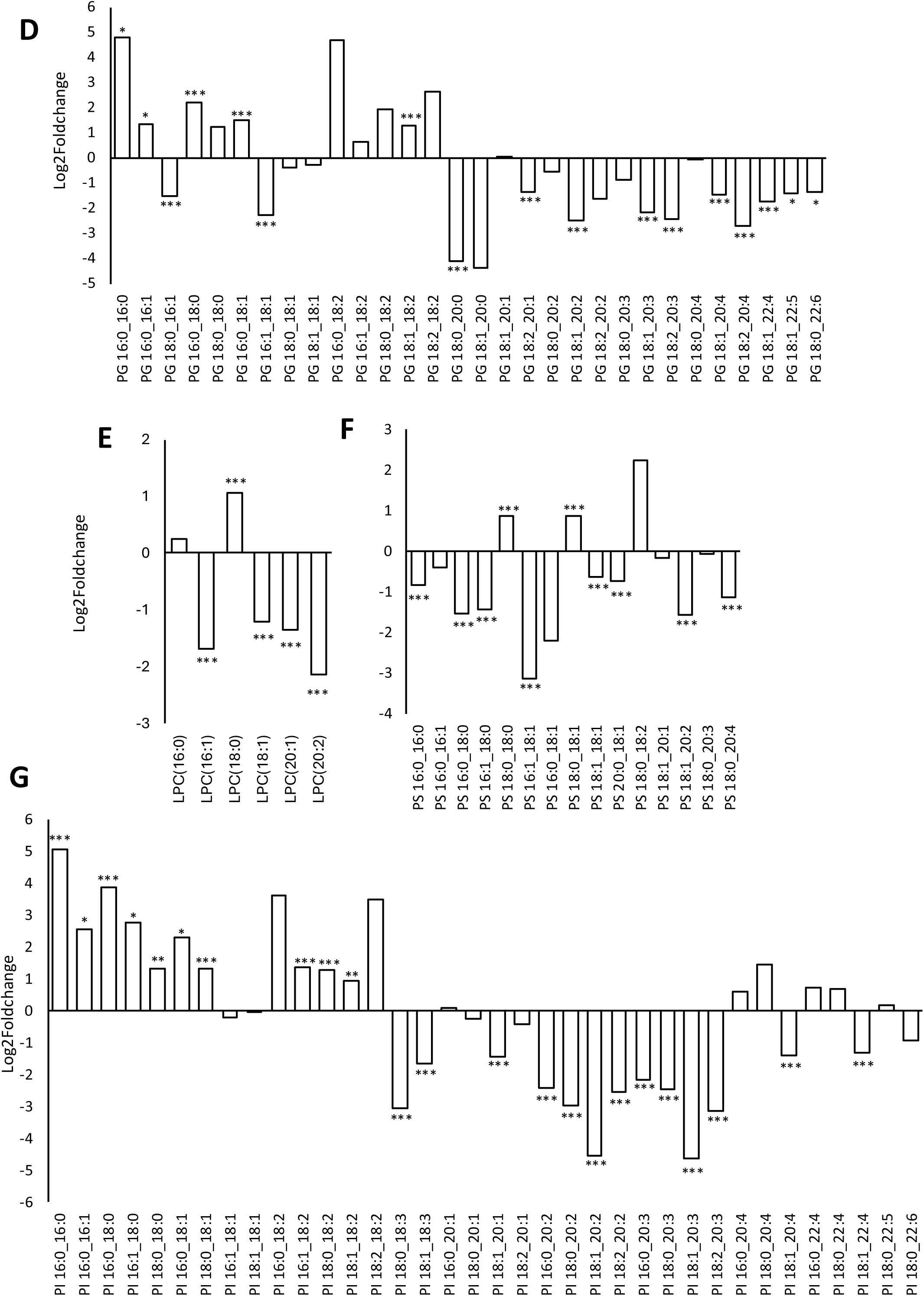
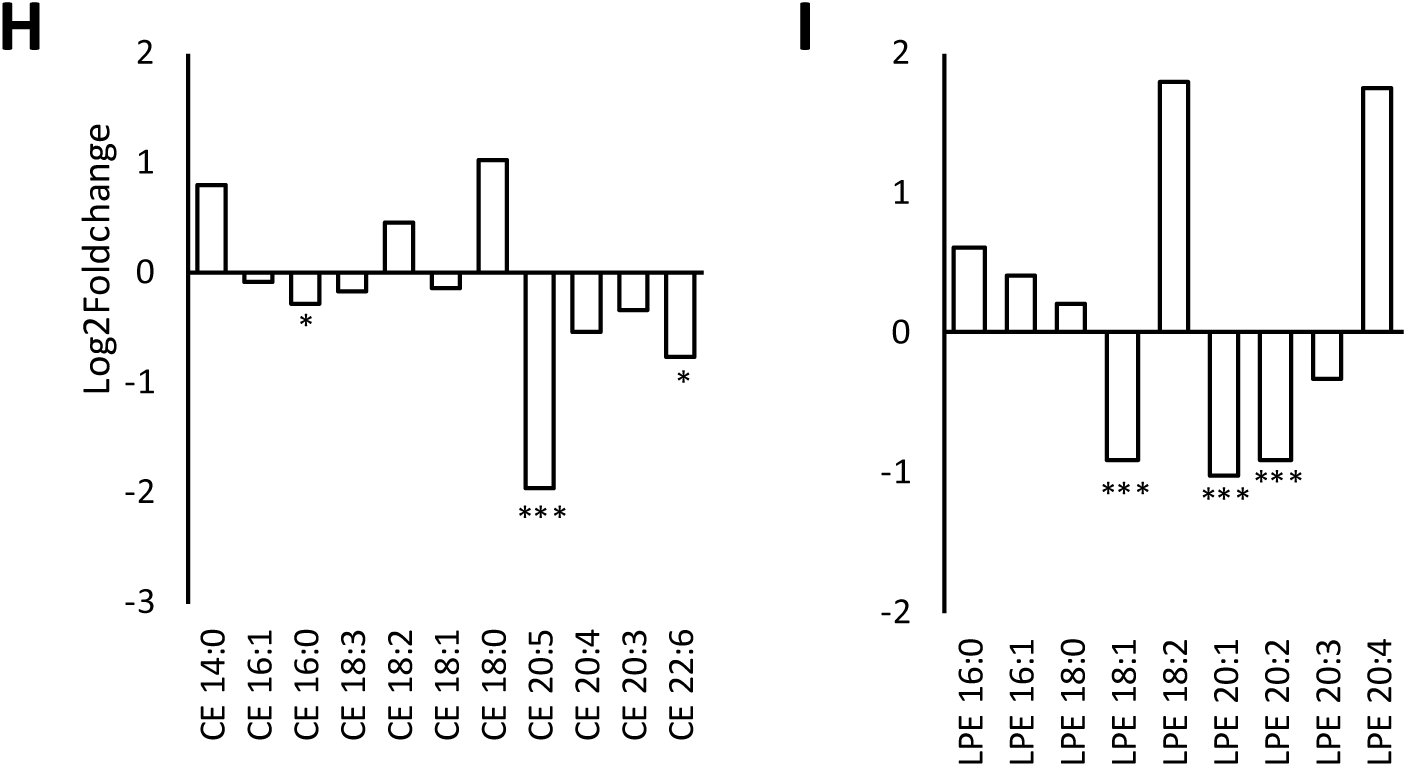
Comparison of clinical virus with Alpha strain reveals some differences in FA composition in individual PL categories. *Panels A-H.* Lipids were extracted from clinical isolates or Alpha strain then analyzed using LC/MS/MS as outlined in methods. Data from Supplementary Figure 10 is expressed as log2foldchange. A decrease means reduction in the species by IL-4, (n=26, clinical isolates, n=3 Alpha, mean +/- SEM), unpaired Student t-test, * P<0.05, ** P<0.01, ***P<0.005.

**Supplementary Figure 12.**
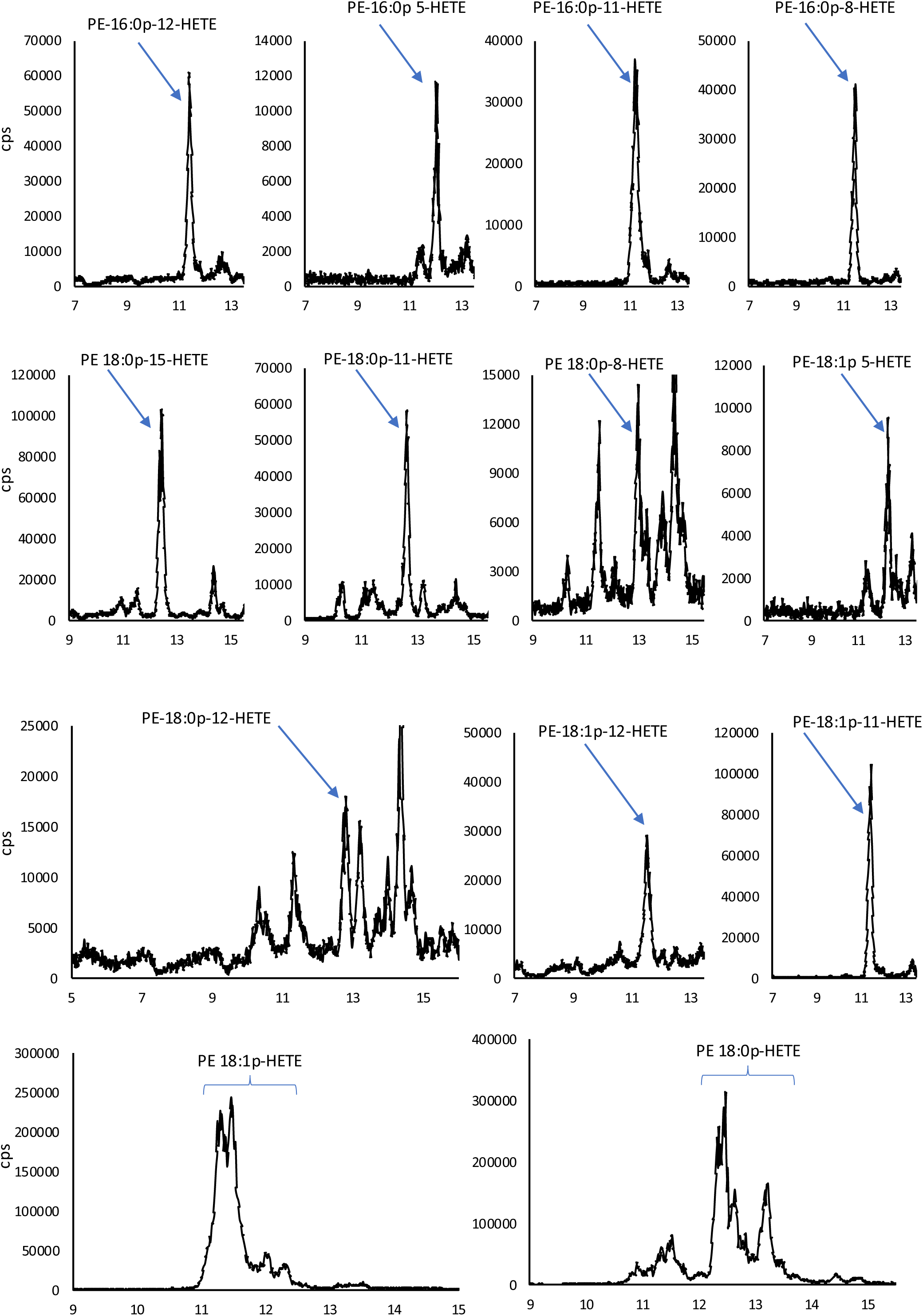
Representative chromatograms for oxPL measured in the study. Identification using internal daughter ions is based on coelution with the intact oxylipin fragment (m/z 319.2) as well as relative retention time as compared to standards (the mixed isomers of PE 18:0a_HETE with characteristic elution order).

**Supplementary Figure 13.**
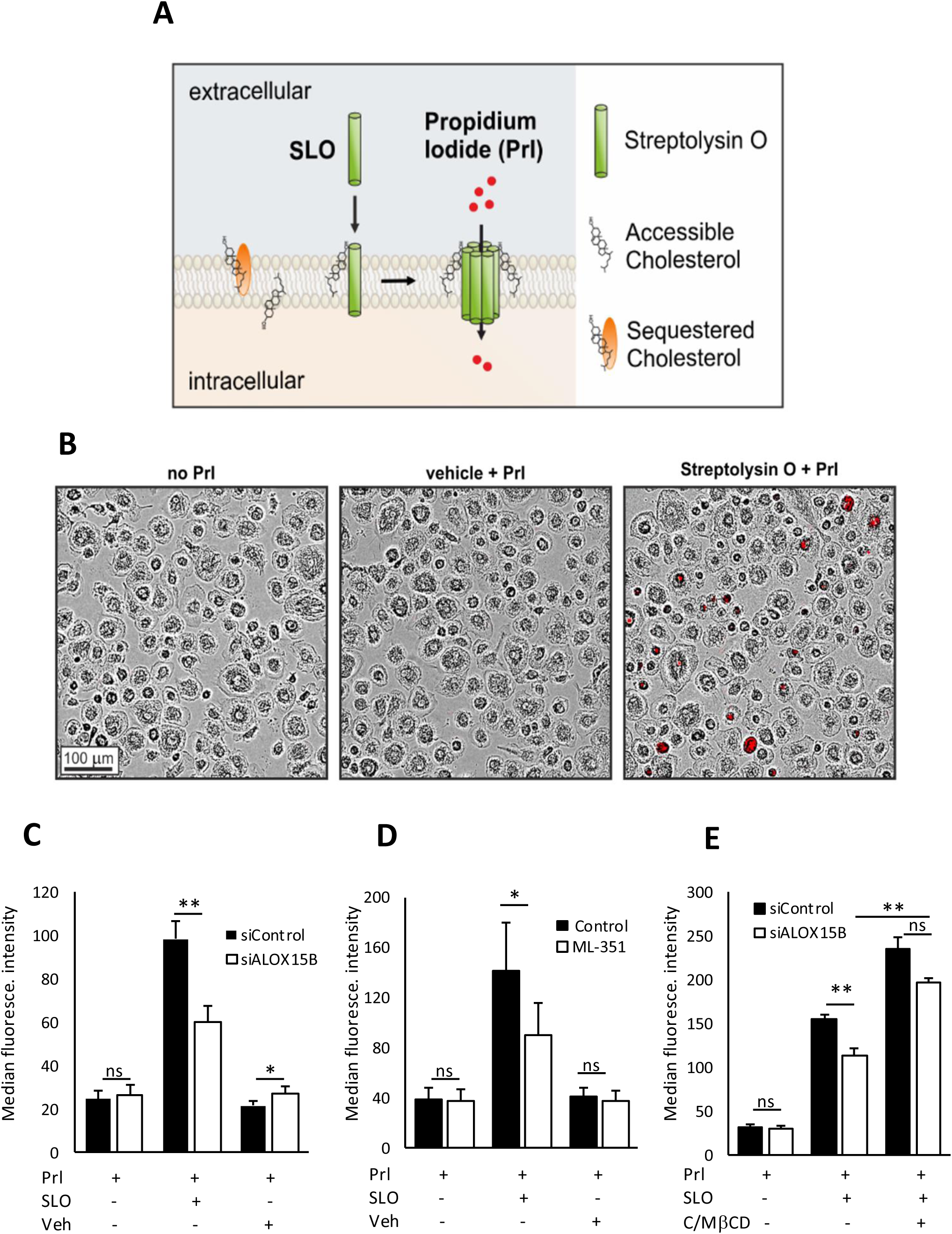
Accessible plasma membrane cholesterol is reduced in ALOX15B KD macrophages. *Panel A. Schematic illustration of streptolysin O (SLO)-mediated pore formation in plasma membrane and subsequent intracellular movement of propidium iodide (PrI). Panel B. Incucyte® live-cell imaging of SLO-mediated red PrI-fluorescence in SLO (29 nM) and PrI (1 µg/ml), but not vehicle (DTT, 20 µM) and PrI, treated macrophages. Panel C.* Median fluorescence intensity of PrI in SLO-treated control and ALOX15B KD macrophages. *Panel D.* Median fluorescence intensity of PrI in SLO-treated control and ML-351 pretreated macrophages. *Panel E.* Median fluorescence intensity of PrI in SLO-treated control and ALOX15B KD macrophages pretreated with cholesterol-loaded methyl-β-cyclodextrin (10 µg/ml). (mean ± SE from at least four independent experiments. Statistical analysis was performed using two-tailed student’s t-test for C and E (*P<0.05 and **P<0.01 vs siControl) and D (*P<0.05 and **P<0.01 vs control).

